# Glycosylation of anandamide and other bioactive *N*-acylethanolamines in mammalian cells and tissues

**DOI:** 10.64898/2026.07.16.738921

**Authors:** Anna F. Stevens, Remco E. A. Peter, Berend Gagestein, Maria J. Ferraz, Eline Been, Tijn Vleeshouwer, Iakovia Ttofi, Richard J.B.H.N van den Berg, Tom van der Wel, Laura V. de Paus, Christian Deuschle, Cas van der Horst, Laura H. Heitman, Marta Artola, Daniele Piomelli, María Teresa Grande, Julian Romero, Hermen S. Overkleeft, Kathrin Brockmann, Thomas Gasser, Johannes M. F. G. Aerts, Mario van der Stelt

## Abstract

*N*-acylethanolamines (NAEs), including the endocannabinoid anandamide, are bioactive fatty acid amides that are normally hydrolyzed by fatty acid amide hydrolase (FAAH) or *N*-acyl acid amidohydrolase (NAAA). Strikingly, when canonical NAE degradation is blocked, NAE levels do not increase indefinitely but instead reach a plateau. This apparent metabolic ceiling suggests that additional, underexplored pathways contribute to NAE homeostasis. Identifying these pathways is essential to determine whether NAEs are converted into inactive metabolites or products with distinct biological properties.

Here, we identify NAE glycosylation as a metabolic pathway that links endocannabinoid-related lipid metabolism to glycosphingolipid turnover. We synthesized glycosylated NAEs and their isotope-encoded standards and developed targeted LC–MS/MS assays to monitor their enzymatic processing and quantify their abundance in mouse and human cells, tissues, and plasma. We show that non-lysosomal glucosylceramidase GBA2 transfers glucose or galactose to anandamide, *N*-oleoylethanolamine and *N*-palmitoylethanolamine, and lysosomal glucosylceramidase GCase hydrolyses β-Glycosylated-NAEs (β-Glyco-NAE) back to their parent NAEs. β-Glyco-NAEs occur endogenously in macrophages and neuronal cells, increase when canonical NAE degradation is impaired, and accumulate in human samples with GCase deficiency, including Gaucher disease and *GBA1*-associated Parkinson’s disease. β-Glyco-NAEs do not engage the cannabinoid receptors, TRPV1, or PPARα, and potentiate inflammatory cytokine release, including IL6 and TNFα, from microglia. Based on these findings, we pose that GBA2-dependent NAE glycosylation may constitute an overflow lipid-remodeling pathway that connects NAE metabolism to lysosomal dysfunction, inflammation and neurodegeneration.

## Introduction

Lipids are not merely structural components of biological membranes and energy storage molecules. They are also dynamic signaling metabolites that convey information on cellular state, stress and disease. *N*-acylethanolamines (NAEs) form a family of bioactive fatty acid amides in which an ethanolamine head group is bound to a fatty acyl chain through an amide bond. Variation in acyl chain length and unsaturation generates structurally distinct NAEs with partially overlapping, biologically diverse functions. NAEs regulate neurotransmission, inflammation, energy balance, appetite, stress responses and anxiety through a broad repertoire of molecular targets, including the cannabinoid CB_1_ and CB_2_ receptors, the GPR55, GPR110, GPR119 and TRPV1 ion channels and the nuclear receptors PPARα and PPARγ.^1,2^

The best-studied NAEs are *N*-arachidonoylethanolamine (AEA; anandamide), *N*-palmitoylethanolamine (PEA) and *N*-oleoylethanolamine (OEA). The endocannabinoid, AEA is an endogenous cannabinoid CB_1_ and CB_2_ receptor (CBR) ligand that also activates TRPV1, thereby linking NAE signaling to synaptic transmission, motor control, emotional behavior, pain, memory and stress adaptation.^3,4^ PEA and OEA do not signal through cannabinoid receptors, but engage PPARα, GPR55 and/or GPR119.^5,6^ Through these targets, PEA exerts anti-inflammatory, analgesic and neuroprotective effects, whereas OEA acts as a satiety signal that reduces food intake and contributes to metabolic regulation.^7^ Together, these molecules illustrate how subtle changes in lipid structure impact signaling output across neuronal, immune and metabolic systems.

Dysregulated NAE signaling has been implicated in several pathological conditions, such as neuroinflammation, metabolic disease and neurodegeneration, which can result in ischemic infarction, obesity and Parkinson’s disease (PD).^8–11^ Changes in NAE levels may represent an endogenous protective response to tissue injury, a driver of disease progression, or both. Indeed, NAEs exert neuroprotective and anti-inflammatory actions through CB_1_R/CB_2_R-and PPARα-dependent mechanisms in various preclinical models including excitotoxicity, stroke, multiple sclerosis and PD. The consequences of increased NAE levels are likely context dependent and shaped by cellular compartment, receptor expression, disease stage and, critically, the metabolic fate of each NAE species.^12–19^

NAE abundance is controlled by a network of biosynthetic and degradative pathways. Canonically, NAEs are produced from *N*-acylphosphatidylethanolamines (NAPEs) by NAPE phospholipase D,^20,21^ although alternative routes involving ABHD4, glycerophosphodiesterases, PLC-type enzymes and phosphatases also contribute to NAE formation.^1^ Termination of NAE signaling has been attributed mainly to hydrolysis of the amide bond,^22^ but oxidative degradation pathways of anandamide have also been reported.^23^ Fatty acid amide hydrolase (FAAH) is a membrane-associated serine hydrolase that is highly expressed in the brain and preferentially hydrolyses anandamide and other long-chain NAEs,^24^ whereas *N*-acylethanolamine-hydrolyzing acid amidase (NAAA) is a lysosomal cysteine hydrolase that preferentially degrades saturated and monounsaturated NAEs including PEA and OEA.^25^

Pharmacological inhibition of FAAH has emerged as a strategy to enhance endogenous NAE signaling without directly activating cannabinoid receptors.^1,2,26,27^ In rodents, FAAH inhibition increases AEA levels by approximately three- to sevenfold and PEA/OEA levels by approximately five- to twentyfold, producing beneficial effects in preclinical models of inflammation, neuropathic pain, PD and psychiatric disorders. ^28–30^ In humans, FAAH inhibitors have shown target engagement and increased circulating NAEs, but clinical translation has been limited.^31–41^

A striking observation is that NAE levels do not increase indefinitely when canonical degradation is blocked.^26,28,29^ Instead, inhibition of FAAH or NAAA results in a plateau in NAE accumulation. Pharmacokinetic and pharmacodynamic modelling of human FAAH inhibition indicates similarly that known hydrolytic and oxidative routes do not fully account for NAE turnover.^42^ This apparent metabolic ceiling implies that additional, underexplored pathways contribute to NAE homeostasis, particularly under conditions in which canonical hydrolysis is impaired or saturated. Identifying such pathways is essential, because they may determine whether NAEs remain bioactive signaling molecules, are stored in reservoirs, or are converted into metabolites with distinct biological properties.

One credible candidate pathway is glycosylation, a transformation that would connect NAE biology to the metabolism of glycosphingolipids. Glycosphingolipids are membrane lipids built on a ceramide backbone, comprising a fatty acyl amide-linked to a sphingoid base, coupled to one or more sugar residues.^43,44^ Ceramide itself is a central bioactive lipid involved in membrane organization, stress signaling, apoptosis, inflammation and cellular differentiation. Glucosylceramide (GlcCer) and galactosylceramide (GalCer) both serve as structural membrane components and as precursors for more complex glycosphingolipids.^43^

The catabolism of GlcCer is mediated by β-glucosylceramidases. GCase, encoded by the *GBA1* gene, is the lysosomal β-glucosylceramidase that hydrolyses GlcCer into ceramide and glucose. Homozygous loss-of-function mutations in *GBA1* cause Gaucher disease, the most common lysosomal storage disorder, in which GlcCer accumulates prominently within lysosomes of macrophages, giving rise to the characteristic Gaucher cells.^44–47^ Neurological manifestations occur in the neuronopathic forms of Gaucher disease, and heterozygous *GBA1* mutations, such as E326K (PD risk variant), N370S (PD mild variant), and L444P (PD severe variant), are among the strongest genetic risk factors for PD, being present in up to 30% of patients.^48,49^ These mutations reduce GCase activity, elevate GlcCer and glucosylsphingosine (GlcSph) levels, and promoteα-synuclein accumulation, contributing to neurotoxicity.^48–58^ These links place GlcCer metabolism at the core of lysosomal dysfunction, neuroinflammation and neurodegeneration.

A second enzyme, GBA2, encoded by *GBA2*, also hydrolyses GlcCer but its subcellular localization is distinct from GCase.^55^ Unlike lysosomal GCase, GBA2 is a non-lysosomal, membrane-associated β-GlcCer localized to the cytosolic face of intracellular membranes, including the endoplasmic reticulum, Golgi apparatus and plasma membrane.^59,60^ The existence of GBA2 helps explain why GlcCer handling is not restricted to lysosomes and why loss of GCase activity does not uniformly lead to GlcCer accumulation in all cell types. Thus, GCase and GBA2 define compartmentalized GlcCer metabolism: one operating within the degradative lysosomal system, the other at non-lysosomal membranes where lipid signaling and membrane remodeling are tightly coupled.

Importantly, the activity of GBA2 is not limited to hydrolysis. It can also transglycosylate, in which a sugar moiety from a glycosphingolipid donor is transferred to an acceptor lipid containing a hydroxyl group.^61–63^ Cholesterol is a known acceptor in this reaction, yielding glucosylated or galactosylated sterols.^58,64,65^ This catalytic versatility reveals an underappreciated chemical capacity of GBA2: rather than simply degrading glycosphingolipids, they can redistribute sugar groups across lipid classes and thereby generate new classes of metabolites.^66–68^

NAEs are structurally well suited to participate in such transglycosylation reactions. Similar to cholesterol, they contain a hydroxyl group that can serve as an acceptor for glycosyl transfer. Moreover, the ceramide backbone of GlcCer contains an *N*-acylethanolamine motif, providing a conceptual bridge between NAEs and glycosphingolipid biochemistry. We therefore hypothesized that NAEs may act as acceptor substrates for glucosylceramidase-mediated transglycosylation, leading to the formation of glycosylated NAEs (**Fig. 1**). This previously unrecognized potential mode of NAE metabolism, distinct from hydrolysis and oxidation, would directly couple endocannabinoid-related lipid signaling to glycosphingolipid turnover.

**Fig. 1.**
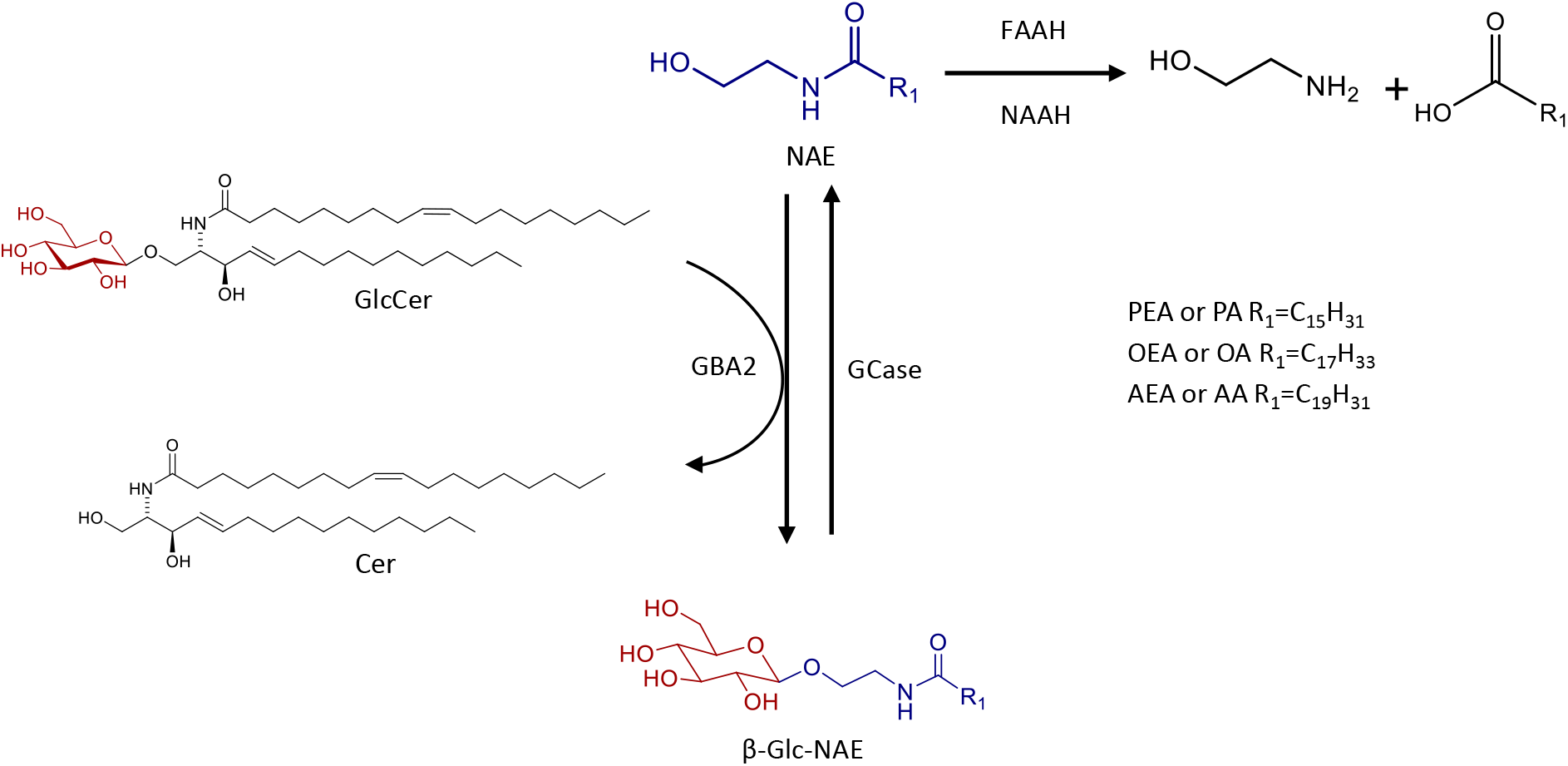
GBA2-dependent glycosylation connects *N*-acylethanolamine metabolism to glycosphingolipid turnover. The scheme illustrates glycosylation as a non-hydrolytic lipid-remodeling pathway. Bioactive *N*-acylethanolamines (NAEs), including AEA, OEA and PEA, can be hydrolyzed by FAAH or NAAA to their corresponding fatty acids (arachidonic acid (AA), oleic acid (OA) or palmitic acid (PA)). Non-lysosomal GBA2 transfers glucose or galactose from glycosphingolipid donors, such as glucosylceramide (GlcCer) or galactosylceramide (GalCer), to the hydroxyl group of NAEs, generating β-glucosylated or β-galactosylated NAEs. GCase removes the sugar moiety from the β-glycosylated-NAE.

In this study, we investigated whether GBA2 converts NAEs into glycosylated metabolites. We examine their formation and abundance in macrophages and neuronal cells, mouse tissues and disease-relevant contexts. Our findings reveal a new biochemical intersection between NAE signaling and glycosphingolipid metabolism, two lipid networks previously considered separate. We found that GBA2 transfers glucose or galactose to NAEs and that these β-glycosylated-NAEs are subsequently degraded by lysosomal GCase. This suggests that glycosylation can act as an alternative fate for bioactive fatty acid ethanolamides when canonical NAE degradation is impaired. More broadly, this work establishes GBA2-dependent glycosylation as a lipid-remodeling mechanism with potential relevance to inflammation, lysosomal dysfunction and neurodegenerative disease.

## Results

### Development and validation of LC-MS/MS method for β-Glycosylated NAE quantification

To enable the detection and quantification of β-glycosylated *N*-acylethanolamines (β-Glyco-NAEs), we first synthesized analytical standards for β-glucosyl (β-Glc)- and β-galactosyl (β-Gal)- AEA, OEA and PEA, together with their corresponding ^13^C_6_-labeled glycosylated internal standards (see methods). Next, we developed a targeted liquid chromatography (LC) mass spectrometry (MS)-MS -method to detect and quantify the β-Glyco-NAEs. Direct infusion of the synthetic compounds into the mass spectrometer showed that the protonated molecular ion [M+H]^+^ is the dominant adduct for all analytes (**Fig. 2a**). Upon collision-induced fragmentation, each compound predominantly lost its glucose or galactose moiety, yielding product ions corresponding to the parent NAE: m/z 348.3 for AEA, m/z 326.3 for OEA and m/z 300.3 for PEA.

**Fig. 2.**
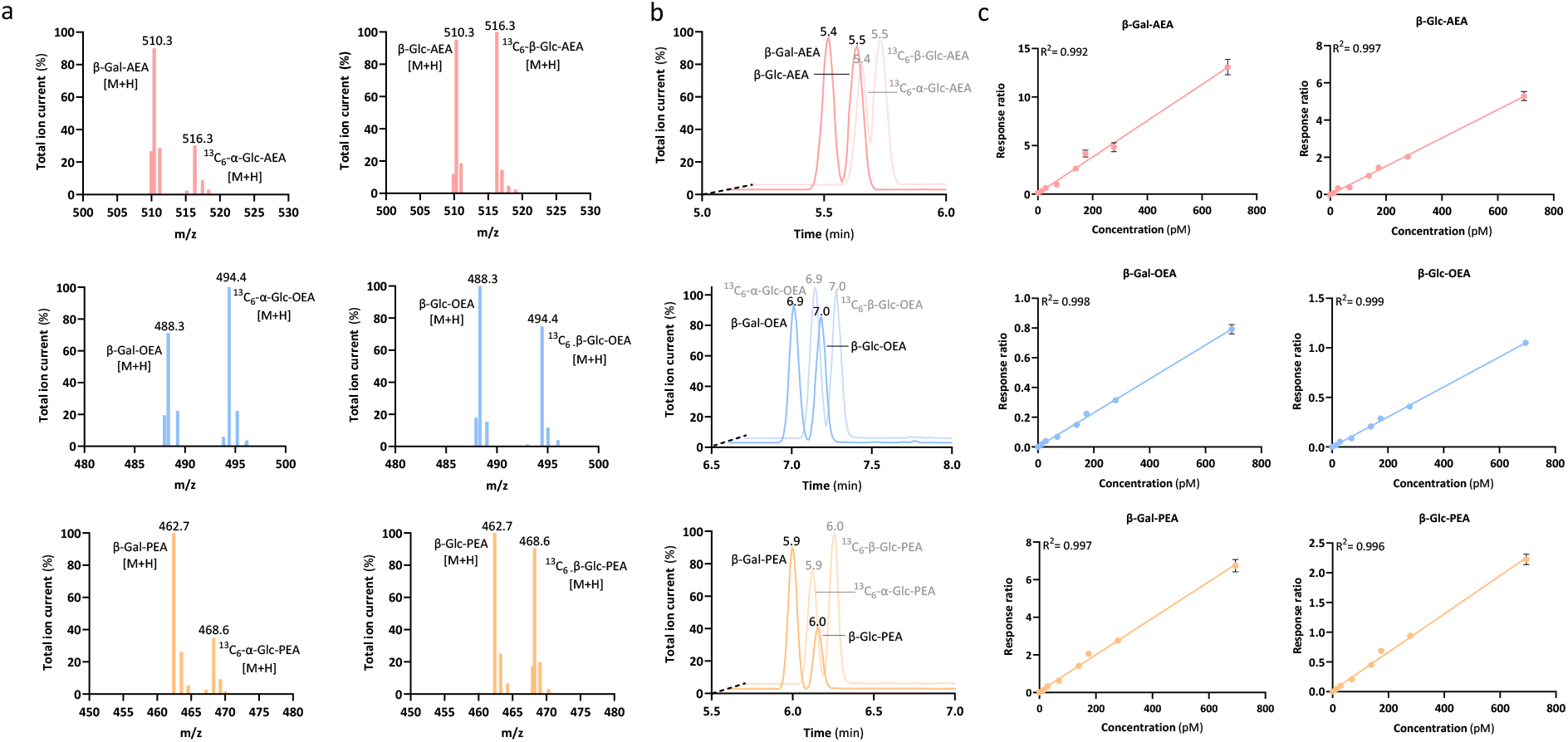
Development and validation of a targeted LC–MS/MS method for β-glyco-NAE quantification. **a,** Optimized multiple reaction monitoring transitions for β-Glc-AEA, β-Glc-OEA and β-Glc-PEA and the corresponding ^13^C_6_-labelled internal standards. Upon collision-induced fragmentation, β-glyco-NAEs predominantly lose the sugar moiety, yielding product ions corresponding to the parent NAE. **b,** Representative LC–MS/MS chromatograms showing chromatographic separation of β-Glc-NAEs and ^13^C_6_-labelled β-Glc-NAE internal standards. **c**, Calibration curves. Limits of detection and quantification, precision, matrix effects and recovery are shown in **Fig. S1**.

An LC method was developed to achieve baseline separation and accurate quantification of the glycosylated NAE species. Because these metabolites are more hydrophilic than their parent NAEs, the standard LC gradient^69^ was optimized to enhance retention and chromatographic resolution. This enabled clear separation of both β-Glc-NAEs and the corresponding ^13^C_6_-β-Glc-labeled standards (**Fig. 2b**). No interference was observed between unlabeled analytes and isotope-labeled internal standards, supporting the use of the ^13^C_6_-labeled standards for correction of extraction efficiency, ionization efficiency and chromatographic variation.

The LC-MS/MS method was then validated in mouse brain matrix. Blank samples, mouse brain extracts and spiked mouse brain were used to assess selectivity, specificity and carry-over. No carry-over was observed in blank injections, and no relevant crosstalk was detected between MRM transitions. Endogenous β-Glyco-NAE peaks were detected in mouse brain under the validation conditions (*vide infra*). Spiked samples showed clear analyte peaks at the expected retention times. Calibration curves were linear over the selected concentration range for all six β-Glyco-NAEs, with correlation coefficients above 0.99 (**Fig. 2c**). Limits of quantification (LOQ) were in the low picomolar range for most analytes, typically approximately 40–70 pM, with β-Gal-AEA showing a somewhat higher LOQ. These values indicate that the method is sufficiently sensitive for endogenous detection in biological matrices.

Precision was evaluated at low, medium and high concentrations over four days (**Fig. S1**). In mouse brain, intraday precision ranged from 9–28% for most compounds, whereas interday precision ranged from 12–42%. In human plasma, intraday precision ranged from 7–35% and interday precision from 8–38%. Precision at low concentrations was more variable, most likely because of matrix complexity and ion suppression, whereas precision at medium and high concentrations was generally within acceptable limits.

Matrix effects and extraction recovery were assessed using the ^13^C_6_-encoded internal standards (**Fig. S1**). Matrix effects ranged from 82–121% in mouse brain and 61–105% in human plasma. Recovery ranged from 21–41% from mouse brain and 37–64% from human plasma. Although recoveries were modest, the isotope-labeled internal standards compensated for losses during extraction and matrix-dependent ionization effects. Together, these data establish a targeted LC-MS/MS workflow for quantifying endogenous β-Glc- and β-Gal-NAEs in biological samples.

### GBA2 catalyzes transglycosylation of NAEs *in vitro*

Having established an analytical method for β-Glyco-NAE detection, we next investigated whether GBA2 can generate glycosylated NAEs. AEA, OEA and PEA were incubated with the glucosyl donor 4-methylumbelliferyl β-glucose (4MU-Glc) or 4-methylumbelliferyl β-galactose (4MU-Gal) and lysates from GBA2-overexpressing HEK293T cells. β-Glc-AEA, β-Glc-OEA and β-Glc-PEA (**Fig. 3a**) as well as their galactosyl counterparts (**Fig. 3b**) were detected in a dose-dependent manner upon increasing the NAE concentrations in the assay (**Fig. 3c,d**). Product formation was blocked by GBA2 inhibition with **JJB367** (200 nM) but not by GCase inhibition with **ME655** (50 nM) (**Fig. 3a,b**), demonstrating that GBA2 efficiently catalyzes NAE transglycosylation under these conditions. The relative efficiency of transglycosylation followed the order OEA > AEA > PEA and did not reach complete conversion at the highest NAE concentration tested.

**Fig. 3.**
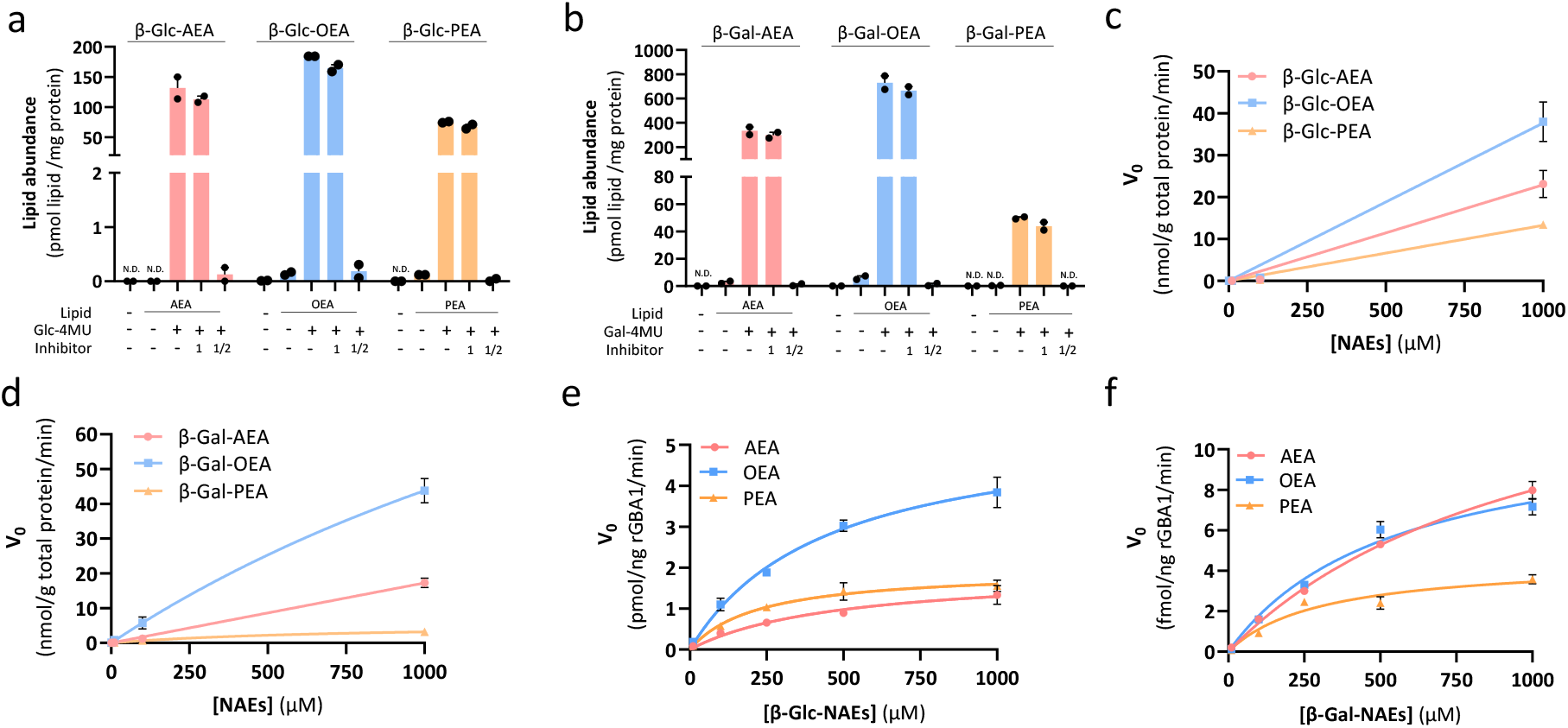
GBA2 forms β-Glyco-NAEs and GCase hydrolyses them *in vitro*. **a,** LC–MS/MS quantification of β-Glc-AEA, β-Glc-OEA and β-Glc-PEA or **b,** β-Gal-AEA, β-Gal-OEA and β-Gal-PEA formation after incubation of AEA, OEA or PEA with β-Glc-4MU and lysates from GBA2-overexpressing HEK293T cells. Reactions were performed in the presence or absence of GCase (#1, ME655, 50 nM) or GBA2 inhibitors (#1/2, JJB367, 200 nM) as indicated. Product formation was reduced by GBA2 inhibition but not by selective GCase inhibition (n=2). N.D. is not detected. **c**, concentration-dependent formation of β-Glc-NAEs or **d,** β-Gal-NAEs by HEK293 lysates overexpressing GBA2, **e,** Hydrolysis of β-Glc-AEA, β-Glc-OEA and β-Glc-PEA or **f,** β-Gal-AEA, β-Gal-OEA and β-Gal-PEA by recombinant GCase, quantified by formation of the corresponding parent NAEs. Data are shown as mean ± SEM; n=3 independent experiments unless otherwise indicated.

We next assessed whether β-Glyco-NAEs, just like β-GlcCer and β-Glc-cholesterol, are GCase substrates. Recombinant human GCase proved to accept β-Glc-AEA, β-Glc-OEA and β-Glc-PEA as well as their β-Gal-NAE counterparts as substrates, releasing the corresponding parent NAEs (**Fig. 3e,f**). Kinetic analysis yielded Michealis-Menten (K_M_) values in the range of approximately 200–500 μM and V_max_ values of approximately 1.9–5.6 pmol/ng/min for the β-Glc-NAEs, whereas the β-Gal-NAEs were 1000-fold less efficiently metabolized by GCase (**Table 1**). GCase hydrolyzes β-Glc-OEA most efficiently.

**Table 1.** Enzyme kinetic parameters for β-Glyco-NAE hydrolysis. Kinetic parameters for GCase-mediated hydrolysis of β-Glyco-NAEs. Values are reported as mean ± SEM (n=3). With K_M_ in µM, V_max_ in pmol/ng/min, k_cat_ in min^-1^ and k_cat_/K_M_ in M^-1^s^-1^.

| | β-Glc-AEA | β-Glc-OEA | β-Glc-PEA | β-Gal-AEA | β-Gal-OEA | β-Gal-PEA |
| --- | --- | --- | --- | --- | --- | --- |
| $K_M$ | 483 ± 190 | 442 ± 96 | 221 ± 61 | > 1000 ± 1713 | 542 ± 131 | 306 ± 110 |
| $V_{max}$ | 1.9 ± 0.3 | 5.6 ± 0.5 | 2.0 ± 0.2 | 0.016 ± 0.002 | 0.011 ± 0.001 | 0.005 ± 0.001 |
| $k_{cat}$ | 3919 ± 710 | 11348 ± 1096 | 3979 ± 373 | 33 ± 3.3 | 23 ± 2.7 | 9.2 ± 1.3 |
| $k_{cat}/K_M$ | 1.35E+05 | 4.28E+05 | 3.01E+05 | 6.80E+02 | 7.55E+02 | 5.32E+02 |

### β-Glc-NAEs are formed in living cells through a GBA2-dependent pathway

We next examined whether β-Glc-NAEs are formed in intact cells. Wildtype HEK293T cells were incubated with AEA, OEA or PEA, with or without GCase- or GBA2 selective inhibitors. All three β-Glc-NAEs were detected after NAE feeding (**Fig. 4a**). Inhibition of GBA2 (**MZ31**, 25 nM) reduced β-Glc-NAE formation, whereas inhibition of GCase (**ME655**, 50 nM) did not, indicating that GBA2 is the principal enzyme responsible for cellular β-Glc-NAE production in this system. Similar results were obtained for the cellular production of β-Gal-NAEs (**Fig. 4b**). In contrast to the *in vitro* lysate experiment, AEA was most efficiently glycosylated in intact HEK293T cells, possibly because β-Glyco-AEA is less efficiently hydrolyzed by GCase and/or because OEA and PEA are more rapidly metabolized by other NAE-degrading enzymes.

**Fig. 4.**
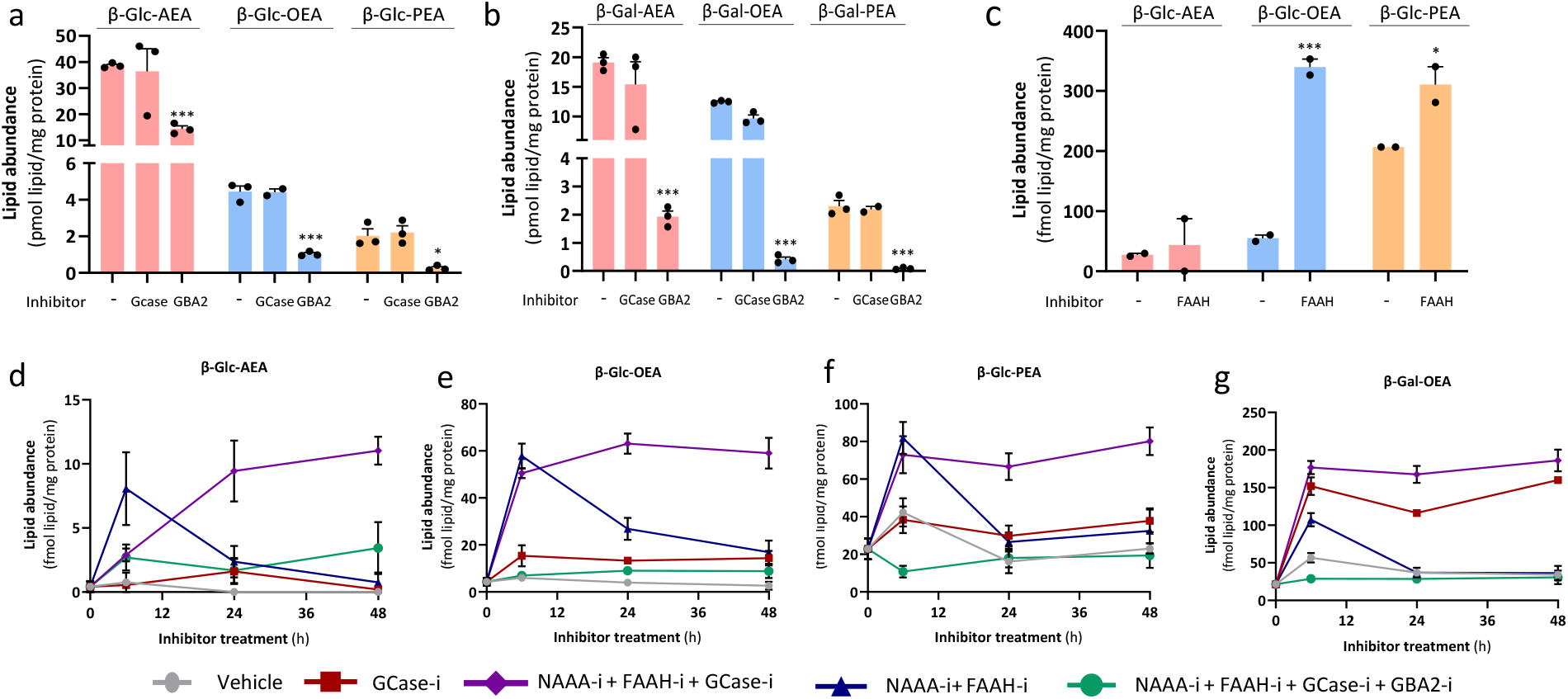
Endogenous β-Glyco-NAE formation in cells depends on GBA2 and parent NAE availability. **a,** Formation of β-Glc-AEA, β-Glc-OEA and β-Glc-PEA and **b**, β-Gal-AEA, β-Gal-OEA and β-Gal-PEA in wild-type HEK293T cells after feeding with AEA, OEA or PEA in the presence or absence of GCase or GBA2 inhibitors. **c,** Endogenous β-Glc-NAE levels in differentiated Neuro-2a cells treated with vehicle or the FAAH inhibitor PF-04457845 (1 μM, 48 h). **d,** Time-dependent β-Glc-NAE and β-Gal-OEA accumulation in RAW264.7 macrophages treated with inhibitors of FAAH (PF04457845, 1 µM), NAAA (ARN19702, 5 µM), GCase (ME655, 50 nM) and/or GBA2 (MZ31, 25 nM), as indicated. Data are shown as mean ± SEM; n=3 for HEK293T cells, n=2 for Neuro-2a cells and n=4 for RAW264.7 macrophages, unless otherwise indicated. Statistical significance was determined by two-way ANOVA with *p < 0.05, **p < 0.01, ***p < 0.001 compared to vehicle (no inhibitor).

To determine whether β-Glc-NAEs can be produced endogenously, we analyzed differentiated Neuro-2a cells, a neuronal cell model known to produce NAEs upon retinoic acid-induced differentiation. Differentiated Neuro-2a cells contained endogenous β-Glc-OEA and β-Glc-PEA, and lower levels of β-Glc-AEA, whereas β-Gal-NAEs were not detected (**Fig. 4c**). Inhibition of FAAH with **PF04457845** (1 µM) further increased β-Glc-OEA and β-Glc-PEA levels (**Fig. 4c**), indicating that increasing the availability of parent NAEs enhances flux into the glucosylation pathway.

We also analyzed endogenous β-Glc-NAE formation in mouse RAW264.7 macrophages over time. Basal β-Glc-AEA, β-Glc-OEA and β-Glc-PEA levels were low but detectable (**Fig. 4d-f**). Inhibition of GCase (**ME655**, 50 nM), or combined inhibition of FAAH (**PF04457845**, 1 µM) and NAAA (**ARN19702**, 5 µM), increased β-Glc-NAE levels. Combined inhibition of FAAH, NAAA and GCase produced a stronger and more sustained increase over 48 hours then treatment with inhibitors on their own. (**Fig. 4d-f**). GBA2 inhibition (**MZ31**, 25 nM) prevented this accumulation, demonstrating that endogenous β-Glc-NAE production in these macrophages depends on GBA2. Similar results were found for β-Gal-OEA (**Fig. 4g**), but β-Gal-PEA and β-Gal-AEA were not detected. These findings indicate that endogenous cellular β-Glc-NAE levels are shaped by two opposing processes: GBA2-mediated formation from NAEs and GCase-mediated hydrolysis of β-Glc-NAEs. Furthermore, FAAH/NAAA limits NAE substrate availability for GBA2, thereby controlling β-Glc-NAE formation.

### β-Glyco-NAEs are endogenous metabolites in mouse tissues

We next asked whether β-Glyco-NAEs are present *in vivo*. Lipids extracted from nine tissues from male C57BL/6J mice were analyzed by LC-MS/MS. Levels of β-Glyco-NAEs, NAEs, GlcCer, GlcChol and GalCer were quantified in brain, heart, lung, liver, kidney, intestine, spleen, testes and pancreas. β-Glc-NAEs and β-Gal-NAEs were detected across the tissue panel, although abundance varied markedly between tissues (**Fig. 5a,b**). β-Glc-OEA was the most abundant β-Glc-NAE overall, with particularly high levels in intestine and brain. β-Gal-OEA was the most abundant β-Gal-NAE (brain) (**Fig. 5b**). In intestine, approximately 7% of the OEA pool was estimated to be present in its glucosylated form (**Fig. 5c**), whereas in the brain and spleen approximately 10 and 60% was galactosylated (**Fig. 5d**). β-Glc-PEA was detected in all tissues and was most abundant in intestine, brain, spleen and testes. β-Glc-AEA was the least abundant species, consistent with the relatively (to OEA and PEA) lower abundance of AEA itself but showed its highest levels in brain.

**Fig. 5.**
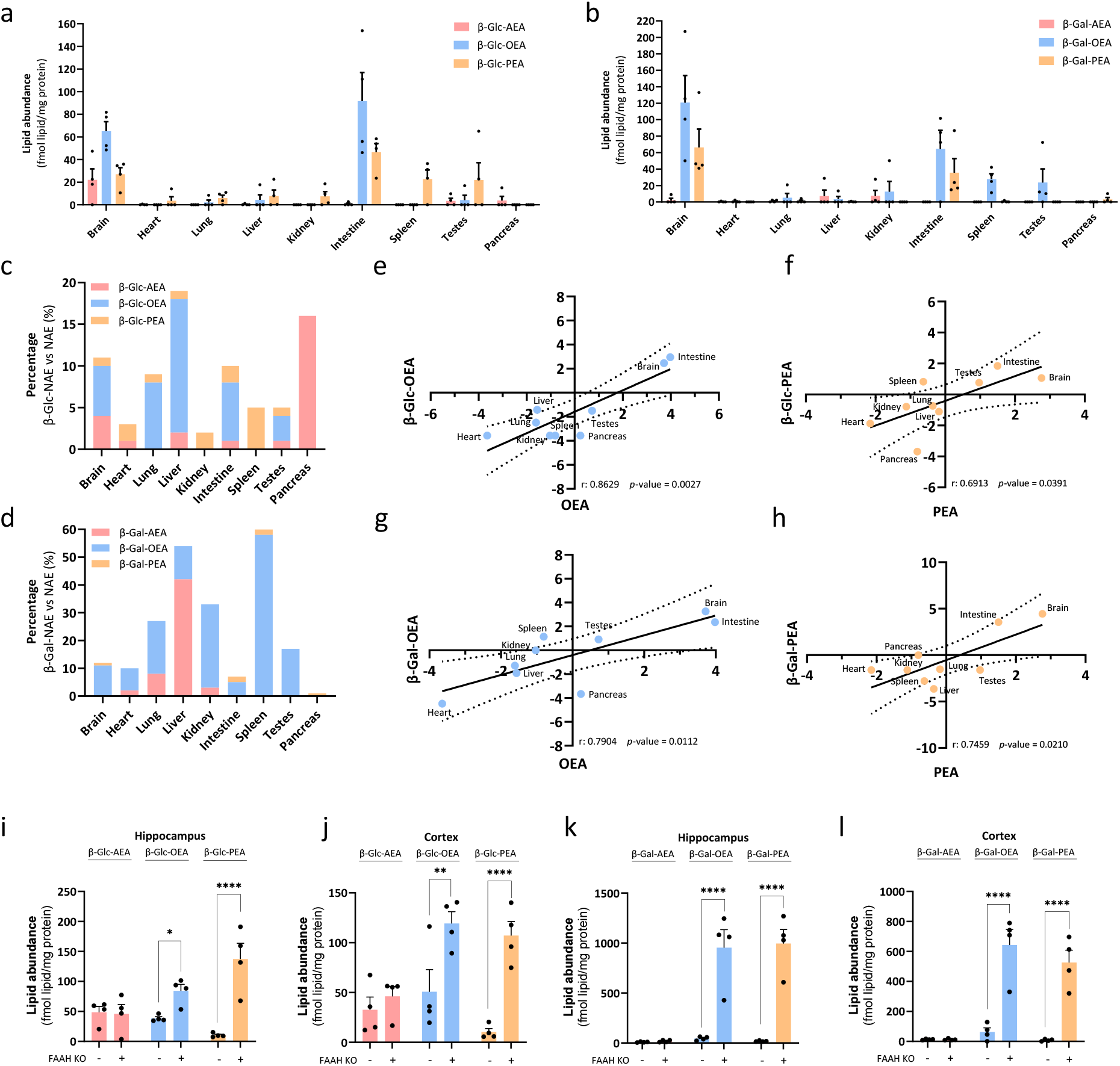
β-Glyco-NAEs are endogenous metabolites in mouse tissues and are regulated by NAE and glucosylceramidase metabolism in vivo. **a,** Tissue distribution of β-Glc-AEA, β-Glc-OEA and β-Glc-PEA, **b**, β-Gal-AEA, β-Gal-OEA and β-Gal-PEA in mouse brain, heart, lung, liver, kidney, intestine, spleen, testes and pancreas. Data is shown as mean ± SEM (N=4). **c,** Ratio of β-Glc-NAEs or **d**, β-Gal-NAEs over their parent NAEs expressed as percentage in distinct mouse tissues **e-f,** Correlation plots of β-Glc-NAE and **g, h,** β-Gal-NAE levels versus their parent NAEs in mouse tissues. All data is represented as Log2 of the relative amount. A two-tailed Pearson correlation analysis was performed and the dotted lines represent 95% confidence interval of the best fitted line (solid line) as determined by linear regression analysis. **I-l,** β-Glc-NAE and β-Gal-NAE levels in mouse hippocampus (**i, k**) and cortex (**j, l**) from FAAH-knockout mice and corresponding controls. **i-j** Data are shown as mean ± SEM; N=4 for all tissues with ** p < 0.01 and *** p < 0.001 control wildtype versus FAAH KO (two-way ANOVA).

The distribution of β-Glyco-NAEs generally reflects the abundance of their parent NAEs, suggesting that NAE availability is an important determinant of β-Glyco-NAE formation in vivo (**Fig. 5e-h**). In contrast, levels of the potential glycosyl donors, GlcCer, GalCer and GlcChol did not closely correlate with β-Glyco-NAE abundance across tissues (**Fig. S2**). GlcCer levels were 100- to 1000-fold higher than glucosylated cholesterol, indicating that GlcCer is likely the main endogenous sugar donor, but that donor abundance alone does not explain tissue-specific β-Glc-NAE levels.

To test whether FAAH limits the availability of NAEs for β-Glyco-NAE formation *in vivo*, we quantified β-Glc-NAEs and β-Gal-NAEs in brain from FAAH-knockout (KO) mice (**Fig. 5i-l**). Both in hippocampus and cortex β-Gal-OEA and β-Gal-PEA levels were highly upregulated (> 500-fold) followed by their glucosylated counterparts (< 100-fold), whereas the β-Glyco-AEA was not significantly increased compared to wildtype controls.

### β-Glc-NAE levels are elevated in Gaucher disease patient samples

Because GCase hydrolyzes β-Glc-NAEs *in vitro* and *in vivo*, we investigated whether impaired GCase activity leads to β-Glc-NAEs accumulation under pathological conditions in human samples. We quantified β-Glc-NAEs in spleen and plasma from patients with type 1 Gaucher disease and control donors. β-Glc-NAEs were detected at low levels in control samples. In Gaucher disease spleen, β-Glc-AEA and β-Glc-PEA were significantly increased, whereas β-Glc-OEA showed no significant change. In plasma from Gaucher disease patients, β-Glc-AEA, β-Glc-OEA and β-Glc-PEA were increased compared with controls. Furthermore, we have identified these β-Glc-NAEs in the plasma of PD-patients carrying heterozygous E326K, N370S and L444P *GBA1*-mutations. Increased β-Glc-PEA and β-Gal-OEA plasma levels appeared to be associated with age of *GBA1*-associated PD patients, but not in *non-GBA* PD or controls (**Fig. 6d,e**). These data indicate that β-Glc-NAEs and β-Gal-NAEs are endogenous human metabolites and that their levels are elevated in the context of GCase deficiency and with age.

**Fig. 6.**
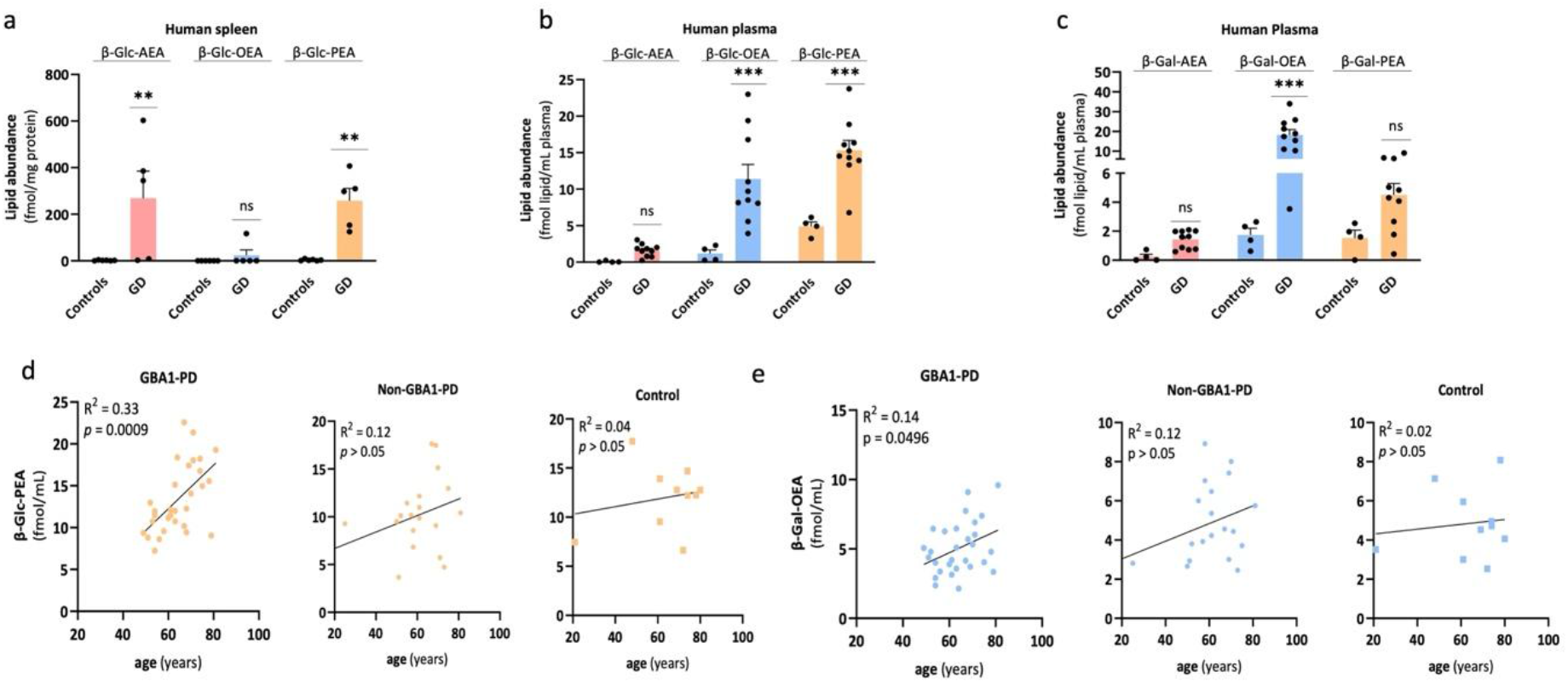
β-Glc-NAEs accumulate in human disease settings associated with GCase deficiency. **a,** β-Glc-AEA, β-Glc-OEA and β-Glc-PEA levels in spleen and **b,** plasma samples from patients with type 1 Gaucher disease and control donors. **c,** β-Gal-AEA, β-Gal-OEA and β-Gal-PEA levels in plasma samples from patients with type 1 Gaucher disease and control donors. N=6 for controls and N=5 for GD type 1, technical n =2, and human plasma N=4 for control and N=10 for GD type 1, technical n=2. With **p < 0.01 and *** p < 0.001 control versus GD patients (two-way Anova) **d,** Plasma β-Glc-PEA and **e,** β-Gal-OEA levels in patients with Parkinson’s disease stratified by *GBA1* mutation status and controls. β-Glyco-NAE levels are shown in relation to age in *GBA1*-associated Parkinson’s disease (N=20), non-*GBA* Parkinson’s disease (N=20) and controls (N=10). Data are shown as mean ± SEM

### β-Glyco-NAEs potentiate pro-inflammatory responses

Finally, we examined whether β-Glyco-NAEs retain the signaling activity of their parent NAEs. AEA and β-Glc-AEA were tested in CB_1_R and CB_2_R binding assays and in a TRPV1 activation assay. Whereas AEA interacts with these targets, β-Glc-AEA did not show detectable CB_1_R and CB_2_R binding or TRPV1 activation (**Fig. 7a,b**). Similarly, PEA activated PPARα in a reporter assay, whereas β-Glc-PEA did not (**Fig. 7c**). These results indicate that glucosylation prevents interaction with the canonical receptors of the parent NAEs. Similar results were obtained for the β-Gal-NAEs (**Fig. 7a-c**). Thus, NAE glycosylation may function as an inactivation pathway for CB_1_R-, CB_2_R-, TRPV1- and PPARα-mediated signaling.

**Fig. 7.**
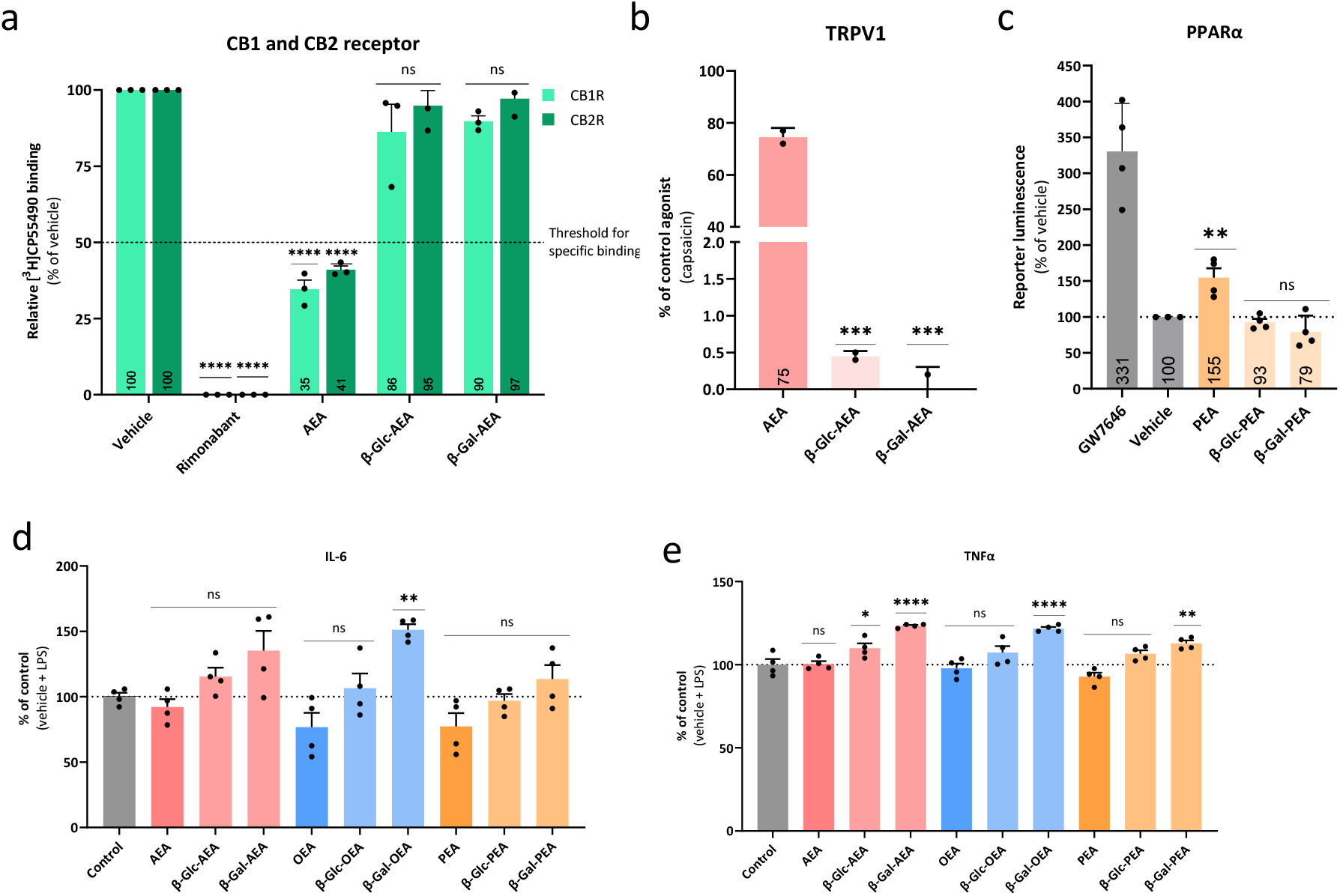
Glycosylation prevents canonical NAE receptor engagement but enhances inflammatory cytokine responses. **a,** CB1R and CB2R binding assays comparing AEA (10 μM) with β-Glc-AEA (10 μM) and β-Gal-AEA (10 μM) with rimonabant (10 μM) as positive control (n=3). **b,** TRPV1 activation assay comparing AEA (10 μM) with β-Glc-AEA (10 μM) and β-Gal-AEA (10 μM) (n=2) **c,** PPARα reporter assay comparing PEA (25 μM) with β-Glc-PEA (25 μM) with GW7646 (1 μM) as positive control (N=4, n=3). Cytokine responses in LPS-stimulated (1 ng/mL) microglial cells treated with NAEs (10 μM) or β-Glyco-NAEs (10 μM) for 24h, showing effects on pro-inflammatory cytokines **d,** IL-6 and **e,** TNFα. Data is expressed as percentage of control (vehicle + LPS) and represented as mean ± SEM with * p < 0.05, ** p < 0.01 and *** p < 0.001 (N=4, n=2) (one-way Anova, Dunnet’s multiple comparisons test);

Since GBA2 inhibitors are known to have anti-inflammatory effects and prevent β-Glyco-NAE formation, we tested whether β-Glyco-NAEs may have any intrinsic pro-inflammatory properties themselves. To this end, we incubated N9 microglial cells stimulated with LPS and β-Glyco-NAEs and found that the release of both pro-inflammatory cytokines IL6 and TNFα into the medium was significantly elevated by β-Gal-OEA, whereas β-Gal-PEA and β-Gal-AEA only increased TNFα levels and the β-Glc-NAEs did not elevate cytokine levels (**Fig 7d,e**).

## Discussion

This study identifies glycosylation as an novel branch of *N*-acylethanolamine (NAE) metabolism and places GBA2 at the core of this pathway. Rather than acting solely as a glucosylceramidase, GBA2 can remodel bioactive NAEs by transferring glucose or galactose to AEA, OEA and PEA. The detection of β-Glyco-NAEs in macrophages and neuronal cells, mouse tissues and human samples indicates that this chemistry is not an artefact of *in vitro* enzymology but represents an endogenous lipid-remodeling pathway that links NAE signaling to glycosphingolipid metabolism.

Conceptually, this pathway differs from the established routes of NAE inactivation by FAAH and NAAA. FAAH and NAAA terminate NAE signaling by hydrolyzing the amide bond, thereby destroying the NAE scaffold and releasing ethanolamine and a fatty acid.^24,25^ In contrast, GBA2-mediated glycosylation preserves the *N*-acylethanolamine backbone, however, β-Glyco-NAEs no longer engage CB_1_R, CB_2_R, TRPV1 or PPARα, suggesting that glycosylation may attenuate NAE receptor signaling.

The NAE glycosylation pathway appears to be governed by a balance between non-lysosomal formation and lysosomal turnover. GBA2 generates β-Glyco-NAEs in the cytoplasm, whereas GCase can hydrolyze these metabolites back to their parent NAEs in the lysosome. This opposing activity provides a compartmentalized mechanism for controlling β-Glyco-NAE abundance and connects cytosolic or membrane-associated lipid signaling with lysosomal lipid catabolism. The accumulation of β-Glyco-NAEs in Gaucher and Parkinson’s disease samples, where GCase activity is reduced, supports this model and broadens the metabolic consequences of GCase deficiency beyond GlcCer and GlcSph storage. Furthermore, we propose that glycosylation of NAEs by GBA2 may provide an alternative cellular disposal mechanism of NAEs that is operational at high cellular NAE concentrations, such as found during differentiation, inflammation and neurodegeneration.^1,15,70–74^ The glycosyl group of β-Glyco-NAEs may serve to solubilize the NAE or as a tag for its transportation, possibly via yet unknown binding or transporter proteins, into the lysosome, where they will be degraded by GCase and NAAA. This alternative metabolic pathway may provide a potential explanation why NAE levels do not rise indefinitely when FAAH is inhibited.^42^ This could have potential consequences for the clinical studies with FAAH inhibitors, especially at high and multi-day dosing^26^, or in patient populations with reduced GCase activity.^48–50^

Whether β-Glyco-NAEs are merely inactive clearance products or have biological or pathological functions of their own remains an important question. Their lack of activity at CB1R, CB2R, TRPV1 and PPARα supports a role in signal termination, whereas their effects on the release of inflammatory cytokines suggest that β-Glyco-NAEs may also engage innate immune pathways. Further glycan modifications could generate more complex glycolipid species capable of interacting with C-type lectin receptors, such as MINCLE, or other carbohydrate-recognition receptors.^75,76^ In addition, β-Glyco-NAEs may influence membrane organization, lipid trafficking or inflammatory signaling, or engage currently unknown protein targets. Thus, glycosylation may not simply switch NAE signaling off but redirect it into a distinct biochemical and cellular context especially when GCase activity is reduced. Given the strong genetic link between GCase mutations and Parkinson’s disease^49–57^, altered β-Glyco-NAE metabolism may also be relevant in neurodegenerative settings where lysosomal function, glycosphingolipid turnover and NAE signaling are perturbed.

Several questions remain unresolved. The physiological sugar donors for β-Glyco-NAE formation need to be defined, although GlcCer and GalCer are clear candidates. In addition, β-Gal-NAEs appear to be less efficiently hydrolyzed by GCase than β-Glc-NAEs, raising the possibility that they differ in stability, turnover or biological function. In this regard, it is noteworthy that β-Gal-OEA compared to β-Gal-AEA and β-Gal-PEA was the most abundant species identified in mice tissue (brain, intestines), highly increased in the hippocampus and cortex of FAAH knockout animals, found in human plasma of Gaucher patients, and also the most potent β-Glyco-NAE stimulating the release of IL6 and TNFα. Are there other enzymes involved in their processing, such as galactocerebrosidase (GalC), and are they neurotoxic? Are the β-glyco-NAEs in human plasma from GD and PD-patients reflecting what is occurring in the brain? Finally, the therapeutic relevance of the pathway could be tested using genetic and pharmacological perturbation of GBA2 combined with quantitative lipidomics and disease-relevant phenotyping.

The here-presented research focused on AEA, OEA and PEA, which comprise the best-characterized NAEs. The NAE family is however structurally diverse, and GBA2 may accept additional Indeed, β-glucosylated laureoylethanolamide (C12:0)^77^ was identified in the plant *Arabidopsis thaliana* and UDP-glycosyltransferase 76G1 (UGT76G1) from the plant *Stevia rebaudiana* was able to glucosylate AEA and docosahexaenoylethanolamide (DHEA).^78^ Expanding the analytical platform to a broader range of NAE species will be important to determine whether glycosylation selectively regulates signaling lipids or functions as an overflow pathway when canonical NAE degradation is impaired. Taken together, our findings establish β-Glyco-NAEs as endogenous metabolites and define GBA2-dependent glycosylation as a new biochemical link between endocannabinoid-related signaling, glycosphingolipid metabolism and lysosomal disease.

## Methods

### Chemicals and reagents

ULC/MS-grade acetonitrile and water, HPLC-grade methanol, tert-butyl methyl ether (MTBE) and formic acid were purchased from Biosolve. Unless stated otherwise, all other reagents were obtained from Sigma-Aldrich/Merck. AEA, OEA and PEA were obtained from commercial suppliers or synthesized in-house where indicated. The following inhibitors were used: the GCase inhibitor ME655 (synthesized in house),^79^ the GBA2 inhibitor MZ31 (synthesized in house),^8080^ the dual GCase/GBA2 inhibitor JJB367 (synthesized in house),^81^ the FAAH inhibitor PF-04457845 (Sigma) and the NAAA inhibitor ARN726 (synthesized in house)^82^ Glc-4MU and Gal-4MU (Glycosynth and Biosynth respectively) were used as artificial glycosyl donors for *in vitro* transglycosylation assays.

**Supplementary scheme S1:**
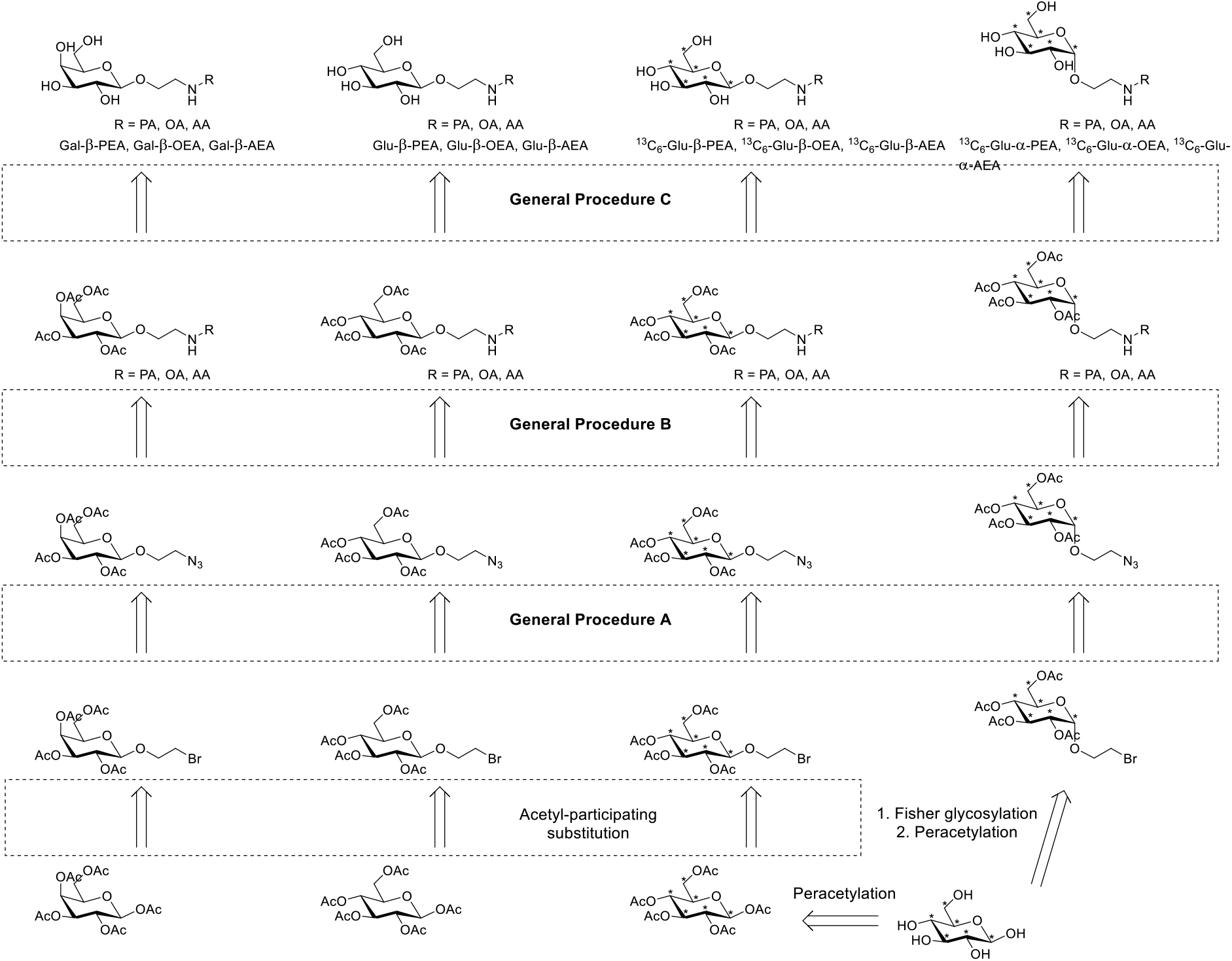
General synthesis route for glycol-NAEs. PA, palmitic acid; OA, oleic acid; AA, arachidonic acid.

### Synthetic procedures

β-Glc- and β-Gal-NAE standards, together with ^13^C_6_-labelled β-Glc- and α-Glc-NAE internal standards, were synthesized as follows. Reagents were purchased from Sigma Aldrich, Acros, Alfa Aesar or Fluka and were used without further purification, unless otherwise stated. [^13^C_6_]-D-glucopyranose (¹³C₆, 99%) was purchased from Cambridge Isotope laboratories Inc. and used as received. Solvents were purchased from Sigma Aldrich, Biosolve, Fluka or Honeywell and were dried over 3Å activated molecular sieves (8 to 12 mesh, Acros Organics), unless otherwise stated, and stored under N_2_ atmosphere. All moisture sensitive reactions were performed under nitrogen atmosphere. Reactions were followed by thin layer chromatography (TLC) using 0.20 mm silica gel 60 UV_254_, supplied by Macherey-Nagel. Compounds were visualized using KMnO_4_ stain (KMnO_4_ (6 g), K_2_CO_3_ (40 g), H_2_O (600 mL)) or CAM stain (Ce(NH_4_)_4_(SO_4_)_4_.2H_2_O (ceric ammonium sulfate :10 g); ammonium molybdate (25 g); conc. H_2_SO_4_ (100 mL); H_2_O (900 mL)). Column chromatography purification was performed on silica gel 60 Å (0.04 – 0.063 nm, Macherey-Nagel) or with Biotage Isolera Four 2.0.6 on SiliaSep^TM^ HP flash cartridges (4 g or 12 g, UltraPure irregular silica gel 60 Å, 40-63 μm, pH = 6.5-7.5, Screening Devices). Liquid chromatography-mass spectrometry (LC/MS) spectra were recorded on a Vanquish UHPLC system (Thermo Scientific) equipped with a C18 column (50 x 4.6 mm, 3 µm; Nucleodur Gravity, Macherey-Nagel). High-resolution mass spectra (HRMS) were recorded on a Thermo Scientific Q Exactive HF Orbitrap mass spectrometer equipped with an electrospray ion source in positive mode (source voltage 3.5 kV, sheath gas low 10, capillary temperature 275 °C) with resolution R = 60.000 at m/z = 400 (mass range = 150-1500). ^1^H and ^13^C NMR spectra were recorded on a Bruker AV400 (400 MHz), a DRX500 (500 MHz), an AV600 (600 MHz) or an AV850 (850 MHz) instrument. NMR spectra were recorded in deuterated chloroform (CDCl_3_) or methanol (methanol-d_4_). Chemical shift values are reported in ppm with solvent resonance as internal standard (CDCl_3_: δ 7.26 for ^1^H, δ 77.16 for ^13^C; methanol-d_4_: δ 3.31 for ^1^H, δ 49.00 for ^13^C). Interpretation of NMR data was achieved using Bruker TopSpin 1.3 and MestreNova 12.0. Data are reported as follows: chemical shifts (δ), multiplicity (s = singlet, d = doublet, t = triplet, q = quartet, p = pentet, bs = broad singlet, dd = double doublet, dt = double triplet, td = triple doublet, tt = triple triplet, m = multiplet) coupling constants *J* (Hz) and integration. NMR peak assignment was achieved using COSY and HSQC. Additional HMBC measurements were taken, if deemed necessary. All given ^13^C-APT spectra are ^1^H decoupled. All given ^1^H spectra for ^13^C-labelled compounds are ^13^C-decoupled.

**General procedure A:** Bromoethylglycoside substitution with sodium azide

To a solution of 2’-bromoethyl 2,3,4,6-tetra-*O*-acetyl-D-galacto/glucopyranoside (1 equiv.) in dry DMF (0.1 M) was added NaN_3_ (10 equivalents). The reaction was stirred overnight at ambient temperature, subsequently filtered and the filtrate diluted with water (100 mL). The mixture was extracted with EtOAc (3×50 mL) and the combined organic layers were washed with H_2_O (2×100 mL), brine (3×100 mL), dried (MgSO_4_), filtered and concentrated under reduced pressure.

**General procedure B:** Reduction of azide group and subsequent amide coupling

**Step 1:** In a microwave vial PtO_2_ (0.4 equiv.) and N_2_-purged EtOH (3 mL) were added and placed under N_2_ atmosphere. The catalyst was activated under H_2_ atmosphere, and the vial was subsequently placed under N_2_ atmosphere. To this solution was added 2’-azidoethyl 2,3,4,6-tetra-*O*-acetyl-D-galacto/glucopyranoside (1 equiv.) dissolved in N_2_-purged EtOH (3 mL). The reaction was placed under H_2_ atmosphere and stirred until TLC analysis indicated full conversion of the starting material (typically 16 hours). The reaction mixture was filtered through a celite pad and the pad was rinsed with EtOH. The filtrate was concentrated to afford the corresponding 2’-aminoethyl 2,3,4,6-tetra-*O*-acetyl-α/β-D-galacto/glucopyranoside, which was used in the next step without further purification.

**Step 2:** To a cooled solution (0 °C) of OSu-ester (1 equiv.) in dry DCM (0.02 M), DIPEA (5 equiv.) was added. 2’-Aminoethyl 2,3,4,6-tetra-*O*-acetyl-D-galacto/glucopyranoside (1 equiv., 0.02 M in dry DCM). was added dropwise. The reaction was allowed to reach room temperature and it was stirred until TLC analysis indicated full conversion of the starting material (typically 1 hour). The solvent was evaporated under reduced pressure and purification of the residue with silica gel column chromatography (20→80% EtOAc in pentane) afforded the corresponding 2’-amidethyl 2,3,4,6-tetra-*O*-acetyl-α/β-D-galacto/glucopyranoside.

**General Procedure C:** Global deprotection of glycolipids

2’-NAE-amidethyl 2,3,4,6-tetra-*O*-acetyl-α/β-D-galacto/glucopyranoside (1 equiv.) was dissolved in MeOH (0.03 M, distilled over Mg/I_2_ and stored over 3 Å molecular sieves). A catalytic amount of NaOMe was added until the reaction mixture reached pH = 9 – 10. The reaction was stirred until TLC analysis indicated full conversion of the starting material (typically 2 hours). The reaction was neutralized using resin Amberlite 120H^+^ and the resin was filtered off and rinsed with MeOH (10 mL). The filtrate was concentrated under reduced pressure and the residue purified with silica gel column chromatography (1→15% MeOH in DCM) to afford the corresponding deprotected 2’NAE-amidethyl α/β-D-galacto/glucopyranoside.

### Example of NMR peak assignment

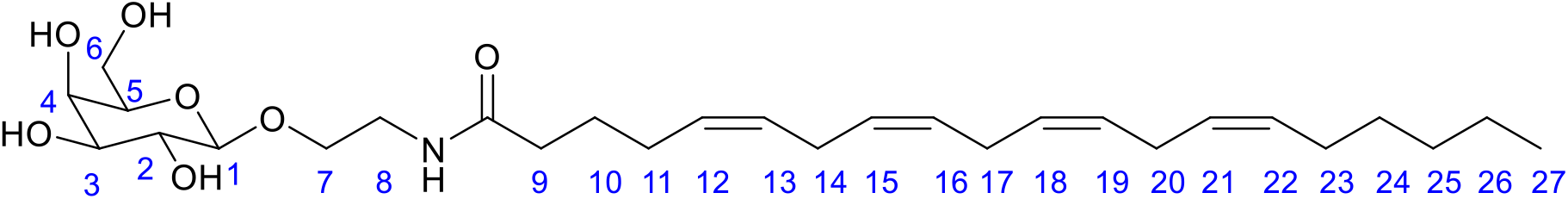

### 2,5-Dioxopyrrolidin-1-yl palmitate (1)

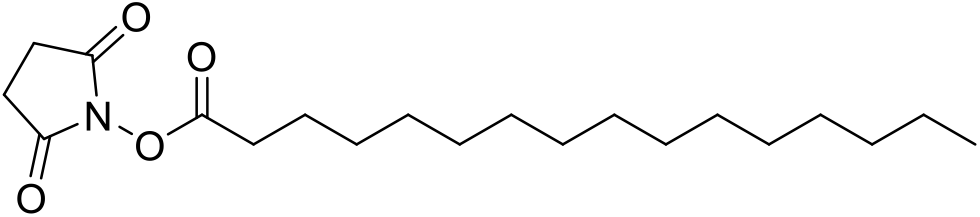

Palmitic acid (500 mg, 1.95 mmol) was dissolved in dry DCM (49 mL). NHS (337 mg, 2.92 mmol), EDC (561 mg, 2.92 mmol) and DIPEA (511 μL, 2.92 mmol) were added and the mixture was stirred overnight at room temperature. The mixture was diluted with brine (20 mL) and extracted with DCM (3×50 mL). The combined organic layers were dried (Na_2_SO_4_), filtered, and concentrated to afford the title compound as a white solid (0.689 g, 1.95 mmol, 100%). R*_f_* = 0.8 (1:1 EtOAc/pentane). ^1^H NMR (400 MHz, CDCl_3_) δ 2.84 (d, *J* = 4.5 Hz, 4H), 2.60 (t, *J* = 7.5 Hz, 2H), 1.74 (p, *J* = 7.5 Hz, 2H), 1.45 – 1.34 (m, 2H), 1.25 (bs, 22H), 0.88 (t, *J* = 7.0 Hz, 3H). ^13^C NMR (101 MHz, CDCl_3_) δ 32.08, 31.09, 29.83, 29.81, 29.77, 29.70, 29.51, 29.24, 28.95, 25.74, 24.72, 14.28.

### 2,5-Dioxopyrrolidin-1-yl oleate (2)

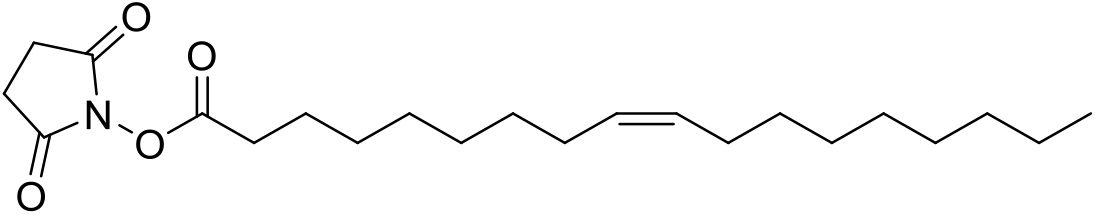

Oleic acid (477 mg, 1.69 mmol) was dissolved in dry DCM (42 mL). NHS (291 mg, 2.53 mmol), EDC (485 mg, 2.53 mmol) and DIPEA (442 μL, 2.53 mmol) were added and the mixture was stirred overnight at room temperature. The mixture was diluted with brine (20 mL) and extracted with DCM (3×50 mL). The combined organic layers were dried (Na_2_SO_4_), filtered, and concentrated to afford the title compound as a waxy yellow solid (0.641 g, 1.69 mmol, 100%). R*_f_* = 0.7 (1:1 EtOAc/pentane). ^1^H NMR (400 MHz, CDCl_3_) δ 5.40 – 5.29 (m, 2H), 2.83 (d, *J* = 4.4 Hz, 4H), 2.60 (t, *J* = 7.5 Hz, 2H), 2.07 – 1.96 (m, 4H), 1.74 (p, *J* = 7.5 Hz, 2H), 1.48 – 1.32 (m, 2H), 1.36 – 1.21 (m, 18H), 0.87 (t, *J* = 7.1 Hz, 3H). ^13^C NMR (101 MHz, CDCl_3_) δ 169.31, 168.81, 130.18, 129.84, 32.04, 31.08, 29.91, 29.78, 29.67, 29.46, 29.14, 28.91, 27.36, 27.29, 25.73, 24.70, 22.83, 14.26.

### 2,5-Dioxopyrrolidin-1-yl (5Z,8Z,11Z,14Z)-eicosa-5,8,11,14-tetraenoate (3)

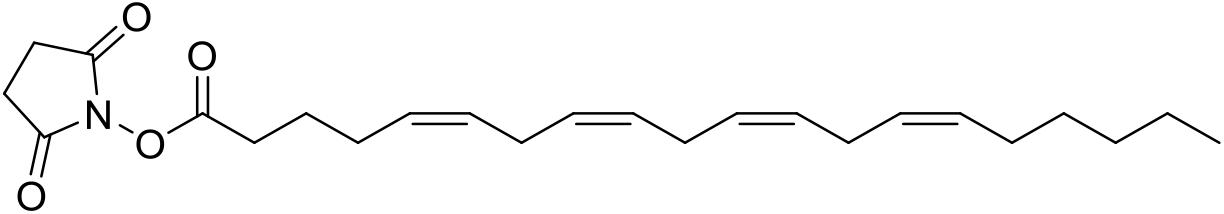

(5Z,8Z,11Z,14Z)-Eicosa-5,8,11,14- tetraenoic acid (374 mg, 1.23 mmol) was dissolved in dry DCM (31 mL). NHS (212 mg, 1.84 mmol), EDC (353 mg, 1.84 mmol) and DIPEA (322 μL, 1.84 mmol) were added and the mixture was stirred overnight at room temperature. The mixture was diluted with brine (20 mL) and extracted with DCM (3×50 mL). The combined organic layers were dried (Na_2_SO_4_), filtered, and concentrated under reduced pressure to afford the title compound as an orange oil (0.493 g, 1.23 mmol, 100%). R*_f_* = 0.8 (1:1 EtOAc/Pentane). ^1^H NMR (500 MHz, CDCl_3_) δ 5.47 – 5.30 (m, 8H), 2.93 – 2.77 (m, 10H), 2.62 (t, *J* = 7.5 Hz, 2H), 2.20 (q, *J* = 7.3 Hz, 2H), 2.05 (q, *J* = 7.1 Hz, 2H), 1.83 (p, *J* = 7.5 Hz, 2H), 1.42 – 1.18 (m, 6H), 0.89 (t, *J* = 6.9 Hz, 3H). ^13^C NMR (126 MHz, CDCl_3_) δ 176.11, 169.25, 168.67, 130.65, 129.77, 128.72, 128.48, 128.28, 128.18, 128.04, 127.71, 31.67, 30.46, 29.48, 27.37, 26.32, 25.74, 24.61, 14.22.

### 2’-Bromoethyl 2,3,4,6-tetra-*O*-acetyl-β-D-galactopyranoside (4)

⍰To a cooled (0 °C) mixture of 1,2,3,4,6-penta-*O*-acetyl-D-galactopyranoside (3.9 g, 10 mmol) and 2-bromoethanol (0.92 mL, 13 mmol) in dry DCM (25 mL) was added BF_3_.OEt_2_ (3.3 mL, 26 mmol) over a period of 15 minutes. Stirring was continued for 2 hours at 0 °C and the reaction was subsequently stirred overnight at room temperature. The solution was poured onto ice H_2_O (80 mL) and extracted with DCM (3×100 mL). The combined organic layers were washed with saturated NaHCO_3_ (3×80 mL), H_2_O (2×80 mL), brine (3×80 mL), dried (MgSO_4_), filtered and concentrated under reduced pressure. Purification of the residue by column chromatography (50→100% EtOAc in pentane) afforded title compound (3.42 g, 7.51 mmol, 75%) as a colorless oil. R*_f_* = 0.4 (1:9 EtOAc/DCM). ^1^H NMR (400 MHz, CDCl_3_) δ 5.43 – 5.37 (m, 1H, H-4), 5.23 (t, *J* = 9.1 Hz, 1H, H-2), 5.03 (dd, *J* = 10.5, 4.6 Hz, 1H, H-3), 4.53 (d, *J* = 7.9 Hz, 1H, H-1), 4.23 – 4.07 (m, 3H, H_2_-6, H-7a), 3.92 (t, *J* = 6.7 Hz, 1H, H-5), 3.87 – 3.76 (m, 1H, H-7b), 3.47 (m, 2H, H_2_-8), 2.15 (s, 3H, CH_3_), 2.09 (s, 3H, CH_3_), 2.05 (s, 3H, CH_3_), 1.99 (s, 3H, CH_3_). ^13^C NMR (101 MHz, CDCl_3_) δ 170.55, 170.37, 170.30, 169.69 (4xC=O), 101.69 (C-1), 70.95, 70.88 (C-3, C-5), 69.91 (C-7), 68.67 (C-2), 67.08 (C-4), 61.39 (C-6), 30.10 (C- 8), 21.02, 20.84, 20.82, 20.74 (4x CH_3_). HRMS calculated for C_16_H_23_BrO_10_, [M+NH_4_]^+^ 473.08458; found, 473.08418.

### 2’-Azidoethyl 2,3,4,6-tetra-*O*-acetyl- β-D-galactopyranoside (5)

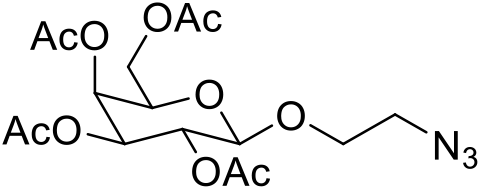

The title compound was synthesized from compound **4** following general procedure A on a 7.5 mmol scale. It was purified by column chromatography (0→50% EtOAc in toluene) to afford title compound as a colorless oil (3.14 g, 7.5 mmol, 100%). R*_f_* = 0.6 (1:1 EtOAc/toluene). ^1^H NMR (500 MHz, CDCl_3_) δ 5.40 (dd, *J* = 3.5, 1.2 Hz, 1H, H- 4), 5.25 (dd, *J* = 10.5, 8.0 Hz, 1H, H-2), 5.03 (dd, *J* = 10.5, 3.4 Hz, 1H, H-3), 4.56 (d, *J* = 8.0 Hz, 1H, H-1), 4.19 (dd, *J* = 11.3, 6.6 Hz, 1H, H-6a), 4.13 (dd, *J* = 11.3, 6.8 Hz, 1H, H-6b), 4.07 – 4.02 (m, 1H, H-7a), 3.92 (td, *J* = 6.7, 1.2 Hz, 1H, H-5), 3.67 (m, 1H, H-7b), 3.57 – 3.47 (m, 1H, H-8a), 3.36 – 3.27 (m, 1H, H-8b), 2.16 (s, 3H, CH_3_), 2.07 (s, 3H, CH_3_), 2.05 (s, 3H, CH_3_), 1.99 (s, 3H, CH_3_).^13^C NMR (126 MHz, CDCl_3_) δ 170.46, 170.30, 170.22, 169.55 (4xC=O), 101.25 (C-1), 71.00 (C-5), 70.93 (C-3), 68.64 (C-2), 68.45 (C-7), 67.10 (C-4), 61.34 (C-6), 50.67 (C-8), 20.96, 20.75, 20.74, 20.65 (4x CH_3_). HRMS calculated for C_16_H_23_N_3_O_10_, [M+Na]^+^ 440.12756; found, 440.12714.

### 2’-Palmitamidoethyl 2,3,4,6-tetra-*O*-acetyl-β-D-galactopyranoside (6)

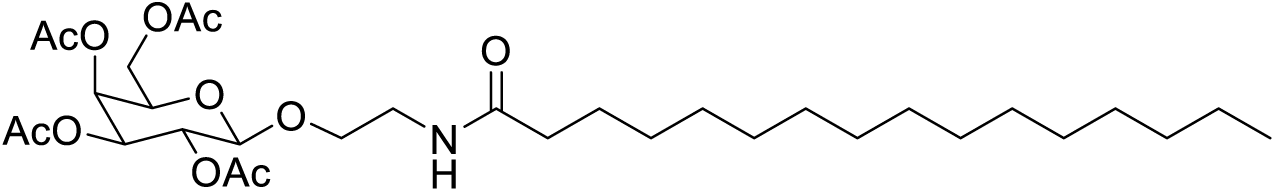

The title compound was synthesized from compound **5** following general procedure B on a 0.12 mmol scale and purified to afford title compound as a colorless oil (35.4 mg, 0.056 mmol, 35%). R*_f_* = 0.4 (3:1 EtOAc/pentane). ^1^H NMR (500 MHz, CDCl_3_) δ 5.88 (bs, 1H, NH), 5.39 (dd, *J* = 3.4, 1.2 Hz, 1H, H-4), 5.18 (dd, *J* = 10.5, 7.9 Hz, 1H, H-2), 5.01 (dd, *J* = 10.5, 3.4 Hz, 1H, H-3), 4.46 (d, *J* = 7.9 Hz, 1H, H-1), 4.19 – 4.09 (m, 2H, H_2_-6), 3.91 (td, *J* = 6.5, 1.0 Hz, 1H, H-5), 3.89 – 3.84 (m, 1H, H-7a), 3.70 – 3.65 (m, 1H, H-7b), 3.56 – 3.38 (m, 2H, H_2_-8), 2.17 (t, *J* = 7.1 Hz, 2H, H_2_-9), 2.15 (s, 3H, CH_3_), 2.06 (s, 3H, CH_3_), 2.04 (s, 3H, CH_3_), 1.98 (s, 3H, CH_3_), 1.61 (t, *J* = 7.4 Hz, 2H, H_2_-10), 1.24 (s, 24H, H_2_-11 to H_2_-22), 0.87 (t, *J* = 6.9 Hz, 3H, H_3_-23). ^13^C NMR (126 MHz, CDCl_3_) δ 173.42, 170.50, 170.29, 170.20, 169.77 (5xC=O), 101.56 (C-1), 70.98 (C-5), 70.85 (C-3), 69.33 (C-7), 69.07 (C-2), 67.09 (C-4), 61.43 (C-6), 39.22 (C-8), 36.84 (C-9), 32.05, 29.82, 29.78, 29.64, 29.51, 29.49, 29.49, 25.83, 22.82 (13xCH_2_, C-10 to C-22), 20.96, 20.79, 20.78, 20.70 (4xCH_3_), 14.25 (C-23). HRMS calculated for C_32_H_55_NO_11_, [M+H]^+^ 630.38479; found, 630.38436.

### 2’-Oleamidoethyl 2,3,4,6-tetra-*O*-acetyl-β-D-galactopyranoside (7)

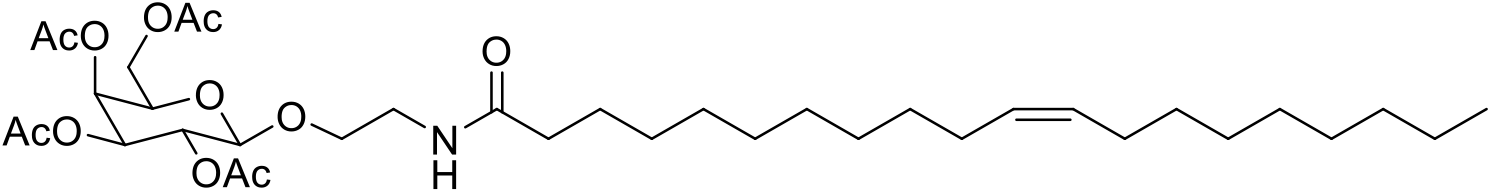

The title compound was synthesized from compound **5** following general procedure B on a 0.24 mmol scale and purified to afford the title compound as a colorless oil (67.8 mg, 0.103 mmol, 43%). R*_f_* = 0.5 (3:1 EtOAc/pentane). ^1^H NMR (400 MHz, CDCl_3_) δ 5.90 (t, *J* = 5.7 Hz, 1H, NH), 5.36 (dd, *J* = 3.5, 1.1 Hz, 1H, H-4), 5.35 – 5.26 (m, 2H, H-16, H-17), 5.16 (dd, *J* = 10.5, 7.9 Hz, 1H, H- 2), 4.99 (dd, *J* = 10.5, 3.4 Hz, 1H, H-3), 4.45 (d, *J* = 7.9 Hz, 1H, H-1), 4.18 – 4.06 (m, 2H, H_2_-6), 3.90 (td, *J* = 6.6, 1.2 Hz, 1H, H-5), 3.87 – 3.81 (m, 1H, H-7a), 3.70 – 3.61 (m, 1H, H-7b), 3.52 – 3.34 (m, 2H, H_2_-8), 2.16 – 2.10 (m, 5H, CH_3_, H_2_-9), 2.03 (s, 3H, CH_3_), 2.02 (s, 3H, CH_3_), 1.96 (s, 3H, CH_3_), 1.59 (t, *J* = 7.4 Hz, 2H, H_2_-10), 1.30 – 1.20 (m, 20H, H_2_-11 to H_2_-14, H_2_-19 to H_2_-24), 0.87 – 0.82 (t, *J* = 7.1 Hz, 3H, H_3_-25). ^13^C NMR (101 MHz, CDCl_3_) δ 173.29, 170.42, 170.23, 170.12, 169.70 (5xC=O), 130.04, 129.79 (C-16, C-17), 101.47 (C-1), 70.89 (C-3), 70.78 (C-5), 69.25 (C-7), 68.99 (C-2), 67.03 (C-4), 61.36 (C-6), 39.13 (C-8), 36.75 (C-9), 31.95, 29.81, 29.78, 29.57, 29.39, 29.37, 29.21, 27.27, 27.24, 25.73, 22.73 (13xCH_2_, C-10 to C-15, C-18 to C-24), 20.89, 20.72, 20.71, 20.62 (4xCH_3_), 14.18 (C-25). HRMS calculated for C_34_H_57_NO_11_, [M+Na]^+^ 678.38238; found, 678.38213.

### 2’-(5Z,8Z,11Z,14Z)-Icosamidoethyl 2,3,4,6-tetra-*O*-acetyl-β-D-galactopyranoside (8)

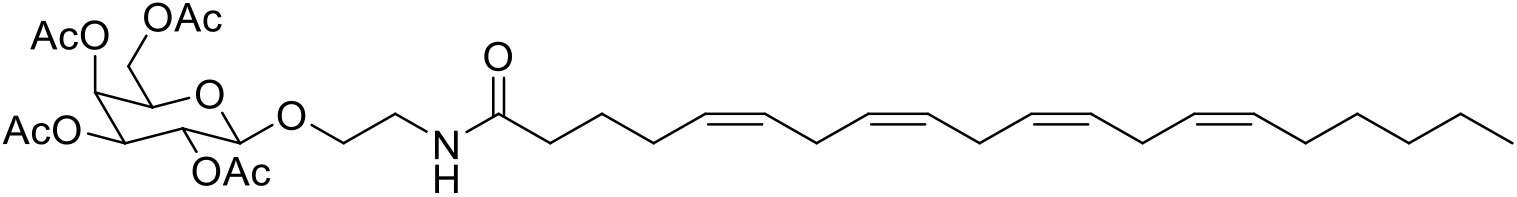

The title compound was synthesized from compound **5** following general procedure B on a 0.192 mmol scale and purified to afford title compound as a colorless oil (65.2 mg, 0.096 mmol, 50%). R*_f_* = 0.7 (3:1 EtOAc/pentane). ^1^H NMR (500 MHz, CDCl_3_) δ 5.82 (t, *J* = 6.0 Hz, 1H, NH), 5.45 – 5.27 (m, 9H, H-4, H-12, H-13, H-15, H-16, H-18, H-19, H-21, H-22), 5.19 (dd, *J* = 10.5, 7.9 Hz, 1H, H-2), 5.02 (dd, *J* = 10.5, 3.5 Hz, 1H, H-3), 4.47 (d, *J* = 7.9 Hz, 1H, H-1), 4.20 – 4.08 (m, 2H, H_2_-6), 3.94 – 3.84 (m, 2H, H-5, H-7a), 3.70 – 3.64 (m, 1H, H-7b), 3.54 – 3.38 (m, 2H, H_2_-8), 2.88 – 2.78 (m, 6H, H_2_-14, H_2_- 17, H_2_-20), 2.21 – 2.17 (m, 2H, H_2_-9), 2.16 (s, 3H, CH_3_), 2.08 – 2.04 (m, 6H, 2x CH_3_), 1.99 (s, 3H, CH_3_), 1.72 (p, *J* = 7.7 Hz, 2H, H_2_-10), 1.40 – 1.23 (m, 10H, H_2_-11, H_2_-23, H_2_-24, H_2_-25, H_2_-26), 0.89 (t, *J* = 6.9 Hz, 3H, H_3_-27). ^13^C NMR (126 MHz, CDCl_3_) δ 172.97, 170.51, 170.31, 170.22, 169.77 (5xC=O), 130.67, 129.24, 128.94, 128.76, 128.40, 128.31, 128.00, 127.68 (C-12, C-13, C-15, C-16, C-18, C-19, C-21, C-22), 101.59 (C-1), 71.00 (C-3), 70.88 (C-5), 69.32 (C-7), 69.08 (C-2), 67.10 (C-4), 61.43 (C-6), 39.23 (C-8), 36.16 (C-9), 31.66, 29.47, 27.37, 26.86, 25.79, 25.78, 25.78, 25.62, 22.72 (9xCH_2_, C-10, C-11, C-14, C-17, C-20, C-23, C-24, C-25, C-26), 21.00, 20.82, 20.81, 20.72 (4xCH_3_), 14.23 (C-27). HRMS calculated for C_36_H_55_NO_11_, [M+Na]^+^ 700.36673; found, 700.36704.

### 2’-Palmitamidoethyl β-D-galactopyranoside (9)

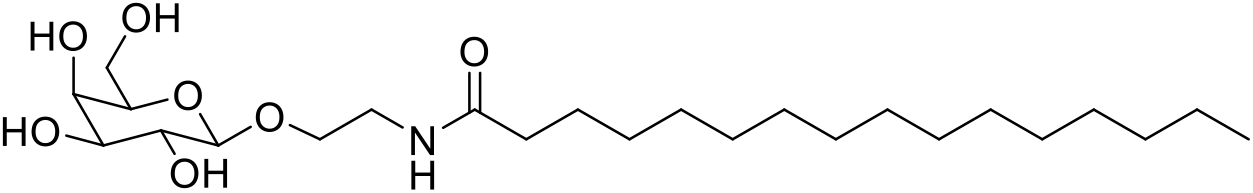

The title compound was synthesized from compound **6** on a 0.056 mmol scale following general procedure C and purified to afford compound title compound as a white solid (23.5 mg, 0.051 mmol, 91%). R*_f_* = 0.3 (1:9 MeOH/DCM). ^1^H NMR (600 MHz, methanol-d_4_) δ 4.23 (d, *J* = 7.6 Hz, 1H, H-1), 3.95 – 3.89 (m, 1H, H-7a), 3.82 (dd, *J* = 3.4, 1.1 Hz, 1H, H-4), 3.76 (dd, *J* = 11.3, 7.0 Hz, 1H, H-6a), 3.72 (dd, *J* = 11.3, 5.2 Hz, 1H, H-6b), 3.69 – 3.61 (m, 1H, H-7b), 3.56 – 3.50 (m, 2H, H-2, H-5), 3.50 – 3.44 (m, 2H, H-3, H-8a), 3.38 – 3.33 (m, 1H, H-8b), 2.20 (t, *J* = 7.9 Hz, 2H, H_2_-9), 1.60 (q, *J* = 7.4 Hz, 2H, H_2_-10), 1.31 (bs, 24H, H_2_- 11 to H_2_-22), 0.90 (t, *J* = 7.0 Hz, 3H, H_3_-23). ^13^C NMR (151 MHz, methanol-d_4_) δ 176.46 (C=O), 105.13 (C-1), 76.77 (C-5), 74.88 (C-3), 72.57 (C-2), 70.27 (C-4), 69.64 (C-7), 62.53 (C-6), 40.62 (C-8), 37.12 (C-9), 33.09, 30.82, 30.81, 30.79, 30.78, 30.77, 30.66, 30.50, 30.38, 27.04, 23.76 (13xCH_2_, C-10 to C-22), 14.46 (C-23). HRMS calculated for C_24_H_47_NO_7_, [M+Na]^+^ 484.32447; found, 484.32414.

### 2’-Oleamidoethyl β-D-galactopyranoside (10)

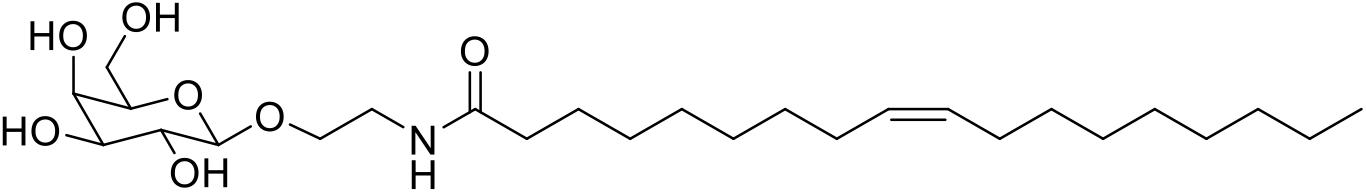

The title compound was synthesized from compound **7** on a 0.103 mmol scale following general procedure C and purified to afford title compound as a white solid (50.2 mg, 0.103 mmol, 100%). R*_f_* = 0.4 (1:9 MeOH/DCM). ^1^H NMR (500 MHz, methanol-d_4_) δ 5.39 – 5.31 (m, 2H, H-16, H-17), 4.23 (d, *J* = 7.5 Hz, 1H, H-1), 3.95 – 3.88 (m, 1H, H-7a), 3.83 (dd, *J* = 3.3, 1.1 Hz, 1H, H-4), 3.79 – 3.69 (m, 2H, H_2_-6), 3.69 – 3.63 (m, 1H, H-7b), 3.57 – 3.50 (m, 2H, H-2, H-5), 3.50 – 3.44 (m, 2H, H-3, H-8a), 3.39 – 3.32 (m, 1H, H-8b), 2.20 (t, *J* = 7.9 Hz, 2H, H_2_-9), 2.07 – 1.99 (m, 4H, H_2_-15, H_2_-18), 1.60 (p, *J* = 7.5 Hz, 2H, H_2_-10), 1.32 (m, 22H, H_2_-11 to H_2_-14, H_2_-18 to H_2_-24), 0.93 – 0.88 (t, *J* = 7.2 Hz, 3H, H_3_-25). ^13^C NMR (126 MHz, methanol-d_4_) δ 176.39 (C=O), 130.86, 130.79 (C-16, C-17), 105.11 (C-1), 76.74 (C-5), 74.87 (C-3), 72.56 (C-2), 70.26 (C-4), 69.63 (C-7), 62.51 (C-6), 40.61 (C-8), 37.10 (C-9), 33.06, 30.85, 30.85, 30.61, 30.45, 30.39, 30.37, 30.34, 30.25, 28.15, 28.13, 27.02, 23.74 (13xCH_2_, C-10 to C-15, C-18 to C-24), 14.47 (C-25). HRMS calculated for C_26_H_49_NO_7_, [M+H]^+^ 488.35818; found, 488.35815.

### 2’-(5Z,8Z,11Z,14Z)-Icosamidoethyl β-D-galactopyranoside (11)

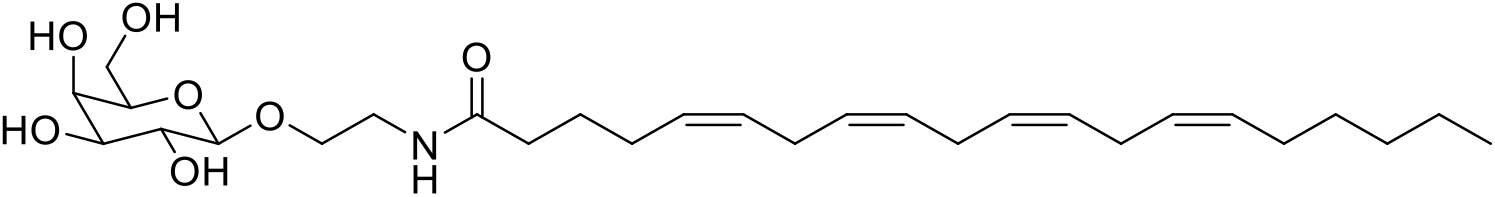

The title compound was synthesized from compound **8** on a 0.096 mmol scale following general procedure C and purified to afford compound title compound as a white solid (39.8 mg, 0.078 mmol, 81%). R*_f_* = 0.3 (1:9 MeOH/DCM). ^1^H NMR (500 MHz, methanol-d_4_) δ 5.43 – 5.31 (m, 8H, H-12, H-13, H-15, H-16, H-18, H-19, H-21, H-22), 4.23 (d, *J* = 7.5 Hz, 1H, H-1), 3.96 – 3.88 (m, 1H, H-7a), 3.83 (dd, *J* = 3.3, 1.1 Hz, 1H, H-4), 3.80 – 3.69 (m, 2H, H_2_-6), 3.69 – 3.62 (m, 1H, H-7b), 3.57 – 3.50 (m, 2H, H-2, H-5), 3.49 – 3.43 (m, 2H, H-3, H-8a), 3.38 – 3.32 (m, 1H, H-8b), 2.91 – 2.77 (m, 6H, H_2_-14, H_2_-17, H_2_-20), 2.25 – 2.19 (m, 2H, H_2_-9), 2.17 – 2.04 (m, 4H, H_2_-11, H_2_-23), 1.73 – 1.64 (m, 2H, H_2_-10), 1.43 – 1.28 (m, 6H, H_2_-24, H_2_-25, H_2_-26), 0.91 (t, *J* = 6.9 Hz, 3H, H_3_-27). ^13^C NMR (126 MHz, methanol-d_4_) δ 176.11 (C=O), 131.21, 130.20, 129.73, 129.47, 129.22, 129.15, 128.91, 128.77 (C-12, C-13, C-15, C-16, C-18, C-19, C-21, C-22), 105.11 (C-1), 76.76 (C-5), 74.90 (C-3), 72.58 (C-2), 70.27 (C-4), 69.61 (C-7), 62.52 (C-6), 40.63 (C-8), 36.56 (C-9), 32.66, 30.75, 30.47, 28.18, 27.76, 26.94, 26.56, 26.55, 23.63 (9xCH_2_, C-10, C-11, C-23, C-24, C-25, C-26), 14.44 (C-27). HRMS calculated for C_28_H_47_NO_7_, [M+Na]^+^ 532.32447; found, 532.32397.

### **⍰**2’-Bromoethyl 2,3,4,6-tetra-*O*-acetyl-β-D-glucopyranoside (12)

To a cooled (0 °C) mixture of 1,2,3,4,6-penta-O-acetyl-β-D-glucopyranose (1.00 g, 2.56 mmol) and 2-bromoethanol (0.235 mL, 3.33 mmol) in dry DCM (6.5 mL) was over a period of 15 minutes added BF_3_.OEt_2_ (0.844 mL, 6.66 mmol), stirring was continued for 2 hours at 0 °C and stirred overnight at room temperature. The solution was poured onto ice H_2_O (30 mL) and extracted with DCM (3×50 mL). The combined organic layers were washed with NaHCO_3_ (3×30 mL), H_2_O (2×30 mL), brine (3×30 mL), dried (MgSO_4_), filtered and concentrated. Purification of the residue by column chromatography (50→100 Et_2_O in pentane) afforded title compound (0.594 g, 1.31 mmol, 51%) as a colorless oil. R*_f_* = 0.6 (3:1 Et_2_O/pentane). ^1^H NMR (400 MHz, CDCl_3_) δ 5.22 (t, *J* = 9.5 Hz, 1H, H-3), 5.08 (dd, *J* = 10.0, 9.3 Hz, 1H, H-4), 5.02 (dd, *J* = 9.7, 7.9 Hz, 1H, H-2), 4.57 (d, *J* = 8.0 Hz, 1H, H-1), 4.26 (dd, *J* = 12.3, 4.8 Hz, 1H, H-6a), 4.20 – 4.11 (m, 2H, H-6b, H-7a), 3.86 – 3.77 (m, 1H, H-7b), 3.75 – 3.67 (m, 1H, H-5), 3.51 – 3.41 (m, 2H, H_2_-8), 2.09 (s, 3H, CH_3_), 2.07 (s, 3H, CH_3_), 2.03 (s, 3H, CH_3_), 2.01 (s, 3H, CH_3_). ^13^C NMR (101 MHz, CDCl_3_) δ 170.79, 170.40, 169.56, 169.54 (4xC=O), 101.17 (C-1), 72.73 (C-3), 72.07 (C-5), 71.14 (C-2), 69.93 (C-7), 68.44 (C-4), 61.97 (C-6), 30.01 (C-8), 20.89, 20.89, 20.76, 20.74 (4xCH_3_). HRMS calculated for C_16_H_23_BrO_10_, [M+NH_4_]^+^ 473.08458; found, 473.08441.

### 2’-Azidoethyl 2,3,4,6-tetra-*O*-acetyl-β-D-glucopyranoside (13)

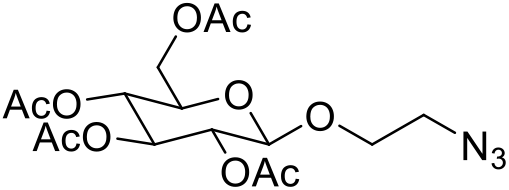

The title compound was synthesized from compound **12** following general procedure A on a 1.2 mmol scale. It was dissolved in heptane (3×20 mL) and concentrated under reduced pressure to afford title compound as colorless crystals (0.425 g, 1.02 mmol, 87%). R*_f_* = 0.6 (1:1 EtOAc/toluene). ^1^H NMR (500 MHz, CDCl_3_) δ 5.22 (t, *J* = 9.5 Hz, 1H, H-3), 5.10 (t, *J* = 9.9 Hz, 1H, H-4), 5.03 (dd, *J* = 9.6, 8.0 Hz, 1H, H-2), 4.60 (d, *J* = 8.0 Hz, 1H, H-1), 4.26 (dd, *J* = 12.3, 4.7 Hz, 1H, H-6a), 4.16 (dd, *J* = 12.3, 2.4 Hz, 1H, H-6b), 4.09 – 3.99 (m, 1H, H-7a), 3.76 – 3.66 (m, 2H, H-5, H-7b), 3.54 – 3.46 (m, 1H, H-8a), 3.34 – 3.26 (m, 1H, H-8b), 2.09 (s, 3H, CH_3_), 2.06 (s, 3H, CH_3_), 2.03 (s, 3H, CH_3_), 2.01 (s, 3H, CH_3_).^13^C NMR (126 MHz, CDCl_3_) δ 170.81, 170.43, 169.56, 168.09 (4xC=O), 100.82 (C-1), 72.93 (C-3), 72.09 (C-5), 71.20 (C-2), 68.71 (C-7), 68.45 (C-4), 61.97 (C-6), 50.67 (C-8), 20.90, 20.84, 20.76, 20.76 (4xCH_3_). HRMS calculated for C_16_H_23_N_3_O_10_, [M+NH_4_]^+^ 435.17217; found, 435.17158.

### 2’-Palmitamidoethyl 2,3,4,6-tetra-*O*-acetyl-β-D-glucopyranoside (14)

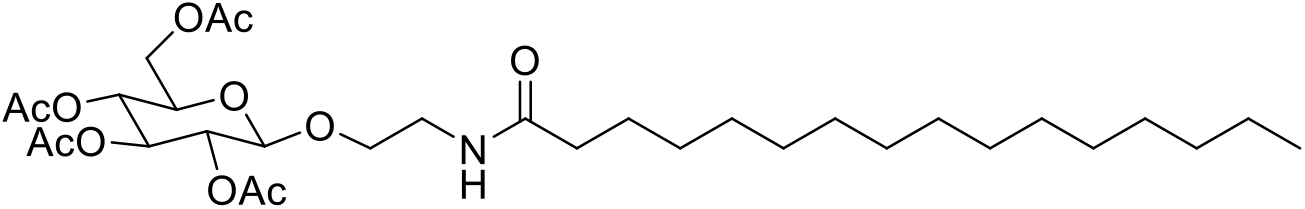

The title compound was synthesized from compound 30 following general procedure B on a 0.12 mmol scale and purified to afford title compound as a white solid (30.8 mg, 0.049 mmol, 41%). R*_f_* = 0.6 (3:1 EtOAc/pentane). ^1^H NMR (400 MHz, CDCl_3_) δ 5.88 (t, *J* = 5.6 Hz, 1H, NH), 5.20 (t, *J* = 9.5 Hz, 1H, H-3), 5.07 (t, *J* = 9.7 Hz, 1H, H-4), 4.98 (dd, *J* = 9.7, 7.9 Hz, 1H, H-2), 4.50 (d, *J* = 7.9 Hz, 1H, H-1), 4.25 (dd, *J* = 12.4, 4.9 Hz, 1H, H-6a), 4.14 (dd, *J* = 12.3, 2.4 Hz, 1H, H-6b), 3.87 – 3.79 (m, 1H, H-7a), 3.74– 3.66 (m, 2H, H-5, H-7b), 3.52 – 3.35 (m, 2H, H_2_-8), 2.15 (dd, *J* = 8.5, 6.9 Hz, 2H, H_2_-9), 2.08 (s, 3H, CH_3_), (5xC=O), 101.09 (C-1), 72.74 (C-3), 72.08 (C-5), 71.47 (C-2), 69.49 (C-7), 68.37 (C-4), 61.94 (C-6), 39.21 (C- 8), 36.83 (C-9), 32.05, 29.82, 29.78, 29.76, 29.64, 29.51, 29.49, 25.83, 22.82 (13xCH_2_, C-10 to C-22), 20.86, 20.86, 20.73, 20.72 (4xCH_3_), 14.26 (C-23). HRMS calculated for C_32_H_55_NO_11_, [M+H]^+^ 630.38479; found, 630.38429.

### 2’-Oleamidoethyl 2,3,4,6-tetra-*O*-acetyl-β-D-glucopyranoside (15)

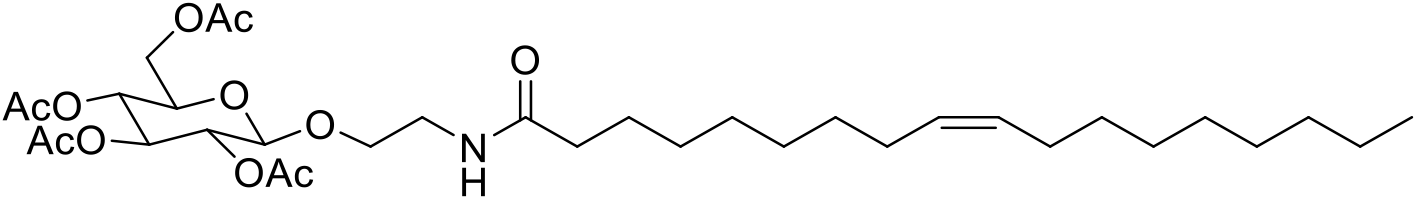

The title compound was synthesized from compound **13** following general procedure B on a 0.24 mmol scale and purified to afford title compound as a colorless oil (57.3 mg, 0.087 mmol, 36%). R*_f_* = 0.5 (3:1 EtOAc/pentane). ^1^H NMR (400 MHz, CDCl_3_) δ 5.91 (t, *J* = 5.7 Hz, 1H, NH), 5.38 – 5.27 (m, 2H, H-16, H-17), 5.19 (t, *J* = 9.5 Hz, 1H, H-3), 5.06 (t, *J* = 9.7 Hz, 1H, H-4), 4.97 (dd, *J* = 9.6, 7.9 Hz, 1H, H-2), 4.49 (d, *J* = 7.9 Hz, 1H, H-1), 4.24 (dd, *J* = 12.3, 4.9 Hz, 1H, H-6a), 4.13 (dd, *J* = 12.3, 2.4 Hz, 1H, H-6b), 3.87 – 3.79 (m, 1H, H-7a), 3.75 – 3.64 (m, 2H, H-5, H-7b), 3.52 – 3.38 (m, 2H, H_2_-8), 2.15 (dd, *J* = 8.5, 6.9 Hz, 2H, H_2_-9), 2.07 (s, 3H, CH_3_), 2.03 (s, 3H, CH_3_), 2.01 (s, 3H, CH_3_), 1.99 (s, 3H, CH_3_), 1.64 – 1.56 (m, 2H, H_2_-10), 1.27 (d, 24H, H_2_-11 to H_2_-15, H_2_-18 to H_2_-24), 0.82 (t, *J* = 6.8 Hz, 3H, H_3_-25). ^13^C NMR (101 MHz, CDCl_3_) δ 173.39, 170.70, 170.28, 169.55,169.53 (5xC=O), 130.09, 129.85 (2xCH, C-16, C-17), 101.06 (C-1), 72.71 (C-3), 72.05 (C-5), 71.44 (C-2), 69.43 (C-7), 68.35 (C-4), 61.91 (C-6), 39.21 (C-8), 36.75 (C-9), 32.00, 29.86, 29.83, 29.62, 29.43, 29.42, 29.40, 29.27, 27.32, 25.79, 22.78 (13xCH_2_, C-10 to C-15, C-18 to C-24), 20.83, 20.70 (4xCH_3_), 14.23 (C-25). HRMS calculated for C_34_H_57_NO_11_, [M+H]^+^ 656.40044; found, 656.39994.

### 2’-(5Z,8Z,11Z,14Z)-Icosamidoethyl 2,3,4,6-tetra-*O*-acetyl-β-D-glucopyranoside (16)

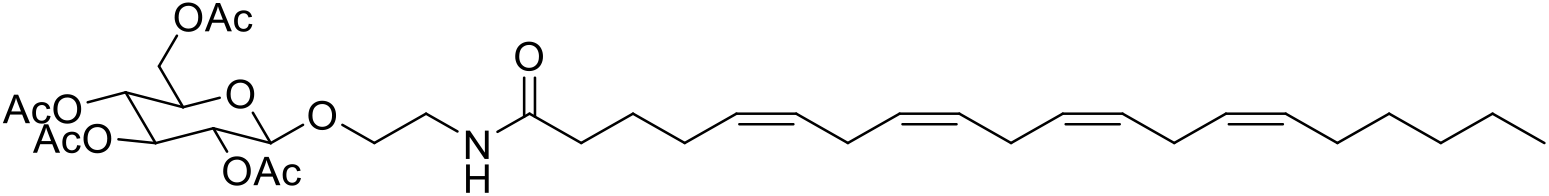

The title compound was synthesized from compound **13** following general procedure B on a 0.192 mmol scale and purified to afford title compound as a colorless oil (81.3 mg, 0.120 mmol, 63%). R*_f_* = 0.6 (3:1 EtOAc/pentane). ^1^H NMR (400 MHz, CDCl_3_) δ 5.85 (t, *J* = 5.5 Hz, 1H, NH), 5.46 – 5.30 (m, 8H, H-12, H-13, H-15, H-16, H-18, H-19, H-21, H-22), 5.21 (t, *J* = 9.5 Hz, 1H, H-3), 5.07 (t, *J* = 9.7 Hz, 1H, H-4), 4.99 (dd, *J* = 9.7, 7.9 Hz, 1H, H-2), 4.50 (d, *J* = 7.9 Hz, 1H, H-1), 4.26 (dd, *J* = 12.4, 4.9 Hz, 1H, H-6a), 4.15 (dd, *J* = 12.4, 2.5 Hz, 1H, H-6b), 3.90 – 3.79 (m, 1H, H- 7a), 3.76 – 3.64 (m, 2H, H-5, H-7b), 3.52 – 3.39 (m, 2H, H_2_-8), 2.89 – 2.76 (m, 6H, H_2_-14, H_2_-17, H_2_-20), 2.18 (dd, *J* = 8.6, 6.7 Hz, 2H, H_2_-9), 2.09 (s, 3H, CH_3_), 2.05 (s, 3H, CH_3_), 2.03 (s, 3H, CH_3_), 2.01 (s, 3H, CH_3_), 1.72 (q, *J* = 7.5 Hz, 2H, H_2_-10), 1.39 – 1.23 (m, 10H, H_2_-11, H_2_-23 to H_2_-26), 0.88 (t, *J* = 6.8 Hz, 3H, H_3_-27). ^13^C NMR (101 MHz, CDCl_3_) δ 173.00, 171.13, 170.74, 170.34, 169.56 (5xC=O), 130.66, 129.23, 128.94, 128.76, 128.40, 128.32, 128.02, 127.69 (C-12, C-13, C-15, C-16, C-18, C-19, C-21, C-22), 101.12 (C-1), 72.78 (C-3), 72.13 (C-5), 71.50 (C-2), 69.46 (C-7), 68.43 (C-4), 61.97 (C-6), 39.25 (C-8), 36.13 (C-9), 31.66, 29.47, 27.37, 26.85, 25.78, 25.63, 22.72 (9xCH_2_, C-10, C-11, C-14, C-17, C-20, C-23 to C-26), 20.88, 20.74 (4x CH_3_), 14.22 (C-27). HRMS calculated for C_36_H_55_NO_11_, [M+Na]^+^ 700.36673; found, 700.36628.

### 2’-Palmitamidoethyl β-D-glucopyranoside (17)

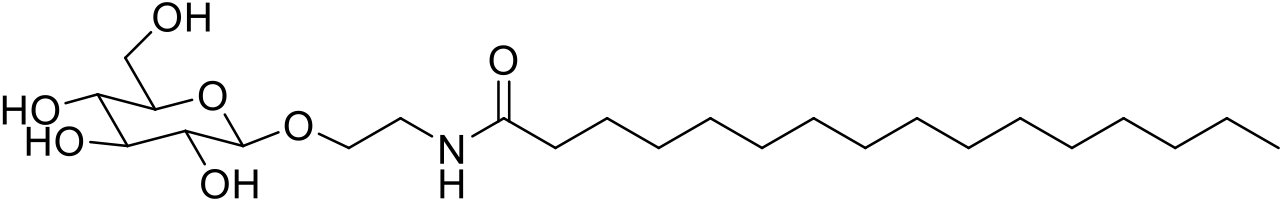

The title compound was synthesized from compound on a 0.049 mmol scale following general procedure C and purified to afford title compound as a white solid (20.2 mg, 0.044 mmol, 89%). R*_f_* = 0.4 (1:9 MeOH/DCM). ^1^H NMR (500 MHz, methanol-d_4_) δ 4.27 (d, *J* = 7.8 Hz, 1H, H-1), 3.94 – 3.89 (m, 1H, H-7a), 3.87 (dd, *J* = 11.8, 1.6 Hz, 1H, H-6a), 3.68 – 3.62 (m, 2H, H-6b, H-7b), 3.50 – 3.44 (m, 1H, H- 8a), 3.38 – 3.32 (m, 2H, H-3, H-8b), 3.29 – 3.26 (m, 2H, H-4, H-5), 3.19 (dd, *J* = 9.2, 7.8 Hz, 1H, H-2), 2.20 (dd, *J* = 8.3, 6.9 Hz, 2H, H_2_-9), 1.60 (q, *J* = 7.1 Hz, 2H, H_2_-10), 1.329 (bs, 24H, H_2_-11 to H_2_-22), 0.90 (t, *J* = 7.4 Hz, 3H, H_3_-23). ^13^C NMR (126 MHz, methanol-d_4_) δ 176.49 (C=O), 104.53 (C-1), 78.01, 77.96 (C-3, C-5), 75.12 (C-2), 71.61(C-4), 69.69 (C-7), 62.71 (C-6), 40.58 (C-8), 37.12 (C-9), 33.09, 31.63, 30.81, 30.78, 30.76, 30.66, 30.49, 30.35, 30.15, 27.04, 24.95, 24.05 (13xCH_2_, C-10 to C-22), 14.45 (C-23). HRMS calculated for C_24_H_47_NO_7_, [M+H]^+^ 462.34253; found, 462.34254.

### 2’-Oleamidoethyl β-D-glucopyranoside (18)

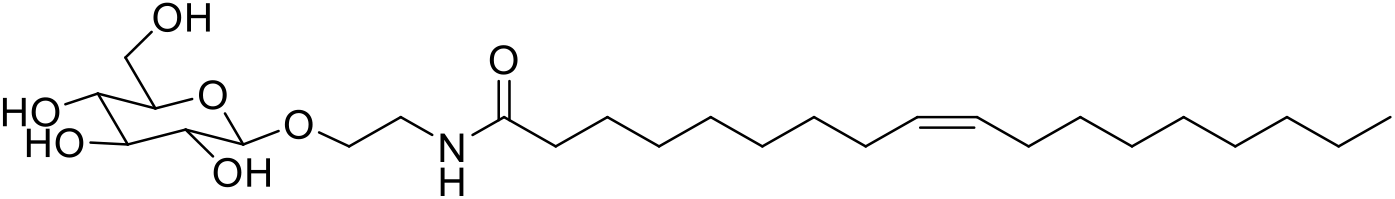

The title compound was synthesized from compound **15** on a 0.087 mmol scale following general procedure C and purified to afford title compound as a white solid (35.1 mg, 0.072 mmol, 82%). R*_f_* = 0.6 (1:9 MeOH/DCM). ^1^H NMR (500 MHz, methanol-d_4_) δ 5.39 – 5.30 (m, 2H, H-16, H-17), 4.27 (d, *J* = 7.8 Hz, 1H, H-1), 3.94 – 3.89 (m, 1H, H-7a), 3.87 (dd, *J* = 11.8, 1.8 Hz, 1H, H-6a), 3.70 – 3.61 (m, 2H, H-6b, H-7b), 3.52 – 3.43 (m, 1H, H-8a), 3.38 – 3.32 (m, 2H, H-3, H-8b), 3.29 – 3.26 (m, 2H, H-4, H-5), 3.19 (dd, *J* = 9.2, 7.8 Hz, 1H, H-2), 2.20 (t, *J* = 7.7 Hz, 2H, H_2_-9), 2.03 (dt, *J* = 7.5, 5.4 Hz, 4H, H_2_-15, H_2_-18), 1.61 (q, *J* = 7.2 Hz, 2H, H_2_-10), 1.31 (d, *J* = 17.8 Hz, 20H, H_2_-11 to H_2_-14, H_2_-19 to H_2_-24), 0.90 (d, *J* = 7.2 Hz, 3H, H_3_-25). ^13^C NMR (126 MHz, methanol-d_4_) δ 176.45 (C=O), 130.88, 130.80 (C-16, C-17), 104.53 (C-1), 78.01, 77.97 (C-3, C-5), 75.11 (C-2), 71.62 (C-4), 69.69 (C-7), 62.72 (C-6), 40.58 (C-8), 37.11 (C-9), 33.06, 30.84, 30.61, 30.45, 30.38, 30.34, 30.25, 28.14, 27.03, 23.74 (13xCH_2_, C-10 to C-15, C-18 to C-24), 14.46 (C-25). HRMS calculated for C_26_H_49_NO_7_, [M+H]^+^ 488.35818; found, 488.35783.

### 2’-(5Z,8Z,11Z,14Z)-Icosamidoethyl 2,3,4,6-tetra-*O*-acetyl-β-D-glucopyranoside (19)

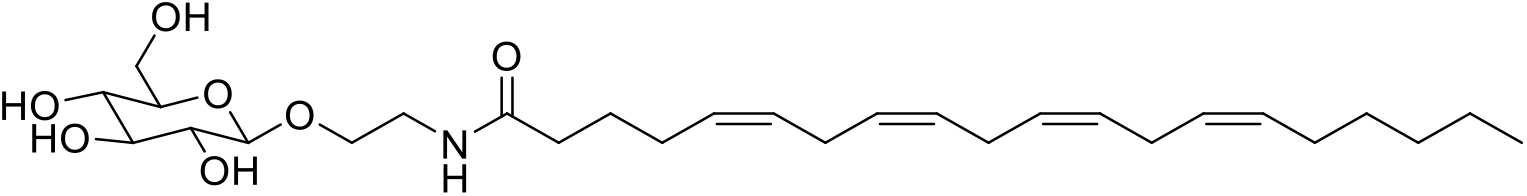

The title compound was synthesized from compound **16** on a 0.120 mmol scale following general procedure C and purified to afford title compound as a white solid (60 mg, 0.12 mmol, 98%). R*_f_* = 0.4 (1:9 MeOH/DCM). ^1^H NMR (500 MHz, methanol-d_4_) δ 5.50 – 5.25 (m, 8H, H-12, H-13, H-15, H-16, H-18, H-19, H-21, H-22), 4.27 (d, *J* = 7.8 Hz, 1H, H-1), 3.95 – 3.89 (m, 1H, H-7a), 3.87 (dd, *J* = 11.8, 1.6 Hz, 1H, H-6a), 3.69 – 3.62 (m, 2H, H-6b, H-7b), 3.50 - 3.43 (m, 1H, H-8a), 3.39 – 3.32 (m, 2H, H-3, H-8b), 3.30 – 3.26 (m, 2H, H-4, H-5), 3.20 (dd, *J* = 9.2, 7.8 Hz, 1H, H-2), 2.87 – 2.80 (m, 6H, H_2_-14, H_2_-17, H_2_-20), 2.22 (t, *J* = 7.9 Hz, 2H, H_2_-9), 2.16 – 2.04 (m, 4H, H_2_-11, H_2_-23), 1.68 (p, *J* = 7.5 Hz, 2H, H_2_-10), 1.41 – 1.27 (m, 6H, H_2_-24, H_2_-25, H_2_-26), 0.91 (t, *J* = 6.8 Hz, 3H, H_3_-27). ^13^C NMR (126 MHz, methanol-d_4_) δ 176.12 (C=O), 131.20, 130.19, 129.72, 129.46, 129.21, 129.14, 128.90, 128.76 (C-12, C-13, C-15, C-16, C-18, C-19, C-21, C-22), 104.48 (C-1), 77.98 (C-4), 77.95 (C-3), 75.09 (C-2), 71.60 (C-4), 69.64 (C-7), 62.71 (C-6), 40.59 (C-8), 36.54 (C-9), 32.65, 30.45, 28.18, 27.73, 26.92, 26.56, 23.62 (9xCH_2_, C-10, C- 11, C-14, C-17, C-20, C-23 to C-26), 14.44 (C-27). HRMS calculated for C_28_H_47_NO_7_, [M+Na]^+^ 532.32447; found, 532.32486.

### 2,3,4,6-Tetra-O-acetyl-β-D-13C6-glucopyranoside (20)

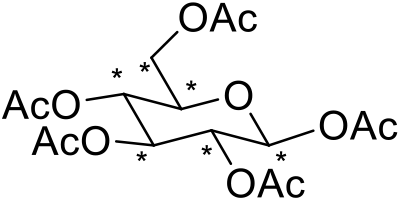

Sodium acetate (66 mg, 0.81 mmol) was dissolved in acetic anhydride (1.52 mL, 16.1 mmol) and brought to reflux. D-^13^C_6_-glucopyranoside (300 mg, 1.67 mmol) was added to the reaction mixture over a period of 10 minutes and stirred for 15 minutes. The reaction was allowed to reach room temperature and quenched by the addition of ice-cold H_2_O (15 mL) and was sonicated for 90 minutes. The precipitate was filtered and rinsed with H_2_O until the odor of acetic acid disappeared. Purification by column chromatography (20→80% Et_2_O in pentane) afforded compound **20** as white crystals (299 mg, 0.754 mmol, 45%). R*_f_* = 0.5 (3:1 Et_2_O/pentane). ^1^H NMR (500 MHz, CDCl_3_) δ 5.71 (d, *J* = 8.2 Hz, 1H, H-1), 5.25 (t, *J* = 9.5 Hz, 1H, H-3), 5.17 – 5.09 (m, 2H, H-2, H-4), 4.29 (dd, *J* = 12.5, 4.4 Hz, 1H, H-6a), 4.11 (d, *J* = 12.3 Hz, 1H, H-6b), 3.83 (d, *J* = 10.1 Hz, 1H, H-5), 2.12 (s, 3H, CH_3_), 2.09 (s, 3H, CH_3_), 2.04 (s, 3H, CH_3_), 2.02 (s, 3H, CH_3_). ^13^C NMR (126 MHz, CDCl_3_) δ 91.85 (dt, *J* = 48.4, 5.0 Hz, C-1), 73.53 – 72.36 (m, C-3, C-5), 70.36 (ddd, *J* = 48.5, 40.8, 3.4 Hz, C-2 or C-4), 67.88 (ddd, *J* = 42.1, 40.8, 3.5 Hz, C-2 or C-4), 61.59 (dt, *J* = 44.8, 4.3 Hz, C-6). HRMS calculated for C_10_^13^C_6_H_22_O_11_, [M+Na]^+^ 419.12556; found, 419.12568.

### 2’-Bromoethyl 2,3,4,6-tetra-*O*-acetyl-β-D-^13^C_6_-glucopyranoside (21)

To a cooled (0 °C) solution of compound **20** (299 mg, 0.754 mmol) in dry DCM (3 mL) 2-bromoethanol (53.2 μL, 0.754 mmol) was added. BF_3_.OEt_2_ (287 μL, 2.26 mmol) was added dropwise over 5 minutes and the reaction was stirred for 2 hours at 0 °C and overnight at room temperature. The mixture was diluted in EtOAc (20 mL) and washed with H_2_O (2×20 mL), NaHCO_3_ (2×20 mL), and brine (3×20 mL), dried (MgSO_4_), filtered and concentrated under reduced pressure. Purification of the residue by column chromatography (50→100 Et_2_O in pentane) afforded title compound (215 mg, 0.466 mmol, 62%) as a colorless oil. R*_f_* = 0.6 (3:1 Et_2_O/pentane).⍰^1^H NMR (500 MHz, CDCl_3_) δ 5.21 (dd, *J* = 10.4, 8.8 Hz, 1H, H-3), 5.12 – 5.05 (m, 1H, H-4), 5.01 (dd, *J* = 9.7, 7.9 Hz, 1H, H-2), 4.57 (d, *J* = 7.9 Hz, 1H, H-1), 4.25 (dd, *J* = 12.3, 4.8 Hz, 1H, H-6a), 4.20 – 4.11 (m, 2H, H-6b, H-7a), 3.86 – 3.78 (m, 1H, H-7b), 3.73 – 3.67 (m, 1H, H-5), 3.51 – 3.42 (m, 2H, H_2_-8), 2.09 (s, 3H, CH_3_), 2.07 (s, 3H, CH_3_), 2.02 (s, 3H, CH_3_), 2.01 (s, 3H, CH_3_). ^13^C NMR (126 MHz, CDCl_3_) δ 170.77, 170.38, 169.54 (4xC=O), 101.17 (dt, *J* = 48.2, 4.8 Hz, C-1), 73.45 – 70.59 (m, C-2, C-3, C-5), 68.45 (td, *J* = 41.4, 3.8 Hz, C-4), 61.98 (dt, *J* = 44.7, 4.3 Hz, C-6), 29.99 (C-8), 20.88, 20.74 (4xCH_3_). HRMS calculated for C_10_^13^C_6_H_23_BrO_10_, [M+NH_4_]^+^ 478.10141; found, 478.10124.

### 2’-Azidoethyl 2,3,4,6-tetra-*O*-acetyl-β-D-^13^C_6_-glucopyranoside (22)

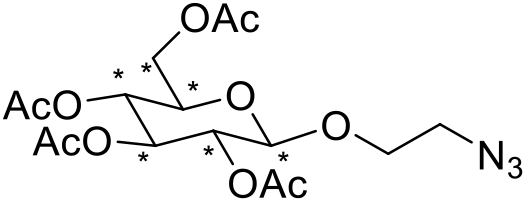

The title compound was synthesized from compound **21** following general procedure A on a 0.47 mmol scale. Purification by column chromatography (10→80% EtOAc in pentane) afforded compound **22** as white solid (173 mg, 0.409 mmol, 88%). R*_f_* = 0.8 (3:1 EtOAc/pentane). ^1^H NMR (500 MHz, CDCl_3_) δ 5.13 (t, *J* = 9.6 Hz, 1H, H-3), 5.01 (td, *J* = 9.6, 2.1 Hz, 1H, H-4), 4.93 (dd, *J* = 9.5, 7.9 Hz, 1H, H-2), 4.53 (d, *J* = 7.9 Hz, 1H, H-1), 4.18 (dd, *J* = 12.3, 4.7 Hz, 1H, H-6a), 4.08 (dd, *J* = 12.3, 2.3 Hz, 1H, H-6b), 3.99 – 3.92 (m, 1H, H-7a), 3.69 – 3.59 (m, 2H, H-5, H-7b), 3.46 – 3.38 (m, 1H, H-8a), 3.26 – 3.18 (m, 1H, H-8b), 2.01 (s, 3H, CH_3_), 1.97 (s, 3H, CH_3_), 1.95 (s, 3H, CH_3_), 1.92 (s, 3H, CH_3_). ^13^C NMR (126 MHz, CDCl_3_) δ 170.52, 170.13, 169.30 (4xC=O), 100.58 (dt, *J* = 48.3, 4.7 Hz, C-1), 73.23 – 70.54 (m, C-2, C-3, C-5), 68.25 (td, *J* = 41.5, 3.6 Hz, C-4), 61.78 (dt, *J* = 45.0, 4.2 Hz, C-6), 50.47 (C-8), 20.64, 20.59, 20.51 (4xCH_3_). HRMS calculated for C_10_^13^C_6_H_23_N_3_O_10_, [M+NH_4_]^+^ 441.19230; found, 441.19209.

### 2’-Palmitamidoethyl 2,3,4,6-tetra-*O*-acetyl-β-D-^13^C_6_-glucopyranoside (23)

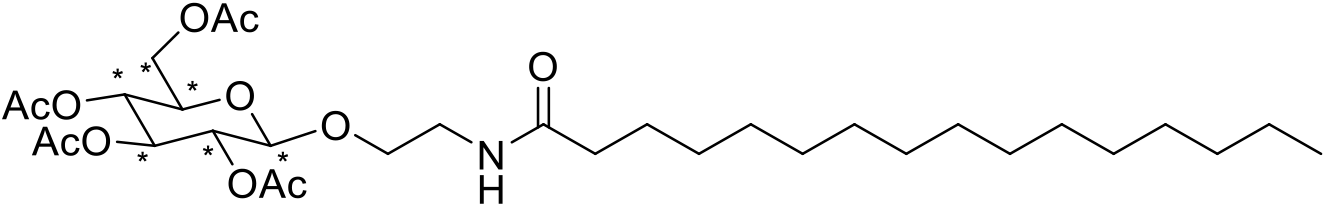

The title compound was synthesized from compound 22 following general procedure B on a 0.106 mmol scale and purified to afford compound **23** as a white solid (44.4 mg, 0.070 mmol, 66%). R*_f_* = 0.6 (3:1 EtOAc/pentane). ^1^H NMR (500 MHz, CDCl_3_) δ 5.88 (t, *J* = 5.8 Hz, 1H, NH), 5.18 (t, *J* = 9.6 Hz, 1H, H-3), 5.09 – 5.01 (m, 1H, H-4), 4.96 (dd, *J* = 9.6, 7.9 Hz, 1H, H-2), 4.48 (d, *J* = 7.9 Hz, 1H, H-1), 4.28 – 4.17 (m, 1H, H-6a), 4.12 (dd, *J* = 12.3, 2.2 Hz, 1H, H-6b), 3.85 – 3.77 (m, 1H, H-7a), 3.73 – 3.64 (m, 2H, H-5, H-7b), 3.50 – 3.37 (m, 2H, H_2_-8), 2.13 (t, *J* = 8.2 Hz, 2H, H_2_-9), 2.06 (s, 3H, CH_3_), 2.03 (s, 3H, CH_3_), 2.01 (s, 3H, CH_3_), 1.99 (s, 3H, CH_3_), 1.59 (p, *J* = 7.5 Hz, 2H, H_2_-10), 1.23 (bs, 24H, H_2_-11 to H_2_-22), 0.85 (t, *J* = 6.9 Hz, 3H, H_3_-23). ^13^C NMR (126 MHz, CDCl_3_) δ 173.35, 170.66, 170.24, 169.51 (5xC=O), 101.04 (ddd, *J* = 48.0, 5.6, 4.2 Hz, C-1), 73.47 – 70.85 (m, C-2, C-3, C-5), 68.34 (ddd, *J* = 42.1, 40.5, 3.9 Hz, C-4), 61.91 (dt, *J* = 44.7, 4.2 Hz, C-6), 39.19 (C-8), 36.79 (C-9), 32.01, 29.78, 29.77, 29.74, 29.72, 29.60, 29.47, 29.46, 29.44, 25.79, 22.77 (13xCH_2_, C-10 to C-22), 20.80, 20.67 (4xCH_3_), 14.21 (C-23). HRMS calculated for C_26_ ^13^C_6_H_55_NO_11_, [M+H]^+^ 636.40492; found, 636.40441.

### 2’-Oleamidoethyl 2,3,4,6-tetra-*O*-acetyl-β-D-^13^C_6_-glucopyranoside (24)

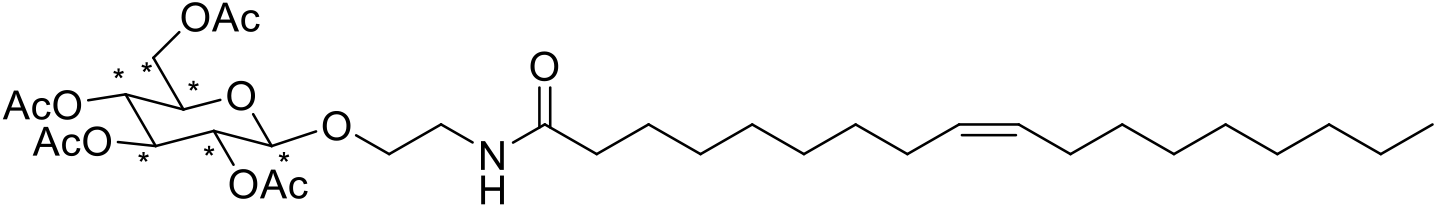

The title compound was synthesized from compound **22** following general procedure B on a 0.152 mmol scale and purified afford to compound **24** as a white solid (54.4 mg, 0.082 mmol, 54%). R*_f_* = 0.4 (3:1 EtOAc/pentane). ^1^H NMR (500 MHz, CDCl_3_) δ 5.89 (t, *J* = 5.8 Hz, 1H, NH), 5.36 – 5.27 (m, 2H, H-16, H-17), 5.18 (t, *J* = 9.6 Hz, 1H, H-3), 5.10 – 5.02 (m, 1H, H-4), 4.96 (dd, *J* =8) 6, 7.9 Hz, 1H, H-2), 4.48 (d, *J* = 7.9 Hz, 1H, H-1), 4.23 (dd, *J* = 12.4, 4.8 Hz, 1H, H-6a), 4.12 (dd, *J* = 12.2, 2.2 Hz, 1H, H-6b), 3.87 – 3.78 (m, 1H, H-7a), 3.73 – 3.64 (m, 2H, H-5, H-7b), 3.50 – 3.36 (m, 2H, H_2_-8), 2.13 (t, *J* = 8.0 Hz, 2H, H_2_-9), 2.06 (s, 3H, CH_3_), 2.03 (s, 3H, CH_3_), 2.01 – 1.96 (m, 10H, H_2_-15, H_2_-18, 2x CH_3_), 1.59 (p, *J* = 7.5 Hz, 2H, H_2_-10), 1.32 – 1.21 (m, 20H, H_2_-11 to H_2_-14, H_2_-19 to H_2_-24), 0.85 (t, *J* = 6.8 Hz, 3H, H_3_-25). ^13^C NMR (126 MHz, CDCl_3_) δ 173.32, 170.67, 170.26, 169.51 (5xC=O), 130.07, 129.83 (C-16, C-17), 101.04 (ddd, *J* = 48.0, 5.7, 4.2 Hz, C-1), 73.36 – 70.68 (m, C-2, C-3, C-5), 68.33 (ddd, *J* = 42.1, 40.4, 3.9 Hz, C-4), 61.90 (dt, *J* = 44.7, 4.2 Hz, C-6), 39.17 (C-8), 36.76 (C-9), 31.98, 29.85, 29.81, 29.60, 29.42, 29.40, 29.39, 29.38, 29.25, 27.30, 27.27, 25.76, 22.76 (13xCH_2_, C-10 to C-15, C-18 to C-24), 20.80, 20.67 4xCH_3_), 14.20 (C-25). HRMS calculated for C_28_^13^C_6_H_57_NO_11_, [M+H]^+^ 662.42057; found, 662.42072.

### 2’-(5Z,8Z,11Z,14Z)-Icosamidoethyl 2,3,4,6-tetra-*O*-acetyl-β-D-^13^C_6_-glucopyranoside (25)

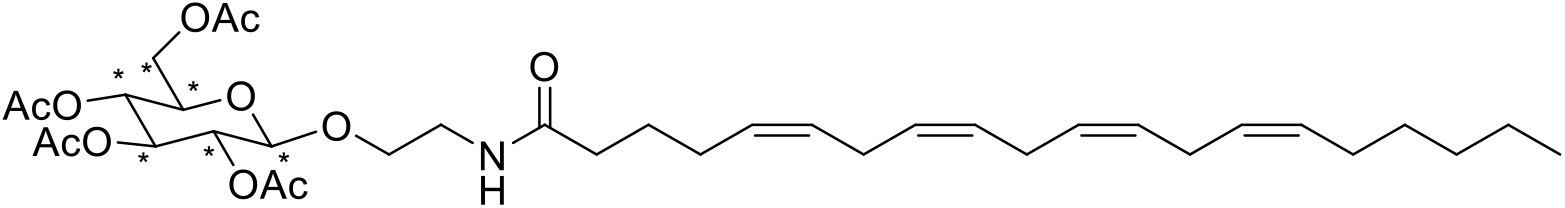

The title compound was synthesized from compound **22** following general procedure B on a 0.148 mmol scale and purified to afford compound **25** as a white solid (37.3 mg, 0.055 mmol, 37%). R*_f_* = 0.6 (3:1 EtOAc/pentane). ^1^H NMR (500 MHz, CDCl_3_) δ 5.87 (t, *J* = 5.7 Hz, 1H, NH), 5.43 – 5.28 (m, 8H, H-12, H-13, H-15, H-16, H-18, H-19, H-21, H-22), 5.19 (t, *J* = 9.6 Hz, 1H, H-3), 5.05 (t, *J* = 9.9 Hz, 1H, H-4), 4.96 (dd, *J* = 9.6, 7.9 Hz, 1H, H-2), 4.49 (d, *J* = 7.9 Hz, 1H, H-1), 4.24 (dd, *J* = 12.3, 4.8 Hz, 1H, H-6a), 4.13 (dd, *J* = 12.3, 2.2 Hz, 1H, H-6b), 3.87 – 3.78 (m, 1H, H-7a), 3.73 –3.64 (m, 2H, H-5, H-7b), 3.50 – 3.38 (m, 2H, H_2_-8), 2.84 – 2.75 (m, 6H, H_2_-14, H_2_-17, H_2_-20), 2.16 (t, *J* = 8.4 Hz, 2H, H_2_-9), 2.12 – 2.02 (m, 10H, H_2_-11, H_2_-23, 2x CH_3_), 2.01 (s, 3H, CH_3,_), 1.99 (s, 3H, CH_3_), 1.69 (p, *J* = 7.5 Hz, 2H, H_2_-10), 1.36 – 1.23 (m, 6H, H_2_-24, H_2_-25, H_2_-26), 0.87 (t, *J* = 6.9 Hz, 3H, H_3_-27). ^13^C NMR (126 MHz, CDCl_3_) δ 173.02, 170.69, 170.28, 169.53 (5xC=O), 130.61, 129.18, 128.89, 128.71, 128.35, 128.27, 127.96, 127.64 (C-12, C-13, C-15, C-16, C-18, C-19, C-21, C-22), 101.05 (ddd, *J* = 48.1, 5.7, 4.2 Hz, C-1), 73.36 – 70.71 (m, C-2, C-3, C-5), 68.34 (ddd, *J* = 42.1, 40.5, 3.9 Hz, C-4), 61.91 (dt, *J* = 44.6, 4.2 Hz, C-6), 39.23 (C-8), 36.06 (C-9), 31.61, 29.42, 27.32, 26.80, 25.58, 22.67 (9xCH_2_, C-10, C-11, C-14, C-17, C-20, C-23 to C- 26), 20.83, 20.69 (4xCH_3_), 14.18 (C-27). HRMS calculated for C_30_^13^C_6_H_55_NO_11_, [M+H]^+^ 684.40492; found, 684.40393.

### 2’-Palmitamidoethyl β-D-^13^C_6_-glucopyranoside (26)

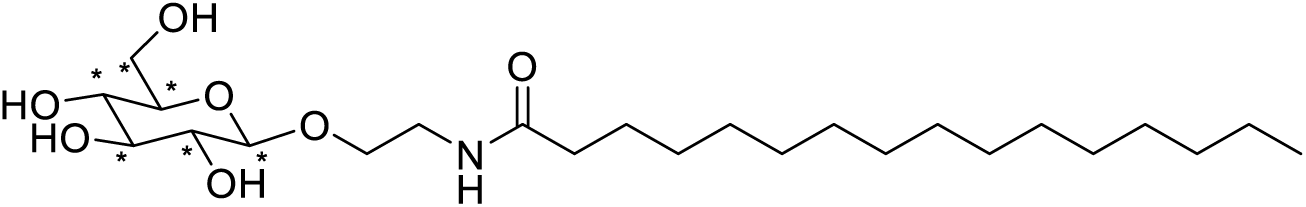

The title compound was synthesized from compound 22 on a 0.070 mmol scale following general procedure C and purified to afford compound **26** as a white solid (23.8 mg, 0.051 mmol, 73%). R*_f_* = 0.2 (1:9 MeOH/DCM). ^1^H NMR (500 MHz, methanol-d_4_) δ 4.27 (d, *J* = 7.8 Hz, 1H, H-1), 3.94 – 3.89 (m, 1H, H-7a), 3.87 (dd, *J* = 11.9, 1.4 Hz, 1H, H-6a), 3.69 - 3.62 (m, 2H, H-6b, H-7b), 3.50 – 3.44 (m, 1H, H-8a), 3.38 – 3.32 (m, 2H, H-3, H-8b), 3.29 – 3.26 (m, 2H, H-4, H-5), 3.19 (dd, *J* = 9.1, 7.7 Hz, 1H, H-2), 2.20 (t, *J* = 7.6 Hz, 2H, H_2_-9), 1.64 – 1.56 (m, 2H, H_2_-10), 1.29 (bs, 24H, H_2_-11 to H_2_-22), 0.90 (t, *J* = 7.1 Hz, 3H, H_3_-23). ^13^C NMR (126 MHz, methanol-d_4_) δ 176.49 (C=O), 104.53 (dt, *J* = 46.7, 4.0 Hz, C-1), 78.66 – 77.37 (m, C-3, C-4), 75.09 (ddd, *J* = 47.0, 39.3, 2.7 Hz, C-2), 71.60 (ddd, *J* = 41.4, 39.7, 2.6 Hz, C-5), 62.71 (dt, *J* = 43.5, 4.3 Hz, C-6), 40.60 (C-8), 37.12 (C-9), 33.08, 30.80, 30.79, 30.77, 30.76, 30.75, 30.65, 30.49, 30.48, 30.35, 27.03, 23.74 (13xCH_2_, C-10 to C-22), 14.44 (C-23). HRMS calculated for C_18_^13^C_6_H_47_NO_7_, [M+Na]^+^ 490.34460; found, 490.34424.

### 2’-Oleamidoethyl β-D-^13^C_6_-glucopyranoside (27)

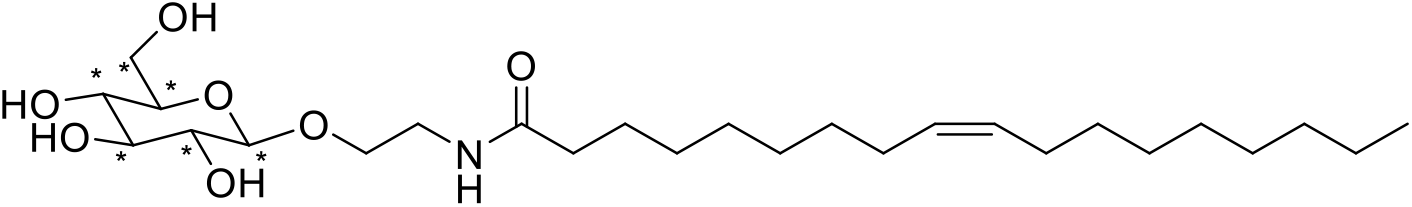

The title compound was synthesized from compound **24** on 0.082 mmol scale following general procedure C and purified to afford compound **27** as a white solid (33.8 mg, 0.068 mmol, 83%). R*_f_* = 0.3 (1:9 MeOH/DCM). ^1^H NMR (500 MHz, methanol-d_4_) δ 5.40 – 5.31 (m, 2H, H-16, H-17), 4.27 (d, *J* = 7.8 Hz, 1H, H-1), 3.94 – 3.89 (m, 1H, H-7a), 3.89 – 3.85 (m, 1H, H-6a), 3.68 – 3.62 (m, 2H, H-6b, H-7b), 3.50 – 3.44 (m, 1H, H-8a), 3.37 – 3.32 (m, 2H, H-3, H-8b), 3.29 – 3.26 (m, 2H, H-4, H-5), 3.19 (dd, *J* = 9.2, 7.7 Hz, 1H, H-2), 2.20 (t, *J* = 8.6 Hz, 2H, H_2_-9), 2.07 – 2.01 (m, 4H, H_2_-15, H_2_-18), 1.61 (q, *J* = 7.4 Hz, 2H, H_2_-10), 1.32 (d, *J* = 17.4 Hz, 20H, H_2_-11 to H_2_-14, H_2_-19 to H_2_-24), 0.90 (t, *J* = 7.3 Hz, 3H, H_3_-25). ^13^C NMR (126 MHz, methanol-d_4_) δ 176.45 (C=O), 130.88, 130.80 (C-16, C-17), 104.52 (dt, *J* = 46.7, 4.0 Hz, C-1), 79.79 – 76.83 (m, C-3, C-5), 75.09 (ddd, *J* = 47.0, 39.3, 2.7 Hz, C-2), 73.64 – 70.38 (m, C-4), 62.71 (dt, *J* = 43.7, 4.2 Hz, C-6), 40.57 (C-8), 37.11 (C-9), 33.06, 30.84, 30.61, 30.45, 30.38, 30.34, 30.25, 28.14, 28.12, 27.03, 23.74 (13xCH_2_, C-10 to C-15, C-18 to C-24), 14.45 (C-25). HRMS calculated for C_20_^13^C_6_H_49_NO_7_, [M+Na]^+^ 516.36025; found, 516.35993.

### 2’-(5Z,8Z,11Z,14Z)-Icosamidoethyl β-D-^13^C_6_-glucopyranoside (28)

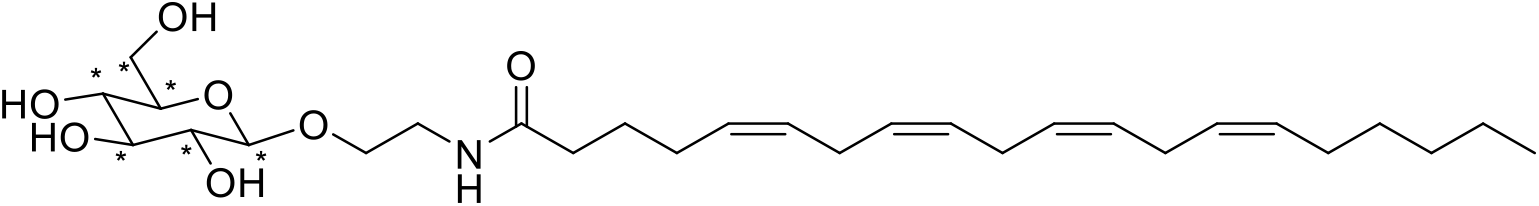

The title compound was synthesized from compound **25** on a 0.055 mmol scale following general procedure C and purified to afford compound **28** as a white solid (24.2 mg, 0.047 mmol, 86%). R*_f_* = 0.4 (1:9 MeOH/DCM). ^1^H NMR (500 MHz, methanol-d_4_) δ 5.42 – 5.31 (m, 8H, H-12, H-13, H-15, H-16, H-18, H-19, H-21, H-22), 4.27 (d, *J* = 7.7 Hz, 1H, H-1), 3.95 – 3.89 (m, 1H, H-7a), 3.87 (d, *J* = 11.7 Hz, 1H, H-6a), 3.68 – 3.61 (m, 2H, H-6b, H-7b), 3.51 – 3.43 (m, 1H, H- 8a), 3.39 – 3.32 (m, 2H, H-3, H-8b), 3.29 – 3.25 (m, 2H, H-4, H-5), 3.19 (dd, *J* = 9.2, 7.7 Hz, 1H, H-2), 2.87 – 2.80 (m, 6H, H_2_-14, H_2_-17, H_2_-20), 2.22 (t, *J* = 7.8 Hz,, 2H, H_2_-9), 2.16 – 2.04 (m, 4H, H_2_-11, H_2_-23), 1.68 (p, *J* = 7.5 Hz, 2H, H_2_-10), 1.41 – 1.26 (m, 6H, H_2_-24, H_2_-25, H_2_-26), 0.91 (t, *J* = 6.9 Hz, 3H, H_3_-27).^13^C NMR (126 MHz, methanol-d_4_) δ 176.15 (C=O), 131.21, 130.20, 129.74, 129.47, 129.22, 129.15, 128.91, 128.77 (C-12, C-13, C-15, C-16, C-18, C-19, C-21, C-22), 104.50 (dt, *J* = 46.8, 4.0 Hz, C-1), 80.10 – 75.84 (m, C-3, C-5), 75.08 (ddd, *J* = 47.1, 39.3, 2.6 Hz, C-2), 71.59 (ddd, *J* = 41.5, 39.7, 2.6 Hz, C-4), 62.71 (dt, *J* = 43.7, 4.2 Hz, C-6), 40.59 (C-8), 36.55 (C-9), 32.66, 30.47, 28.19, 27.74, 26.94, 26.57, 26.56, 26.55, 23.64 (9xCH_2_, C-10, C- 11, C-14, C-17, C-20, C-23 to C-26), 14.44 (C-27). HRMS calculated for C_22_^13^C_6_H_47_NO_7_, [M+Na]^+^ 538.34460; found, 538.34435.

### 2’-Bromoethyl 2,3,4,6-tetra-*O*-acetyl-α-D-^13^C_6_-glucopyranoside (29)

⍰D-^13^C_6_-glucopyranoside (0.200 g, 1.08 mmol) was suspended in 2-bromoethanol (1.14 mL, 16.1 mmol) and La(OTf)_3_ (21 mg, 0.035 mmol) was added and the reaction mixture stirred for 16 hours at 60°C. The reaction mixture was cooled to room temperature and purified by column chromatography (5→20% MeOH in DCM) to 2’-bromoethyl α/β-D-^13^C_6_-glucopyranoside as an inseparable mixture (230 mg, 73%). The crude mixture (230 mg, 0.783 mmol) was dissolved in pyridine (3 mL) and was cooled to 0 °C. Acetic anhydride (1.48 mL, 15.7 mmol) and cat. DMAP were added and the reaction was stirred overnight at room temperature. The reaction mixture was diluted with Et_2_O (20 mL) and washed with H_2_O (2×20 mL), 10% aq. HCl (2×20 mL), NaHCO_3_ (2×20 mL), H_2_O (2×20 mL) and brine (3×20 mL). The organic layer was dried (MgSO_4_), filtered and concentrated. Purification of the residue by silica gel column chromatography (5→20% EtOAc in CHCl_3_) afforded compound **29** as a white solid (76.1 mg, 0.165 mmol, 22% over two steps). R*_f_* = 0.5 (1:9 EtOAc/CHCl_3_). ^1^H NMR (500 MHz, CDCl_3_) δ 5.46 (t, *J* = 9.8 Hz, 1H, H-3), 5.12 (d, *J* = 3.9 Hz, 1H, H-1), 5.03 (t, *J* = 9.7 Hz, 1H, H-4), 4.83 (dd, *J* = 10.0, 3.8 Hz, 1H, H-2), 4.22 (dd, *J* = 12.1, 4.4 Hz, 1H, H-6a), 4.16 – 4.05 (m, 2H, H-5, H-6b), 4.01 – 3.93 (m, 1H, H-7a), 3.87 – 3.77 (m, 1H, H-7b), 3.49 (t, *J* = 5.8 Hz, 2H, H_2_-8), 2.06 (d, *J* =6) 9 Hz, 6H, 2xCH_3_), 2.00 (t, *J* = 8.6 Hz, 6H, 2x CH_3_). ^13^C NMR (126 MHz, CDCl_3_) δ 170.71, 170.34, 170.15, 169.71 (4xC=O), 96.04 (dt, *J* = 46.2, 2.5 Hz, C-1), 71.53 – 69.54 (m, C-2, C-3), 69.21 – 66.94 (m, C-4, C-5), 61.89 (dt, *J* = 45.0, 3.5 Hz, C-6), 30.04 (C-8), 20.82 (2xCH_3_), 20.78 (CH_3_), 20.71 (CH_3_). HRMS calculated for C_10_^13^C_6_H_23_BrO_10_, [M+Na]^+^ 483.05681; found, 483.05677.

### 2’-Azidoethyl 2,3,4,6-tetra-*O*-acetyl-α-D-^13^C_6_-glucopyranoside (30)

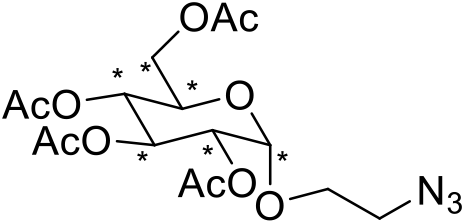

The title compound was synthesized from compound **29** following general procedure A on a 0.165 mmol scale. Purification by column chromatography (10→80% EtOAc in pentane) afforded compound **30** as white solid (68.2 mg, 0.161 mmol, 98%). R*_f_* = 0.8 (3:1 EtOAc/pentane). ^1^H NMR (500 MHz, CDCl_3_) δ 5.46 (t, *J* = 9.6 Hz, 1H, H-3), 5.09 (d, *J* = 4.5 Hz, 1H, H-1), 5.03 (t, *J* = 10.0 Hz, 1H, H-4), 4.85 (dd, *J* = 10.6, 4.1 Hz, 1H, H-2), 4.22 (dd, *J* = 12.8, 5.0 Hz, 1H, H-6a), 4.07 (dd, *J* = 12.5, 2.6 Hz, 1H, H-6b), 4.05 – 3.98 (m, 1H, H-5), 3.88 – 3.80 (m, 1H, H-7a), 3.64 – 3.57 (m, 1H, H-7b), 3.50 – 3.43 (m, 1H, H-8a), 3.41 – 3.35 (m, 1H, H- 8b), 2.06 (s, 3H, CH_3_), 2.04 (s, 3H, CH_3_), 2.00 (s, 3H, CH_3_), 1.98 (s, 3H, CH_3_). ^13^C NMR (126 MHz, CDCl_3_) δ 170.65, 170.35, 170.07, 169.66 (4xC=O), 96.10 (ddt, *J* = 44.9, 3.7, 1.9 Hz, C-1), 71.44 – 69.46 (m, C-2, C-3), 69.29 – 67.02 (m, C-4, C-5), 62.45 – 60.66 (m, C-6), 50.45 (C-8), 20.77, 20.72, 20.71, 20.66 (4xCH_3_). HRMS calculated for C_10_^13^C_6_H_23_N_3_O_10_, [M+Na]^+^ 446.14769; found, 446.14743.

### 2’-Palmitamidoethyl 2,3,4,6-tetra-*O*-acetyl-α-D-^13^C_6_-glucopyranoside (31)

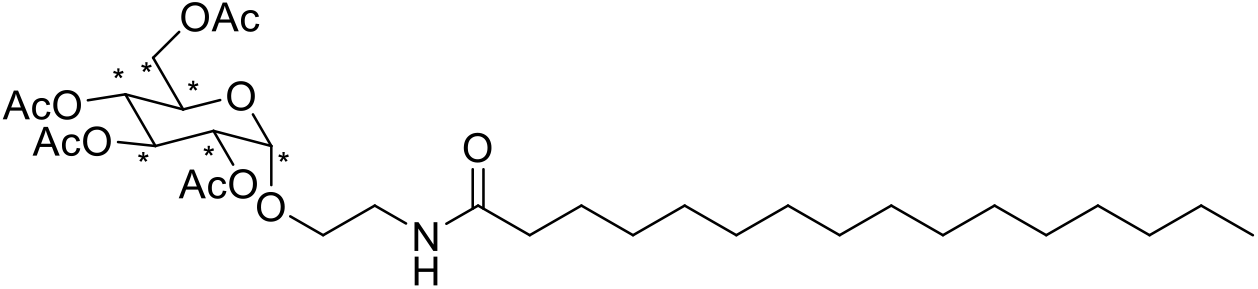

The title compound was synthesized from compound **30** following general procedure B on a 0.050 mmol scale and purified to afford compound **31** as a white solid (21.5 mg, 0.034 mmol, 67%). R*_f_* = 0.6 (3:1 EtOAc/pentane). ^1^H NMR (500 MHz, CDCl_3_) δ 5.91 (t, *J* = 5.8 Hz, 1H, NH), 5.48 (t, *J* = 9.7 Hz, 1H, H-3), 5.06 – 4.98 (m, 2H, H-1, H-4), 4.89 (dd, *J* = 10.2, 3.5 Hz, 1H, H-2), 4.18 (d, *J* = 10.9 Hz, 1H, H-6a), 4.09 (d, *J* = 12.4 Hz, 1H, H-6b), 4.04 – 3.98 (m, 1H, H-5), 3.79 – 3.72 (m, 1H, H-7a), 3.57 – 3.45 (m, 3H, H-7b, H_2_-8), 2.18 (t, *J* = 7.7 Hz, 2H, H_2_-9), 2.08 (s, 3H, CH_3_), 2.06 (s, 3H, CH_3_), 2.03 (s, 3H, CH_3_), 2.02 (s, 3H, CH_3_), 1.65 – 1.59 (m, 2H, H_2_-10), 1.24 (s, 24H, H_2_-11 to H_2_-22), 0.87 (t, *J* = 6.9 Hz, 3H, H_3_-23). ^13^C NMR (126 MHz, CDCl_3_) δ 173.34, 170.78, 170.37, 170.03, 169.68 (5xC=O), 96.09 (dt, *J* = 45.9, 2.6 Hz, C-1), 71.39 – 69.67 (m, C-2, C-3), 69.39 – 67.00 (m, C-4, C-5), 62.02 (dt, *J* = 43.8, 3.5 Hz, C-6), 38.83 (C-8), 36.89 (C-9), 32.04, 30.47, 29.82, 29.82, 29.78, 29.76, 29.63, 29.50, 29.48, 29.04, 25.86, 23.85, 23.11 (13xCH_2_, C-10 to C-22), 20.84, 20.79, 20.75 (4xCH_3_), 14.25 (C-23). HRMS calculated for C_26_^13^C_6_H_55_NO_11_, [M+H]^+^ 636.40492; found, 636.40471.

### 2’-Oleamidoethyl 2,3,4,6-tetra-*O*-acetyl-α-D-^13^C_6_-glucopyranoside (32)

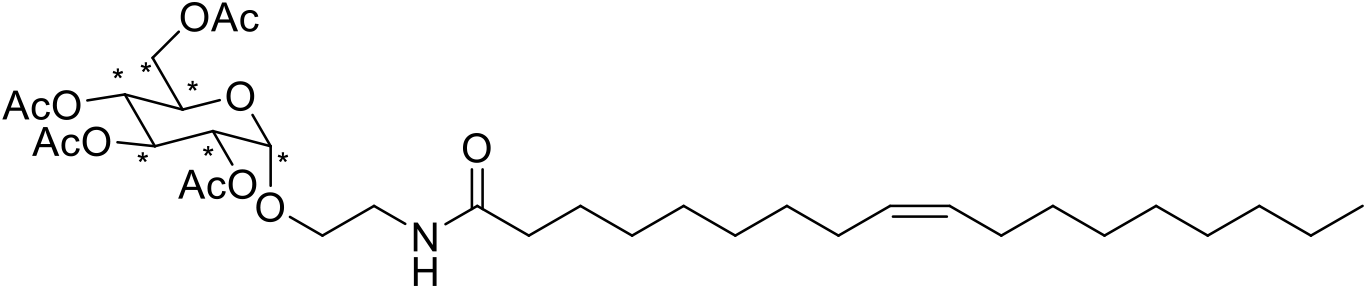

The title compound was synthesized from compound **30** following general procedure B on a 0.050 mmol scale and purified to afford compound **32** as a white solid (18.2 mg, 0.028 mmol, 55%). R*_f_* = 0.6 (3:1 EtOAc/pentane). ^1^H NMR (500 MHz, CDCl_3_) δ 5.92 (t, *J* = 5.8 Hz, 1H, NH), 5.47 (t, *J* = 9.8 Hz, 1H, H-3), 5.37 – 5.29 (m, 2H, H-16, H-17), 5.06 – 4.98 (m, 2H, H-1, H-4), 4.88 (dd, *J* = 10.2, 3.5 Hz, 1H, H-2), 4.22 (dd, *J* = 12.3, 4.7 Hz, 1H, H- 6a), 4.09 (dd, *J* = 12.3, 2.2 Hz, 1H, H-6b), 4.05 – 3.97 (m, 1H, H-5), 3.79 – 3.71 (m, 1H, H-7a), 3.57 – 3.51 (m, 1H, H-7b), 3.51 – 3.44 (m, 2H, H_2_-8), 2.18 (t, *J* = 8.0 Hz, 2H, H_2_-9), 2.08 (s, 3H, CH_3_), 2.06 (s, 3H, CH_3_), 2.03 (s, 3H, CH_3_), 2.02 – 1.97 (m, 7H, H_2_-15, H_2_-18, CH_3_), 1.61 (q, *J* = 7.3 Hz, 2H, H_2_-10), 1.34 – 1.22 (m, 20H, H_2_-11 to H_2_-14, H_2_-19 to H_2_-24), 0.87 (t, *J* = 6.8 Hz, 3H, H_3_-25). ^13^C NMR (126 MHz, CDCl_3_) δ 173.33, 170.75, 170.37, 170.02, 169.67 (5xC=O), 96.09 (dt, *J* = 45.8, 2.6 Hz, C-1), 71.43 – 69.63 (m, C-2, C-3), 68.75 (td, *J* = 39.5, 38.4, 5.8 Hz, C-4), 67.54 (ddt, *J* = 43.3, 40.7, 2.4 Hz, C-5), 62.01 (dt, *J* = 43.9, 3.5 Hz, C-6), 39.11 (C-8), 36.84 (C-9), 32.02, 29.88, 29.85, 29.64, 29.46, 29.44, 29.43, 29.41, 29.28, 27.34, 27.30, 22.80 (13xCH_2_, C-10 to C-15, C-18 to C-24), 20.83, 20.79, 20.75 (4xCH_3_), 14.24 (C-25). HRMS calculated for C_28_ ^13^C_6_H_57_NO_11_, [M+H]^+^ 662.42057; found, 662.42013.

### 2’-(5Z,8Z,11Z,14Z)-Icosamidoethyl 2,3,4,6-tetra-*O*-acetyl-α-D-^13^C_6_-glucopyranoside (33)

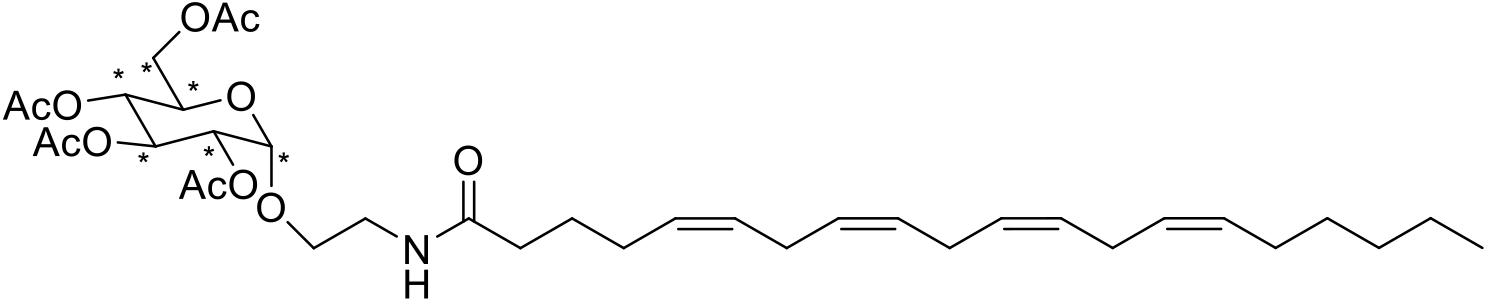

The title compound was synthesized from compound **30** following general procedure B on a 0.060 mmol scale and purified to afford compound **33** as a white solid (8.4 mg, 0.012 mmol, 20%). R*_f_* = 0.6 (3:1 EtOAc/pentane). ^1^H NMR (500 MHz, CDCl_3_) δ 5.88 (t, *J* = 5.8 Hz, 1H, NH), 5.48 (t, *J* = 9.8 Hz, 1H, H-3), 5.44 – 5.29 (m, 8H, H-12, H-13, H-15, H-16, H-18, H-19, H-21, H-22), 5.07 – 4.99 (m, 2H, H-1, H-4), 4.89 (dd, *J* = 10.2, 3.6 Hz, 1H, H-2), 4.23 (dd, *J* = 12.2, 4.8 Hz, 1H, H-6a), 4.10 (dd, *J* = 12.2, 2.1 Hz, 1H, H-6b), 4.04 –3.98 (m, 1H, H-5), 3.79 – 3.73 (m, 1H, H-7a), 3.57 – 3.44 (m, 3H, H-7b, H_2_-8), 2.90 – 2.74 (m, 6H, H_2_-14, H_2_-17, H_2_-20), 2.24 – 2.17 (m, 2H, H_2_-9), 2.15 – 1.98 (m, 18H, H_2_-11, H_2_-23, 4xCH_3_), 1.72 (p, *J* = 7.5 Hz, 2H, H_2_-8), 1.37 – 1.23 (m, 6H, H_2_-24, H_2_-25, H_2_-26), 0.88 (t, *J* = 6.9 Hz, 3H, H_3_-27). ^13^C NMR (126 MHz, CDCl_3_) δ 172.98, 170.78, 170.39, 170.07, 169.71 (5xC=O), 130.67, 129.14, 129.02, 128.77, 128.43, 128.25, 127.97, 127.65 (C-12, C-13, C-15, C-16, C-18, C-19, C-21, C-22), 96.11 (dt, *J* = 45.7, 2.7 Hz, H-1), 71.34 – 69.66 (m, C-2, C-3), 68.78 (td, *J* = 39.5, 38.2, 5.9 Hz, C-4), 67.56 (ddt, *J* = 43.4, 40.8, 2.4 Hz, C-5), 62.03 (dt, *J* = 44.0, 3.5 Hz, C-6), 39.11 (C-8), 36.15 (C-9), 31.66, 29.46, 27.36, 26.83, 25.78, 25.77, 25.64, 22.72 (9xCH_2_, C-10, C-11, C-14, C-17, C-20, C-23 to C-26), 20.87, 20.85, 20.82, 20.78 (4xCH_3_), 14.23 (C-27). HRMS calculated for C_30_^13^C_6_H_55_NO_11_, [M+Na]^+^ 706.38686; found, 706.38633.

### 2’-Palmitamidoethyl α-D-^13^C_6_-glucopyranoside (34)

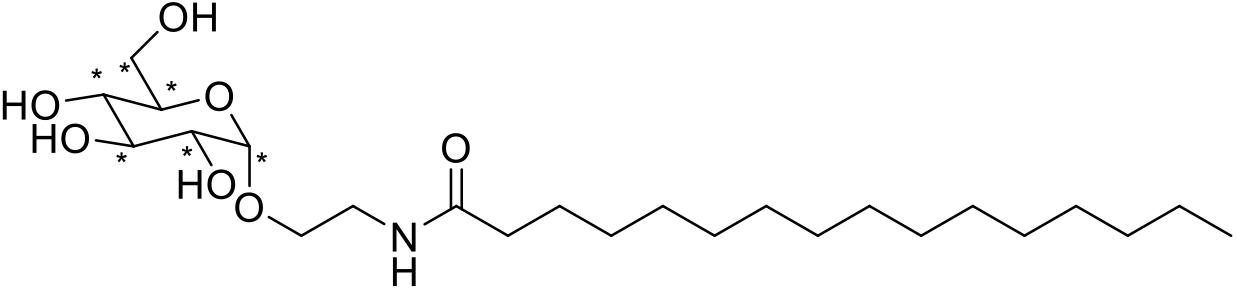

The title compound was synthesized from compound **31** on a 0.034 mmol scale following general procedure C and purified to afford compound **34** as a white solid (11.1 mg, 0.024 mmol, 70%). R*_f_* = 0.2 (1:9 MeOH/DCM). ^1^H NMR (500 MHz, methanol-d_4_) δ 4.79 (d, *J* = 3.7 Hz, 1H, H-1), 3.82 – 3.77 (m, 2H, H-6a, H-7a), 3.68 – 3.61 (m, 2H, H-3, H-6b), 3.58 – 3.52 (m, 2H, H-4, H-8a), 3.51 – 3.46 (m, 1H, H-7b), 3.40 (dd, *J* = 9.7, 3.6 Hz, 1H, H-2), 3.30 –3.25 (m, 2H, H-5, H-8b), 2.20 (t, *J* = 8.2 Hz, 2H, H_2_-9), 1.60 (p, *J* = 7.3 Hz, 2H, H_2_-10), 1.29 (bs, 24H, H_2_-11 to H_2_-22), 0.90 (t, *J* = 6.9 Hz, 3H, H_3_-23). ^13^C NMR (126 MHz, methanol-d_4_) δ 176.46 (C=O), 100.27 (ddd, *J* = 46.1, 3.4, 1.9 Hz, C-1), 75.11 (tdd, *J* = 38.0, 3.8, 1.8 Hz, C-3), 74.35 – 73.00 (m, C-2, C-4), 71.71 (ddd, *J* = 40.8, 38.0, 3.2 Hz, C-5), 62.65 (dt, *J* = 43.3, 3.6 Hz, C-6), 40.22 (C-8), 37.15 (C-9), 33.08, 30.81, 30.80, 30.78, 30.77, 30.76, 30.65, 30.49, 27.05, 23.74 (13xCH_2_, C-10 to C-22), 14.45 (C-23). HRMS calculated for C_18_ ^13^C_6_H_47_NO_7_, [M+H]^+^ 468.36266; found, 468.36260.

### 2’-Oleamidoethyl α-D-^13^C_6_-glucopyranoside (35)

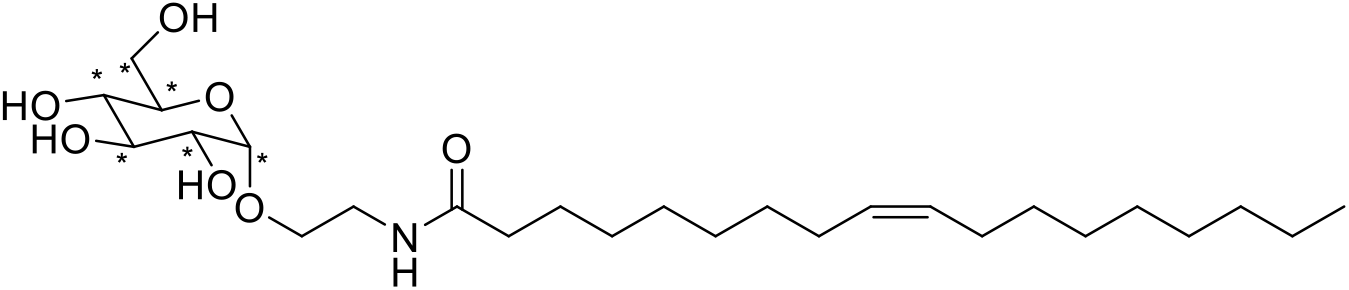

The title compound was synthesized from compound **32** on a 0.028 mmol scale following general procedure C and purified to afford compound **35** as a white solid (13 mg, 0.026 mmol, 96%). R*_f_* = 0.3 (1:9 MeOH/DCM). ^1^H NMR (500 MHz, methanol-d_4_) δ 5.38 – 5.31 (m, 2H, H-16, H-17), 4.79 (d, *J* = 3.7 Hz, 1H, H-1), 3.83 – 3.77 (m, 2H, H-6a, H-7a), 3.68 – 3.61 (m, 2H, H-3, H-6b), 3.58 – 3.52 (m, 2H, H-4, H-8a), 3.51 – 3.46 (m, 1H, H-7b), 3.41 (dd, *J* = 9.7, 3.7 Hz, 1H, H-2), 3.30 – 3.25 (m, 2H, H-5, H-8b), 2.20 (t, *J* = 7.8 Hz, 2H, H_2_-9), 2.07 – 2.00 (m, 4H, H_2_-15, H_2_-18), 1.62 (p, *J* = 7.3 Hz, 2H, H_2_-10), 1.32 (d, *J* = 18.2 Hz, 20H, H_2_-11 to H_2_-14, H_2_-19 to H_2_-24), 0.90 (t, *J* = 6.9 Hz, 3H, H_3_-25). ^13^C NMR (126 MHz, methanol-d_4_) δ 176.42 (C=O), 130.87, 130.80 (C-16, C-17), 100.26 (ddd, *J* = 46.3, 3.5, 1.9 Hz, C-1), 75.10 (tdd, *J* = 38.0, 3.8, 1.9 Hz, C-3), 74.39 – 72.91 (m, C-2, C-4), 71.70 (ddd, *J* = 40.8, 38.0, 3.2 Hz, C-5), 62.65 (dt, *J* = 43.3, 3.6 Hz, C-6), 40.17 (C-8), 37.14 (C-9), 33.07, 31.62, 30.85, 30.61, 30.45, 30.37, 30.34, 30.25, 30.14, 28.15, 28.12, 27.04, 23.74 (13xCH_2_, C-10 to C-15, C-18 to C-24), 14.46 (C-25). HRMS calculated for C_20_^13^C_6_H_49_NO_7_, [M+H]^+^ 494.37831; found, 494.37827.

### 2’-(5Z,8Z,11Z,14Z)-Icosamidoethyl α-D-^13^C_6_-glucopyranoside (36)

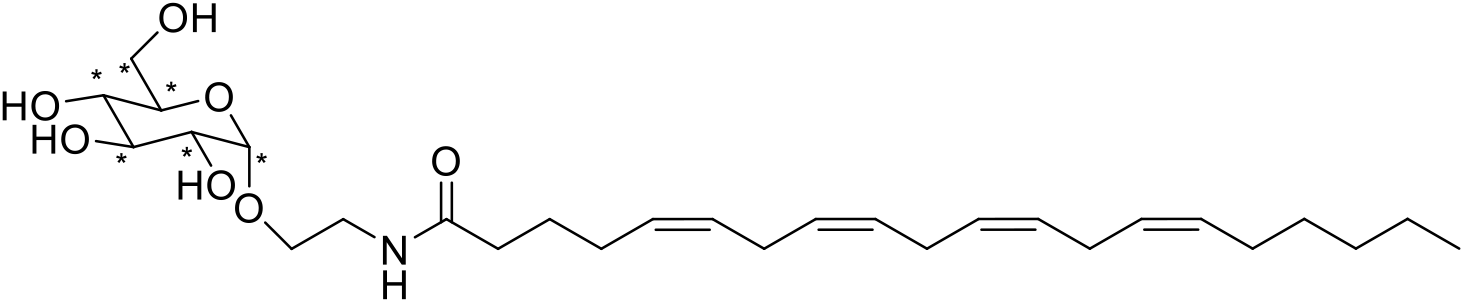

The title compound was synthesized from compound **33** on a 0.012 mmol scale following general procedure C and purified to afford compound **36** as a white solid (5 mg, 0.010 mmol, 82%). R*_f_* = 0.4 (1:9 MeOH/DCM). ^1^H NMR (500 MHz, methanol-d_4_) δ 5.43 – 5.31 (m, 8H, H-12, H-13, H-15, H-16, H-18, H-19, H-21, H-22), 4.78 (d, *J* = 3.7 Hz, 1H, H-1), 3.83 – 3.76 (m, 2H, H-6a, H-7a), 3.68 – 3.60 (m, 2H, H-3, H-7b), 3.59 – 3.51 (m, 2H, H-4, H-8a), 3.52 – 3.45 (m, 1H, H-7b), 3.40 (dd, *J* = 9.6, 3.7 Hz, 1H, H-2), 3.29 – 3.25 (m, 2H, H-5, H-8b), 2.88 – 2.79 (m, 6H, H_2_-14, H_2_-17, H_2_-20), 2.22 (t, *J* = 8.0 Hz, 2H, H_2_-9), 2.16 – 2.04 (m, 4H, H_2_- 11, H_2_-23), 1.72 – 1.65 (m, 2H, H_2_-10), 1.37 – 1.27 (m, 6H, H_2_-24 to H_2_-26), 0.91 (t, *J* = 6.3 Hz, 3H, H_3_-27). ^13^C NMR (214 MHz, methanol-d_4_) δ 176.13 (C=O), 131.20, 130.18, 129.74, 129.46, 129.21, 129.14, 128.90, 128.77 (C-12, C-13, C-15, C-16, C-18, C-19, C-21, C-22), 100.23 (ddd, *J* = 46.5, 3.5, 2.0 Hz, C-1), 75.08 (tdd, *J* = 38.4, 3.8, 1.7 Hz, C-3), 74.26 – 73.03 (m, C-2, C-4), 71.70 (ddd, *J* = 40.9, 38.4, 2.9 Hz, C-5), 62.64 (dt, *J* = 43.3, 3.6 Hz, C-6), 40.21 (C-8), 36.57 (C-9), 32.68, 30.49, 28.19, 27.77, 26.96, 26.56, 26.56, 26.55, 23.66 (9xCH_2_, C-10, C-11, C-14, C-17, C-20, C-23 to C-26), 14.46 (C-27). HRMS calculated for C_22_ ^13^C_6_H_47_NO_7_, [M+Na]^+^ 538.34460; found, 538.34443.

### Preparation of standards and calibration curves

β-Glyco-NAE standards and isotope-labelled internal standards were dissolved at 10 mM in ethanol and further diluted to 1 μM in methanol. Standard working solutions were prepared at 667 pM and internal standard working solutions at 3,330 pM for each analyte and stored at −80 °C. Calibration curves were prepared by serial dilution of the standard working solution in methanol and mixed 1:1 with the internal standard mixture before analysis. Final calibration concentrations were 0, 14, 28, 69, 139, 174, 278 and 694 pM. Internal standard concentrations were selected to approximate endogenous analyte concentrations.

### LC–MS/MS analysis of β-Glyco-NAEs

Targeted LC–MS/MS analysis of β-Glc-AEA, β-Glc-OEA, β-Glc-PEA, β-Gal-AEA, β-Gal-OEA, β-Gal-PEA and their isotope-labelled internal standards was performed on an Acquity UPLC I-Class system coupled to a Xevo TQ-S micro triple quadrupole mass spectrometer (Waters). Chromatographic separation was achieved using an Acquity UPLC HSS T3 column (2.1 × 100 mm, 1.8 μm) equipped with an Acquity UPLC HSS T3 VanGuard pre-column (2.1 × 5 mm, 1.8 μm), maintained at 45 °C.

Mobile phase A consisted of 2 mM ammonium formate and 10 mM formic acid in water. Mobile phase B was acetonitrile. The flow rate was 0.55 ml min^−1^. The gradient started at 25% B for 0.5 min, increased to 52.5% B over 1.5 min, then to 58% B over 5 min, followed by an increase to 100% B over 1.5 min. The column was held at 100% B for 0.5 min and then returned to starting conditions for 2.5 min before the next injection. Total run time was 11.5 min. The autosampler was maintained at 5 °C and 10 μL of each sample was injected.

Analytes were detected in electrospray-positive mode using selected multiple reaction monitoring. Source settings were as follows: capillary voltage, 3.20 kV; declustering potential, 40 V; source temperature, 150 °C; desolvation temperature, 500 °C; cone gas, 50 l h^−1^; desolvation gas, 500 l h^−1^; dwell time, 0.061 s; interchannel delay, 0.5 s; and interscan delay, 0.5 s. MRM transitions were optimized by direct infusion of synthetic standards. The principal transitions were m/z 510.3 > 348.3 for β-Glc/Gal-AEA, m/z 488.4 > 326.3 for β-Glc/Gal-OEA and m/z 462.3 > 300.3 for β-Glc/Gal-PEA. Corresponding ^13^C6-labelled standards were monitored using m/z 516.3 > 348.3, m/z 494.3 > 326.3 and m/z 468.4 > 300.3, respectively.

### LC–MS/MS method validation

The β-Glyco-NAE LC–MS/MS method was validated using mouse brain homogenate and human plasma. Selectivity and specificity were assessed by analyzing blank matrix, non-spiked matrix and matrix spiked with β-Glyco-NAE standards. Carry-over was assessed by injection of blank acetonitrile/water samples between analyte-containing samples. Linearity was evaluated using calibration curves generated in HEPES buffer. Limits of detection and quantification were calculated as 3 × S_a_/b and 10 × S_a_/b, respectively, where S_a_ is the standard deviation of the y-intercept and b is the slope of the calibration curve.

Intraday and interday precision were determined over four days using low, medium and high standard concentrations spiked into mouse brain homogenate or human plasma. Matrix effects were determined by comparing the response of isotope-labelled internal standards spiked after extraction into matrix with that of standards spiked into blank buffer. Recovery was determined by comparing the response of internal standards spiked before extraction with that of internal standards spiked after extraction.

### Lipid extraction for β-Glyco-NAE analysis

Lipid extraction was performed on ice. Samples were thawed and spiked with 10 μL of ^13^C_6_-labelled internal standard mixture, vortexed and incubated for 5 min on ice. Subsequently, 100 μL of 0.5% sodium chloride and 100 μL of 0.1 M ammonium acetate buffer, pH 4, were added. Lipids were extracted with 1 ml MTBE by mixing for 7 min, followed by centrifugation at 16,000 *g* for 11 min at 4 °C. The upper organic layer was transferred to a clean tube and dried under vacuum. Dried lipid extracts were reconstituted in 30 μL acetonitrile/water (50:50, v/v), mixed and centrifuged at 10,000 *g* for 4 min at 4 °C. Supernatants were transferred to LC–MS vials for analysis.

### Analysis of NAEs, glycosphingolipids and glycosylated cholesterol

Parent NAEs were measured by targeted LC–MS/MS using established methods^69^. Glucosylceramide, galactosylceramide, glucosylcholesterol and galactosylcholesterol were quantified by targeted lipidomics as described previously^83^. For all lipidomics measurements, analyte peak areas were normalized to isotope-labelled internal standards where available and corrected for protein concentration or sample volume as appropriate.

### Cell culture

HEK293T (human embryonic kidney, CRL-3216), Neuro-2a (murine neuroblastoma, CCL-131) and RAW264.7 (mouse macrophages, TIB-71) cells were cultured under standard humidified conditions at 37 °C and 7% CO₂ for HEK293T and Neuro-2a and 5% CO2 for RAW264.7. N9 mouse microglial cells were cultured in T75 flasks (Sarstedt) in scove’s Modified Dulbecco’s Medium (IMDM), supplemented with 5% (v/v) sterile-filtered (0.2 μM) FCS, glutamax and pen/strep at 37 °C, 5% CO_2_. HEK293T and Neuro-2a cells were maintained in DMEM (high glucose, Sigma-Aldrich, #D6546) supplemented with glutamax (2 mM final concentration, Sigma-Aldrich), penicillin (pen)/streptomycin (strep) (both 200 μg/mL, Duchefa) and 10% new-born calf serum (NCS, Seradigm). Neuro-2a cells were differentiated with retinoic acid before lipid extraction where indicated. Medium was refreshed every 2 or 3 days and cells were passaged twice a week at 80-90 % confluence, by resuspension via pipetting in fresh medium. RAW264.7 macrophages were maintained in DMEM high glucose supplemented with 10% fetal calf serum (FCS, Seradigm), pen/strep and glutamax. Medium was refreshed every 2-3 days and cells were passaged two times a week at 80-90 % confluence by detaching cells with 0.25% Trypsin in PBS/EDTA and resuspension in fresh medium. All cell lines were purchased at ATCC and checked regularly for mycoplasma contamination. Cell viability was assessed by Trypan Blue exclusion and quantification using a TC20™ Automated Cell Counter (Bio-Rad). Cultures were discarded after 2-3 months of use..

### Generation of GBA2-overexpressing cell lysates

HEK293T cells were transiently transfected with a plasmid encoding human GBA2 as described previously^84^.

### *In vitro* GBA2 transglycosylation assays

For *in vitro* transglycosylation assays, lysates from GBA2-overexpressing HEK293T cells were incubated with AEA, OEA or PEA and either Glc-4MU or Gal-4MU as glycosyl donor. Cells were lysed on ice in lysis buffer (containing 25 mM potassium phosphate buffer pH 6.5, supplemented with 0.1% (v/v) Triton X-100) using a Vibra-Cell™ VCX130 (Sonics; Newtown, USA) for 1 s ON/19 s OFF (5 cycles) at 20% amplitude. Protein concentration was determined with the Pierce™ bicinchoninic acid assay (BCA) using Protein Assay Kit (Thermo Fisher Scientific) and samples were diluted to 2 mg/mL using lysis buffer. Diluted lysates (16 µL) were pre-incubated with vehicle (DMSO), GBA2 inhibitor (MZ31, 25 nM^77^) or GCase/GBA2 inhibitor (JJB367, 200 nM ^85^) for 1 h at 37 °C. Final DMSO (v/v) concentration in all samples was 0.1%. After which samples were incubated with or without, AEA-NBD (2 nmol) or Glc-4MU (3 mM) was added in the appropriate enzyme buffer (84 µL) for GCase as described above or for GBA2 (consisting of 150 mM McIlvaine buffer pH 5.8, supplemented with 0.1% (w/v) BSA). Where indicated, lysates were preincubated with GCase or GBA2 inhibitors before addition of substrates. Reactions were stopped by lipid extraction with MTBE, and β-Glc- or β-Gal-NAE formation was quantified by LC–MS/MS. For kinetic experiments, NAE acceptor concentrations were varied over the indicated concentration range while donor concentration and protein concentration were kept constant. Initial rates were calculated from product formation and fitted to the Michaelis–Menten equation using nonlinear regression using Graphpad Prism v9.

### Recombinant GCase hydrolysis assays

Human recombinant GCase (Imiglucerase, Cerezyme®, obtained from Sanofi Genzyme) was incubated with β-Glc-AEA, β-Glc-OEA, β-Glc-PEA or their β-Gal counterparts in acidic assay buffer (150 mM McIlvaine buffer pH 5.2, supplemented with 0.1% (w/v) BSA, 0.1% (w/v) Triton X-100 and 0.2% (w/v) NaTc, all purchased from Sigma). Reactions were performed at 37 °C for 1 hour and stopped by lipid extraction. Formation of AEA, OEA or PEA was quantified by LC–MS/MS. For kinetic analysis, β-Glyco-NAE concentrations were varied and initial rates were fitted to the Michaelis–Menten equation using nonlinear regression.

### Cellular NAE-feeding experiments

For cellular β-Glyco-NAE formation assays, HEK293T cells were seeded in 12 wells plate and treated with AEA, OEA or PEA in the presence or absence of GCase or GBA2 inhibitors. Inhibitors were added 24 hours before lipid feeding and maintained throughout the experiment. After 4 hours, cells were washed with ice-cold PBS and harvested for lipid extraction. β-Glyco-NAEs were quantified by LC–MS/MS and normalized to protein content or cell number.

### Endogenous β-Glyco-NAE formation in Neuro-2a cells

Neuro-2a cells were differentiated with retinoic acid (50 µM, final concentration) for 48 hours in low-serum medium (DMEM, 2% NCS). Differentiated cells were treated with vehicle or the FAAH inhibitor PF-04457845 at 1 μM for 48 h. Cells were washed with PBS, harvested and subjected to lipid extraction. Endogenous β-Glc- and β-Gal-NAEs were quantified by LC–MS/MS and normalized to protein content or cell number.

### Endogenous β-Glyco-NAE formation in RAW264.7 macrophages

RAW264.7 cells were treated with specific inhibitors of FAAH, NAAA, GCase and/or GBA2, either alone or in combination, for the indicated time points up to 48 h. Cells were washed with PBS and harvested for lipid extraction. β-Glyco-NAE levels were determined by LC–MS/MS and normalized to protein concentration or cell number.

### Mouse tissue collection and preparation

Mouse brain, heart, lung, liver, kidney, intestine, spleen, testes and pancreas were collected from C57BL/6J mice (8-14 weeks old). Mice were killed by cervical dislocation, according to guidelines approved by the ethical committee of Leiden University (license number AVD10600202215851). Organs were snap-frozen in liquid nitrogen immediately after dissection and stored at −80 °C. Frozen tissues were homogenized on ice in 20 mM HEPES buffer, pH 7.2, using 1.0-mm glass beads and a Bullet Blender or FastPrep homogenizer. Homogenates were cleared by centrifugation at 500 *g* for 1 min at 4 °C to remove beads, followed by two centrifugation steps at 2,500 *g* for 3 min at 4 °C to remove tissue debris. Protein concentration was determined using Bradford or BCA assay.

6-month-old FAAH-knockout male mice and their corresponding littermates (all in the C57BL/6J background, N= 4 per group)^86,87^ were housed at the animal facilities of Universidad Francisco de Vitoria (authorization number #281150000013). After sacrifice by cervical dislocation, brains were quickly removed, hippocampi and cortices dissected, and immediately snap-frozen and stored at −80 °C until further use.

For targeted lipidomics, homogenates were diluted to the appropriate protein concentration: 1 mg ml^−1^ for NAE analysis, 2 mg ml^−1^ for β-Glyco-NAE analysis and 5 mg ml^−1^ for glucosylceramide/glucosylated cholesterol analysis. Aliquots were snap-frozen and stored at −80 °C until lipid extraction.

### Human plasma and spleen samples

Control human plasma was obtained from Sanquin (Amsterdam, The Netherlands) and stored in aliquots at −20 °C until use. Plasma and spleen samples from patients with type 1 Gaucher disease were available through the Department of Medical Biochemistry, Leiden University, and had been collected with informed consent. Control spleen samples were obtained from age-matched healthy donors. All human samples were obtained and used in accordance with the Declaration of Helsinki and institutional ethical guidelines.

For spleen samples, tissue sections from multiple regions of the organ were pooled and homogenized in 150–250 μL KPi+ buffer containing 25 mM K₂HPO₄/KH₂PO₄, pH 6.5, and 0.1% (v/v)Triton X-100. Samples were homogenized with 1.0-mm glass beads using a FastPrep 5G homogenizer for two 40-s cycles at 6 m s^−1^. Protein concentration was determined by BCA assay and homogenates were diluted to 4 mg mL^−1^. Aliquots containing 100 μg total protein were stored at −20 °C until lipid extraction.

PD participants were recruited at the outpatient clinic for Parkinson’s Disease at the University Hospital of Tübingen. All participants were examined by a movement disorders specialist. Diagnosis of PD was defined according to UK Brain Bank Society Criteria. Genetic screening for pathogenic variants in the gene GBA1 were done as previously described.^79^ GBA1-subgroup classification of variant severity was based on established genotype risks reported for PD. Some variants that have been reported as nonrelevant for Gaucher disease have been proven to increase the risk for PD and therefore have been included in our analysis, for example, p.E326K and p.T369M. Plasma samples were collected and directly taken from the bedside and centrifuged within 60 minutes after collection and frozen at -80°C within 90 minutes after collection. The study was approved by the Ethics Committee of the University of Tuebingen (26/2007BO1, 404/2010BO1, 199/2011BO1, 702/2013BO1).

### CB1R and CB2R radioligand binding assays

Binding of parent NAEs and β-Glyco-NAEs to CB_1_R and CB_2_R was assessed using membrane preparations expressing human CB1R or CB2R and a radiolabeled cannabinoid ligand [^3^H]CP55,940 (1.5 nM) as previously described by Soethoudt *et al*.^88^ Membranes were incubated with radioligand and test compounds over the indicated concentration range in binding buffer. Non-specific binding was determined in the presence of excess unlabeled ligand. Bound and free radioligand were separated by rapid filtration, and radioactivity was quantified by scintillation counting. Data were normalized to total and non-specific binding and analyzed by nonlinear regression.

### TRPV1 activation assay

TRPV1 activation was measured in cells expressing human TRPV1 using a calcium-mobilization or fluorescence-based assay performed by Eurofins. Cells were loaded with calcium-sensitive dye and stimulated with AEA, β-Glc-AEA or control agonist. Fluorescence was recorded over time and responses were normalized to the positive control. Concentration–response curves were fitted by nonlinear regression where applicable.

### PPARα reporter assay

PPARα activity was measured using a luciferase reporter assay.^89^ Cells were transfected with a PPARα expression construct and a PPAR-responsive luciferase reporter, together with a normalization reporter where applicable. Cells were treated with PEA, β-Glc-PEA or control PPARα agonist for 24 hours at 37 °C. Luciferase activity was measured using a commercial luminescence assay and normalized to the internal control or protein content.

### Cytokine analysis in microglial cells

Microglial cells were seeded in in 96 wells plate (10.000 cells per well) and treated with LPS (1 ng/mL) in the presence or absence of β-Glyco-NAEs. After 24 hours, cytokine expression was measured measured by ELISA according to manufactures protocol (Invitrogen, TNF-α: ref 88-7324, IL-6: ref 88-7064).

### Protein determination

Protein concentrations were determined using Bradford (Bio-Rad) or BCA (Thermo Fisher Scientific) assays according to the manufacturer’s instructions. BSA was used as standard.

### Statistical analysis

Data are presented as mean ± SEM unless stated otherwise. Statistical analyses were performed using GraphPad Prism v9. Comparisons between two groups were performed using unpaired two-tailed Student’s t-tests or non-parametric alternatives where appropriate. Multiple comparisons were analyzed using one-way or two-way ANOVA followed by appropriate post hoc tests. Enzyme kinetic parameters were obtained by nonlinear regression using the Michaelis–Menten equation. P values <0.05 were considered statistically significant. Exact sample sizes, statistical tests and P values are provided in the figure legends or Source Data.

## Data availability

Source data for all figures are provided with this paper. Additional raw LC–MS/MS data and processed lipidomics tables are available from the corresponding authors upon reasonable request.

## Acknowledgements

This work was supported by Ministerio de Ciencia e Innovación – Agencia Estatal de Investigación and Fondo Europeo de Desarrollo Regional (PID2022-138461OB-I00; MICIU/AEI /10.13039/501100011033 and by FEDER, UE). This work is part of the Oncode Accelerator Project that has received funding from the Dutch National Growth Fund (NGF) under grant number NGFOP2201. Eurofins is kindly acknowledged for performing the TRPV1 assay with our compounds.

## Author contributions

M.v.d.S and J.M.F.G.A. conceptualized and supervised the study. A.F.S. developed the LC-MS/MS method, performed the lipidomics experiments, performed and cytokine assays. R.E.A.P. performed lipidomics experiments in human cells, plasma and spleen samples. B.G., I.T. and R.J.B.H. and N.v.d.B. synthesized the standards and internal standards of the glycosylated NAEs. T.v.d.W. setup the PPARα assay. M.J.F. helped with method development and measured the glycol-NAEs in PD samples. E.D. and T.V. performed the kinetics experiments. L.V.d.P., C.v.d.H. and L.H.H. setup and performed the CB binding assay. M.A, D.P. and H.S.O. provided chemical reagents and helped with data analysis. M.T.G. and J.R. provided the FAAH KO mice tissue and helped with their data analysis. C.D. K.B. and T.G. provided the PD plasma samples and helped with analysis. M.v.d.S., J.M.F.G.A and A.F.S. wrote the paper with input from all authors.

## Supplementary Information

**Fig. S1.**
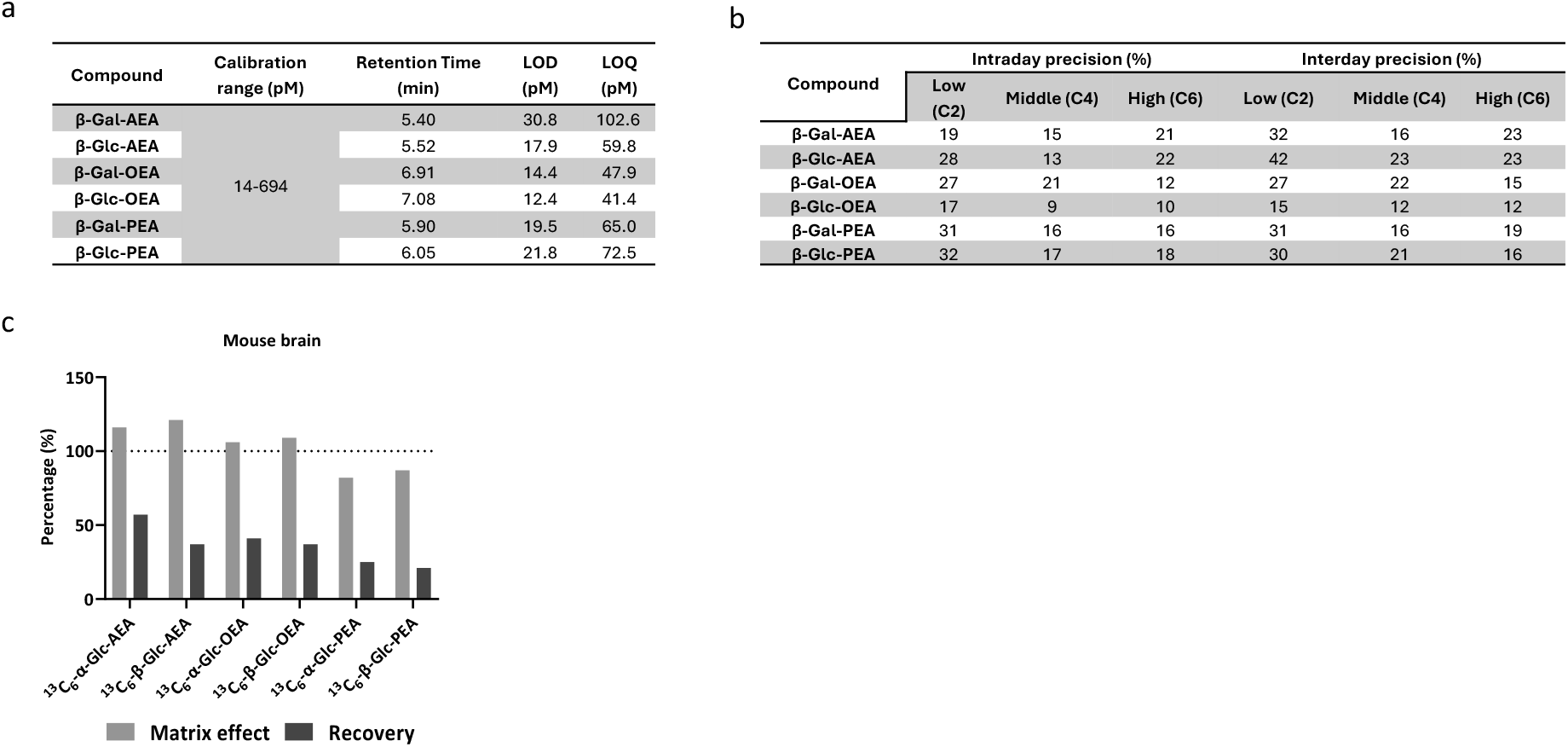
Development and validation of a targeted LC–MS/MS method for β-glyco-NAE quantification. **a,** Validation parameters including calibration range, retention time, limit of detection (LOD) and limit of quantification (LOQ). **b,** Intraday and interday precision RSD (%) for mouse brain. **c,** Matrix effect and recovery of internal standards determined for mouse brain.

**Fig. S2.**
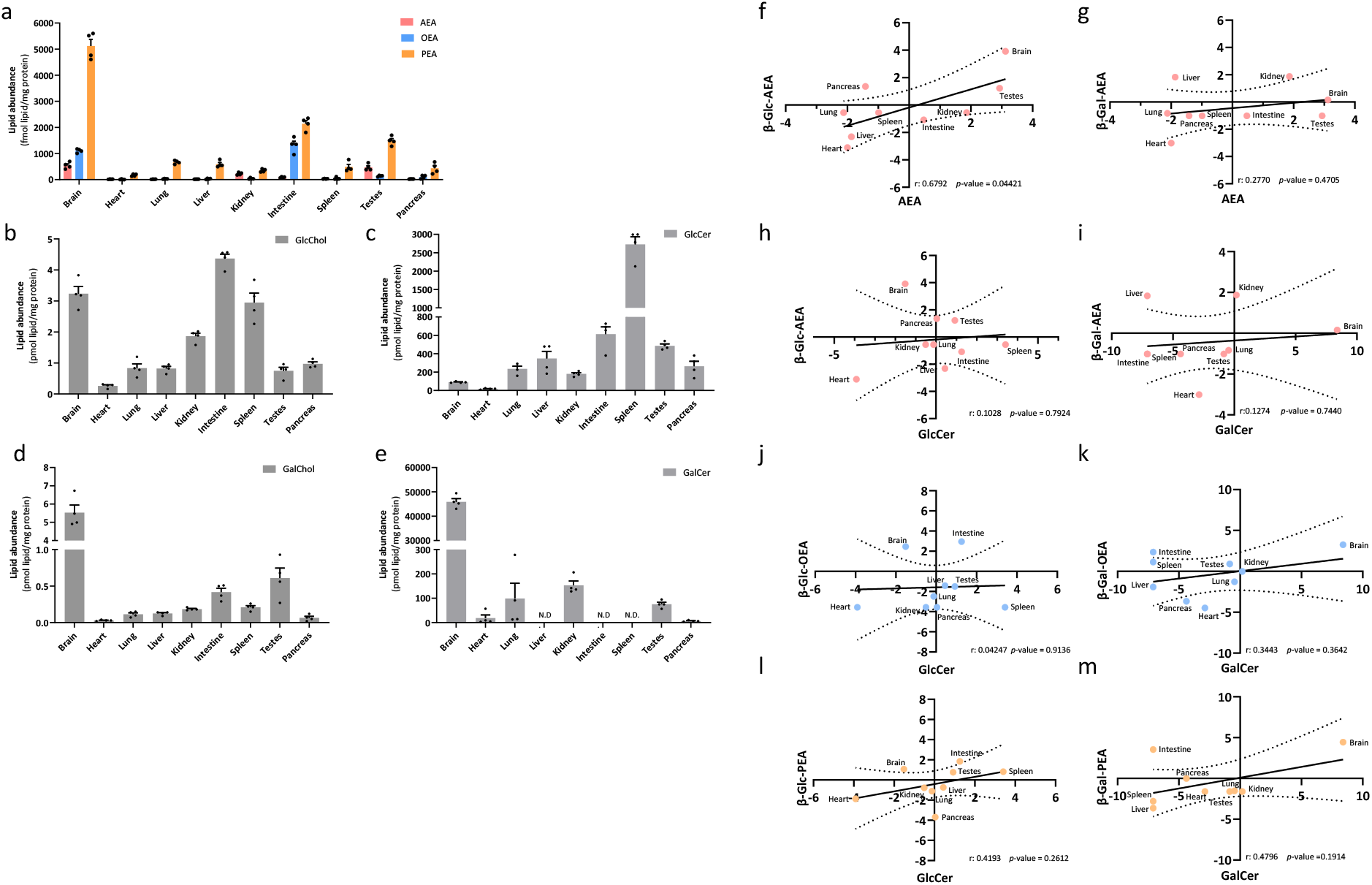
β-Glyco-NAEs are endogenous metabolites in mouse tissues and are regulated by NAE and glucosylceramidase metabolism in vivo. **a,** Tissue distribution of AEA, OEA and PEA, **b,c,** GlcChol and GlcCer, **d,e**, GalChol and GalCer in mouse brain, heart, lung, liver, kidney, intestine, spleen, testes and pancreas. Data is shown as mean ± SEM (N=4). **f,** Correlation plots of β-Glc-AEA and **g,** β-Gal-AEA versus their parent NAEs, **h,j,l** β-Glc-NAE versus GlcCer, **i,k,m,** β-Gal-NAE versus GalCer in mouse tissues. All data is represented as Log2 of the relative amount. A two-tailed Pearson correlation analysis was performed and the dotted lines represent 95% confidence interval of the best fitted line (solid line) as determined by linear regression analysis

